# Flashfm: A Flexible and Shared Information Fine-mapping Approach for Multiple Quantitative Traits

**DOI:** 10.1101/2021.04.09.439186

**Authors:** N Hernández, J Soenksen, P Newcombe, M Sandhu, I Barroso, C Wallace, J Asimit

## Abstract

Joint fine-mapping that leverages information between quantitative traits could improve accuracy and resolution over single-trait fine-mapping. Using summary statistics, flashfm (FLexible And SHared information Fine-Mapping) fine-maps signals for multiple traits, allowing for missing trait measurements and use of related individuals. In a Bayesian framework, prior model probabilities are formulated to favour model combinations that share causal variants to capitalise on information between traits. Simulation studies demonstrate that both approaches produce broadly equivalent results when traits have no shared causal variants. When traits share at least one causal variant, flashfm reduces the number of potential causal variants by 30% compared with single-trait fine-mapping. In a Ugandan cohort with 33 cardiometabolic traits, flashfm gave a 20% reduction in the total number of potential causal variants from single-trait fine-mapping. Flashfm is computationally efficient and can easily be deployed across publicly available summary statistics for signals in up to six traits.

## Introduction

Genome-wide association studies (GWAS) have been extremely successful in identifying genetic variants that are associated with a wide spectrum of diseases and related traits^1^. Among these findings are many examples of pleiotropy, where a gene affects several phenotypes^2^. This could be due to a shared variant affecting a pathway involved in multiple related phenotypes. Identifying the causal mutations which underlie such findings is key to facilitating translation into new therapeutic targets or elucidating new biological insights. However, this is a complex task. Lead SNPs (those with the most significant p-value) are often correlated (are in high linkage disequilibrium - LD) with many other variants in the genome. In addition, lead variants are not necessarily causal and may be detected due to LD with the causal SNP(s). Statistical fine-mapping is therefore needed to refine sets of potential causal variants. The fewer the number of variants identified as potentially causal, the easier it will be to perform downstream functional validation experiments. Here, we focus on three fine-mapping challenges: multiple trait analysis, missing trait values, and related individuals in a cohort.

Bayesian approaches are common in fine-mapping, and use a Bayes’ factor (BF) to summarise the evidence of association, either for each SNP under the assumption of a single causal variant, or for each combination of SNPs in the more flexible multiple causal variant setting. In the more general multiple causal variant setting, the BF for a model compares the evidence for a model consisting of a particular set of SNPs that could be causal for a trait to the null model of no causal SNPs. These BFs can be calculated for different combinations of causal SNPs and, for a specified prior probability, posterior probabilities (PP) for each causal variant model are easily calculated, modelling the pattern of association within a region^3,4,5^. Models may then be prioritised by PPs.

Current fine-mapping methods do not allow missing trait data, so that a portion of the data is disregarded to avoid any missing measurements. Also, fine-mapping methods that make use of GWAS summary statistics (e.g. JAM^4^, FINEMAP^5^) assume the specified sample size *N* relates to independent individuals, whilst the effective sample size after adjustment for relatedness via a linear mixed model, is <*N*. Such inflated *N* will suggest more certainty than actually present in the sample.

Jointly fine-mapping multiple traits could give an improvement in fine-mapping accuracy and resolution, analogous to the power increase for multi-trait GWAS, but this is computationally challenging due to the many possible combinations of models (allowing multiple causal variants) between traits, which is not an issue for multi-trait GWAS that involve testing only one SNP for association with multiple traits. For this reason, few methods exist for fine-mapping with more than two traits.

When multiple traits have signals in the same region, colocalization is often used to evaluate how likely the traits share a causal variant. In some methods colocalization includes the fine-mapping step of identifying potential shared causal variants, under the simplifying assumption of at most one causal variant for each trait^6^. An approach that allows multiple causal variants, eCAVIAR^7^, requires that the traits are measured from independent studies. This is because eCAVIAR is designed to assess if there are shared causal variants between a quantitative trait from a GWAS and expression quantitative trait loci (eQTL), which are often available in independent studies.

One approach for jointly fine-mapping signals for quantitative traits from the same study is to limit the combinations of models by assuming all causal variants are shared between traits and allowing heterogeneity in effects, as in fastPAINTOR^8^. Although this greatly reduces the number of models to consider, this does not allow traits to have different sets of causal variants. Although multi-trait fine-mapping is motivated by traits having shared causal variants, the possibility of a single shared causal variant and additional trait-specific causal variants cannot be ignored, as well as the fact that causal variants may actually differ for all traits. An approach that does not make this assumption is (Multinomial Fine-mapping, MFM^9^) which can be used to fine-map multiple related diseases with shared controls. This approach is made computationally tractable by showing that the joint Bayes’ factor (BF) for *M* diseases with shared controls is a function of the individual disease BFs, model complexity and sample sizes.

We introduce a method for general quantitative multi-trait fine-mapping that solves the issues outlined above, allowing for related individuals and missing trait measurements: flashfm (flexible and shared information fine-mapping). Flashfm uses a similar Bayesian framework (same prior probabilities) to MFM^9^. However, flashfm addresses different statistical challenges than MFM, as we now have multiple quantitative traits measured on the same individuals, so we need to account for correlation between the traits. In addition, the statistical modelling differs between the methods, as MFM uses a multinomial logistic framework, whereas flashfm is in a multivariate regression framework.

Through extensive simulations we demonstrate that flashfm improves fine-mapping accuracy and resolution over single-trait fine-mapping. Specifically, it provides higher support for the correct model (mean PP increase of 0.15). Also, when 10-50% of the data were missing for one of the traits, flashfm refined the group of causal variants from single-trait fine-mapping, with a median percentage reduction of the SNP group size ranging from 31% to 11%.

We subsequently applied flashfm to genetic association signals for 33 cardiometabolic traits measured in 6407 participants from Uganda - the largest GWAS of a single population from Africa^10,11^; within this sample 2907/6407 are at least second degree related. Understanding the underlying genetic contributions to cardiometabolic traits is important due to the growing global burden of disability and death attributed to cardiometabolic disorders such as hypertension, coronary artery disease and type 2 diabetes, which are predicted to become greater in developing countries than that of infectious diseases (e.g. HIV/AIDS)^12^. In this dataset, we see several cases where flashfm improves on single-trait fine-mapping, distinguishing between two models with similar levels of support under single-trait fine-mapping and generally producing higher resolution solutions. In particular, the groups of likely causal variants constructed under flashfm are subsets of those from independent fine-mapping, resulting in finer resolution.

## Results

### Flashfm - Conceptual framework

Flashfm uses GWAS summary statistics to jointly fine-map genetic associations for multiple quantitative traits that have sample overlap, and allows flexibility for missing measurements and for related individuals. The GWAS for each trait could either be from a single cohort or from a meta-analysis of multiple cohorts, where traits overlap between cohorts, though may not be measured in all cohorts.

As the traits are measured on the same individuals, it requires the trait covariance matrix, which may be calculated from an in-sample study or approximated from the GWAS and trait summary statistics^13^. It uses GWAS summary statistics from each trait to fit the joint models, allowing for multiple causal variants for each trait, with no restrictions on shared causal variants between traits. Flashfm shares information between traits by up-weighting joint models (combinations of fine-mapping models across traits) that have a shared causal variant. For each trait, flashfm outputs the top SNP models and the model posterior probability (PP), adjusted for information from the other traits. It also provides the marginal posterior probability (MPP) that a SNP contributes to any model.

In single-trait fine-mapping, models are prioritised by posterior probabilities (PP) which are calculated from the pre-specified prior probability and the BF that is calculated from the data. For multi-trait fine-mapping, we first find the PP for joint models then marginalise the joint PPs to get trait-specific PPs that are adjusted for the other traits; when there are two traits, the multi-trait adjusted PP for model *i* of trait 1, 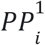, is 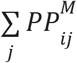, where 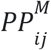 is the joint PP of the joint model consisting of the configuration of model *i* for trait 1 and model *j* for trait 2. In general, for any number of traits, the trait-adjusted PP for a particular model is found by summing over the PPs of all joint model configurations that contain model *i* for trait 1.

To generate posterior support for fine-mapping models, flashfm needs to calculate the ABF (approximate Bayes’ factor) for all possible model combinations across SNPs and traits. To make this computationally feasible, we derive an expression for the joint ABF for *M* traits, denoted ABF^M^

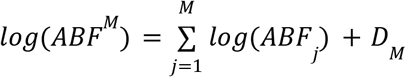

where *D_M_* depends on terms that depend on the GWAS summary statistics, covariance matrix of the traits, and sample sizes; either a genotype matrix or both a SNP covariance matrix and reference allele frequencies (RAFs) are needed from a reference panel or in-study sample (Supplementary Information, Section 1.1). When there is missing data, *D_M_* includes additional terms that account for the individuals that do not have measurements for all traits, using a combinatorial argument. The joint BF is first expressed as the BF for the multiple traits (at a particular model for each trait) on the portion of the sample that have no missing data, then an additional BF is added for each combination of traits with data available, careful to include each individual in only one term (Supplementary Information, Section 1.2).

The prior probability for the joint models includes a term κ that gives more weight to joint models that have a shared causal variant between the traits. For models M_*i*_ (trait 1) and *M_j_* (trait 2), with marginal prior probabilities *p_i_* and *p_j_,* we denote the joint model as configuration C_ij_ and set the prior probability as

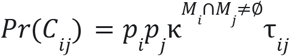

such that the joint prior is simply the product of the marginal priors when there is no overlap of variants between models, and is otherwise upweighted. The term κ is derived in a combinatorial manner and is identical to that used in MFM^9.^ It requires setting a target odds (TO) of the odds for traits not having a shared causal variant compared to having a shared causal variant; setting TO=1 coincides with a 50/50 chance of shared causal variant(s) and is the setting that we use. When κ = 1, there is no weight for joint models with shared causal variants and the flashfm PP for each model for a given trait is the same as that from single-trait fine-mapping, which we also refer to as independent fine-mapping, as it does not make use of data from other traits. The term τ_*ij*_ is a correction factor that anchors the prior probabilities so that the prior probability of traits having particular model sizes is consistent for different values of κ; identical to MFM^9^.

This means that rather than calculating the joint BF for each model, the joint PP could be directly calculated from the model PPs from single-trait fine-mapping; e.g. the model PPs from the *.config file of FINEMAP^5^ could be input to flashfm. In the case of two traits, the trait-adjusted PP for model γ_*i*_ of trait 1 is calculated using

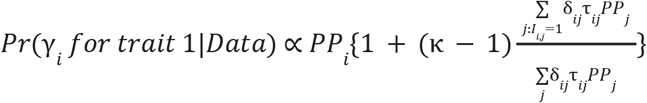

where δ_*ij*_ = *exp*(*D_ij_*) is the value of *D_M_* for *M*=*2* traits at models *i* for trait 1 and *j* for trait 2 (Supplementary Information, Section 1.3). This gains efficiency by making use of previously generated single-trait fine-mapping results. If single-trait fine-mapping results are not available, flashfm includes a function to run an expanded version of JAM^4^ (Methods). So, flashfm requires the same information needed for single-trait fine-mapping, as well as the trait covariance matrix and results from single-trait fine-mapping.

In flashfm, relatedness is accounted for by making use of GWAS summary statistics from a linear mixed model method such as GEMMA^14^ or BOLT-LMM^15^ and using the summary statistics to approximate the effective sample size *N_e_* for each trait - *N_e_* is approximated at each SNP and the median of the *N_e_* is taken as the effective sample size for the trait; the Neff function in flashfm does this calculation.

The final results of interest are the top models for each trait, adjusted for the other traits, rather than the PP for a joint model. This means the results can be used comparably to those from single trait fine mapping, but with an expectation of greater accuracy because of the leverage of information from the other traits.

Even borrowing information across traits does not remove the complication of LD. Two or more SNPs in high LD may provide equivalent statistical information to explain any trait. Therefore, in addition to single SNP posterior probabilities, flashfm output also expresses SNP models in terms of groups of such nearly equivalent SNPs, constructed so that SNPs in the same group are in LD and rarely appear together in a model. These SNP groups could be viewed in a similar way to the credible sets that are constructed for single-trait fine-mapping. As flashfm leverages information from the other traits, the per SNP PPs tend to concentrate on fewer SNPs, so the resulting SNP groups tend to be smaller than those from independent fine-mapping.

### Flashfm has higher accuracy and resolution than independent fine-mapping

Extensive simulations of two quantitative traits with varying sample size, proportion of missing trait measurements, and trait correlation suggest the same general conclusion. Flashfm leads to improvements over independent fine-mapping in terms of higher accuracy, as flashfm gives higher levels of evidence (larger posterior probability) for the correct model (Figure 1), and finer resolution, as indicated by smaller SNP groups that are constructed based on the PPs from flashfm.

**FIGURE 1.**
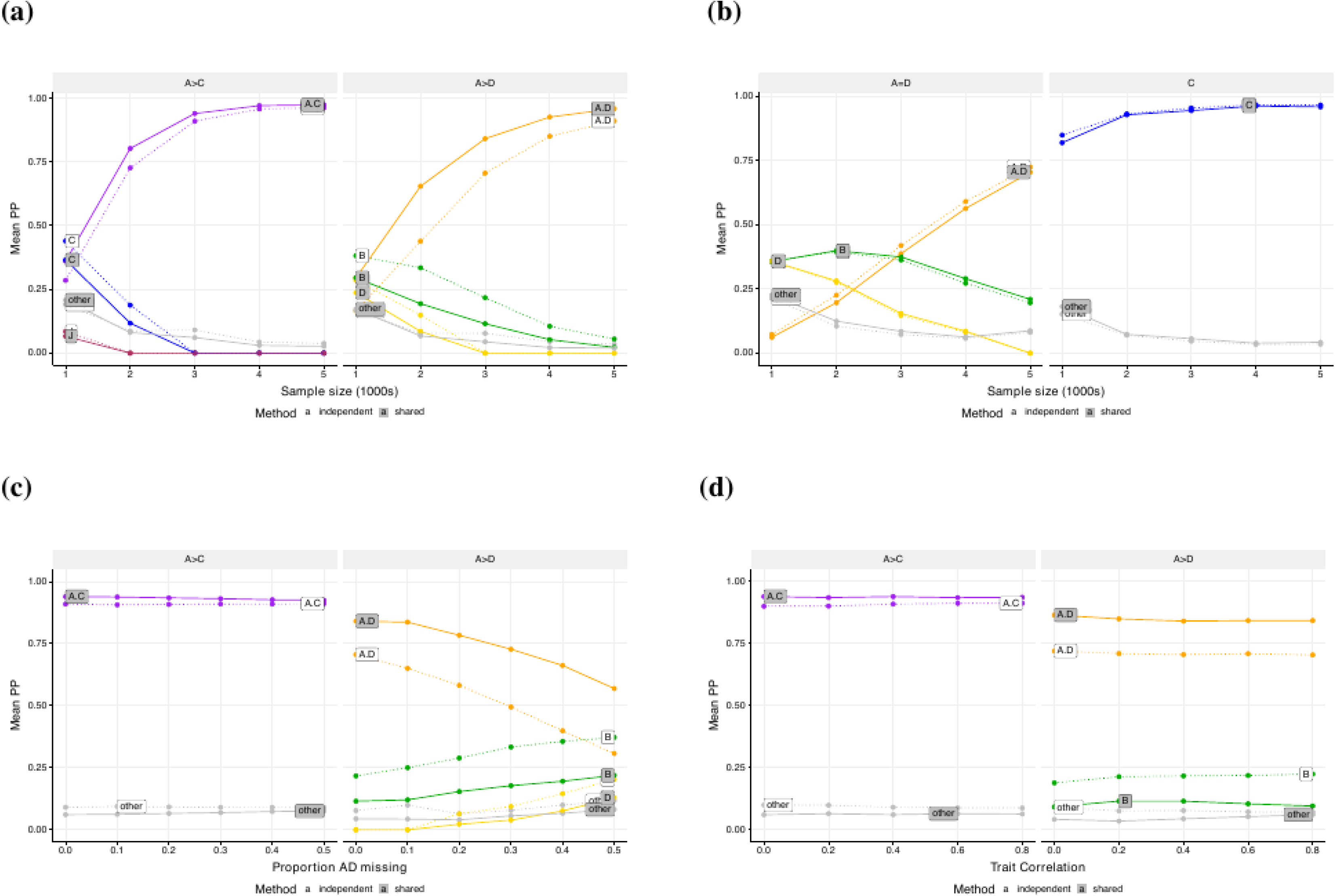
Comparison of fine-mapping from flashfm and single-trait analyses. When traits share a causal variant, flashfm has higher accuracy than single-trait finemapping, regardless of amount of missing data and trait correlation; both methods have similar accuracy when there are no shared causal variants. Causal variants were simulated for two traits with models defined by SNP groups from the *IL2RA* region. In **(a)**, **(b)** and **(d)** Trait 1 has causal variants A+C, while trait 2 has A+D causal variants, both with the same effect size: β_*A*_ = *log*(1. 4) and β_*D*_ = *log*(1. 25). In **(a)** and **(c)** There are no missing data and the sample size varies from 1000 to 5000. In **(b)** the sample size is fixed at 3000 and the proportion of missing data for trait A+D varies from 0 to 0.5. In **(d)** the sample size is fixed as 3000 and the correlation between traits varies. In **(c)** Trait 1 has causal variants A+D with β β_*A*_ *log*(1. 25) and β_*D*_ = *log*(1. 25), while trait 2 has a single causal variant C with β_*C*_ = *log*(1. 25). Source data are provided in Supplementary Data 1 Tables 1-5.

**Table 1:** When traits share a causal variant, flashfm tends to yield smaller SNP groups than those from single-trait fine-mapping, regardless of amount of missing data and trait correlation; both methods have similar resolution and accuracy when there are no shared causal variants. In (a), (c), and (d) SNP A is a causal variant with β=log(1.4) for both traits 1 and 2; trait 1 has a second causal variant D and trait 2 has second causal variant C, both with β=log(1.25). **(a)** Traits 1 and 2 have correlation 0.4 and were both measured on all individuals in a sample of size *N*; **(b)** Trait 1 has causal variants A (β=log(1.25)) and D (β=log(1.25)) and trait 2 has causal variant C (β=log(1.25)). **(c)** Traits 1 and 2 have correlation 0.4, trait 1 has missing measurements for proportion *p_1_* of individuals in a sample of size *N* and trait 2 has no missing measurements. **(d)** Traits 1 and 2 have specified correlation Cor(Y_1_,Y_2_) and were both measured on all individuals in a sample of size 3000.

| Trait 1: A + D |  |  |  | Trait 1: A + D |  |  |
| --- | --- | --- | --- | --- | --- | --- |
| Trait 2: A + C |  |  |  | Trait 2: C |  |  |
| N | (a) Median percentage size reduction | Single-trait group A coverage | Multi-trait group A coverage | (b) Median percentage size reduction | Single-trait group A coverage | Multi-trait group A coverage |
| 1000 | 0 | 1 | 0.986 | 0 | 0.944 | 0.944 |
| 2000 | 10.5 | 0.997 | 0.99 | 0 | 0.98 | 0.97 |
| 3000 | 28.5 | 1 | 1 | 0 | 1 | 0.99 |
| 4000 | 32.5 | 1 | 0.997 | 0 | 0.988 | 0.984 |
| 5000 | 33.3 | 1 | 1 | 0 | 0.996 | 0.996 |
| (c) Trait 1 proportion missing, $p_1$ | Median percentage size reduction | (d) $\text{Cor}(Y_1, Y_2)$ | Median flashfm group A size decrease | | | |
| 0 | 28.5 | 0 | 33.3 |  |  |  |
| 0.1 | 31.4 | 0.2 | 33.3 |  |  |  |
| 0.2 | 27.8 | 0.4 | 28.5 |  |  |  |
| 0.3 | 18.8 | 0.6 | 21.1 |  |  |  |
| 0.4 | 16.7 | 0.8 | 14.3 |  |  |  |
| 0.5 | 10.5 |  |  |  |  |  |

As described in the Methods, according to the specified simulation model, a causal variant was selected from the SNPs A_1_, C_1_, and D_1_, such that one trait had two causal variants A_1_+D_1_, and the other trait either had two causal variants A_1_+C_1_, one of which was common, or a single distinct causal variant C_1_ (Supplementary Data 1.1). These SNPs were selected as they are known to represent an example of LD with joint tagging; a different SNP “B” jointly tags A_1_ and D_1_, such that when A_1_ and D_1_ are causal variants, B is often chosen by fine mapping analyses^9^. The data were simulated across 345 SNPs, including many in LD with A_1_, C_1_, and D_1_ (see Methods for details). We label the groups constructed by each fine-mapping approach such that a group that contains A_1_ is labelled A, and if no group contains A_1_, but there is a group with a SNP that has r^2^> 0.7, then this group is labelled A; likewise for the labelling of groups B, C, D, and J. In all settings, for groups C and D, there were negligible differences in group sizes between those constructed from flashfm and those based on independent fine-mapping; groups C and D were not shared between traits so do not gain from shared information. For all settings there is a negligible difference in probability that the SNP group contains the causal variant used in the simulation (i.e. similar coverage) between single and multi-trait fine-mapping. We provide detailed resolution results for group A (Table 1). From this point, the A+D trait refers to a trait simulated to have causal variants A_1_+D_1_.

When there is no shared causal variant (trait 1 has causal variants A_1_+D_1_ and trait 2 has causal variant C_1_), there are negligible differences between flashfm and independent fine-mapping; the model PPs are nearly indistinguishable (Figure 1(b); Supplementary Data 1.2) and for each sample size the median difference in group A sizes between the methods is zero, with similar coverage (Table 1(b)). Thus, application of joint fine mapping where variants are not shared does not incur a penalty.

When both traits have two causal variants, one of which is shared, flashfm tends to show higher support than independent fine-mapping for the true model (mean PP increase of 0.17 with a sample size of at least 3000; Figure 1(a), Supplementary Figure 1, Supplementary Data 1.3-1.4). In low sample sizes independent fine-mapping of the A+D trait either prefers B or has little difference between A+D and B, then as sample size increases to 3000 there is a clear switch to the correct model, A+D. This switch occurs faster for flashfm (at N=2000) due to the borrowed information between the traits. Also, flashfm tends to result in fewer SNPs in group A with a median size reduction of 32% for N ≥ 3000 (Table 1(a), Supplementary Data 1.5).

For a sample of *N*=3000 and varying proportions (0 to 0.5) of missing data from the trait with causal variants A+D, flashfm showed an average gain of 0.20 in the level of PP support for A+D over that of independent fine-mapping (Figure 1(c); Supplementary Data 1.6). When half the data were missing for trait A+D independent fine-mapping no longer had a preference for the true model. There is negligible difference between mean PP for the two methods for the trait with A+C causal variants, regardless of the proportion of missing data for the A+D trait. Although flashfm has finer resolution for each degree of missingness, the improvement is highest for lower proportions of missingness (median reduction in group size of 23%) (Table 1(c)).

By varying the correlation (0 to 0.8) between two traits (A+D and A+C) we found that the PP remained similar within each method and for trait A+D flashfm gave a median PP increase of 0.14 over independent fine-mapping (Figure 1(d); Supplementary Data 1.7). Compared to independent fine-mapping, flashfm reduces the median group size 28.5%, with the greatest reduction at lower levels of correlation (33.3% reduction when the trait correlation is 0.2 or 0; Table 1(d)). Low/moderate correlation likely gives higher gains from sharing information between traits because there is more for a trait to “learn” from the other trait; as the traits are measured on the same cohort, highly correlated traits will have similar information to each other so not as much as a gain compared to lower correlations.

### Application to cardiometabolic traits in a Ugandan cohort

We considered genetic association signals from 33 cardio-metabolic traits in a Ugandan cohort of 6407 individuals with 45% relatedness of at least second degree^10^ (Supplementary Data 1.8 and 1.9). Based on association signals (p<1E-6) with at least 2 of the traits, we constructed 56 regions for fine-mapping (Supplementary Data 1.10 and 1.11; Supplementary Figure 2).

Within the 56 regions the total number of potential causal variants from the top models of FINEMAP was 1147, whereas flashfm reduced this total by 20% to 914 variants (Supplementary Data 2). Among these regions, 52 of them (93%) indicated improved results of flashfm over FINEMAP^5^ in terms of either increased PP of the top model, finer resolution of the SNP groups, or both. The median PP increase of flashfm over FINEMAP was 15% (mean 24%) for these regions. The remaining 4/56 regions had concordant SNP groups between the two methods, but flashfm had a slightly lower PP than FINEMAP (median decrease 0.05%). In 36% of the regions (20/56), flashfm gave a finer resolution than FINEMAP^5^, based on the total number of SNPs in the SNP groups of the top model (for each trait) from each method; the median reduction in the total number of SNPs for each trait was 31% with a maximum of 91%. In 88% of the regions (49/56), flashfm had higher confidence than FINEMAP for the top model with a median PP increase of 18% (mean increase 29%); in half of the regions (28/56) flashfm had a noticeable gain with an average PP increase of 52%.

We highlight 5 regions that are improved for both higher confidence in model selection and notable reduction of potential causal variants in the top model of at least one trait (Table 2), and also provide functional annotations for these SNPs, based on HaploReg 4.1^16^ and VEP^17^ (Supplementary Data 1.12).

**Table 2.**
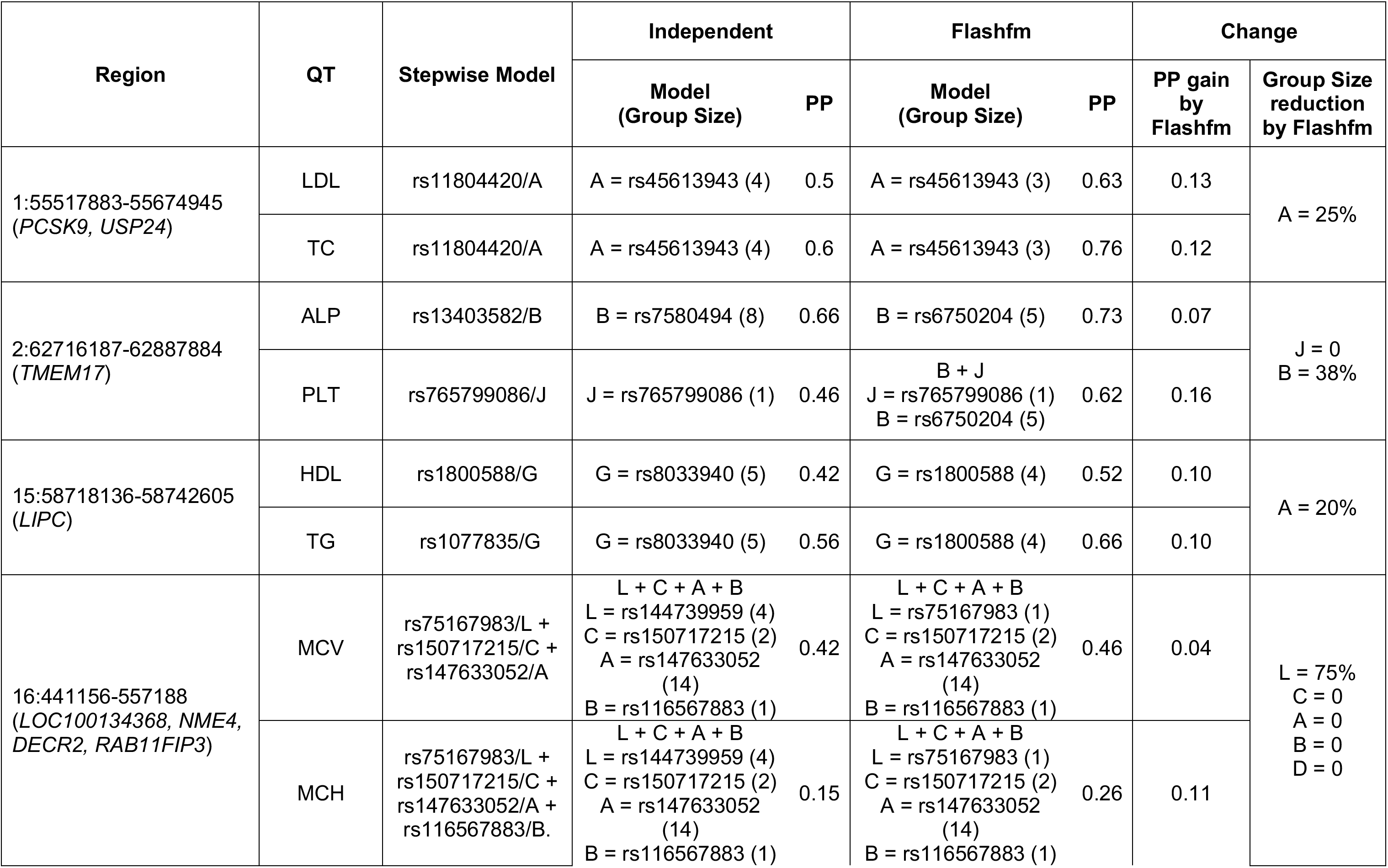
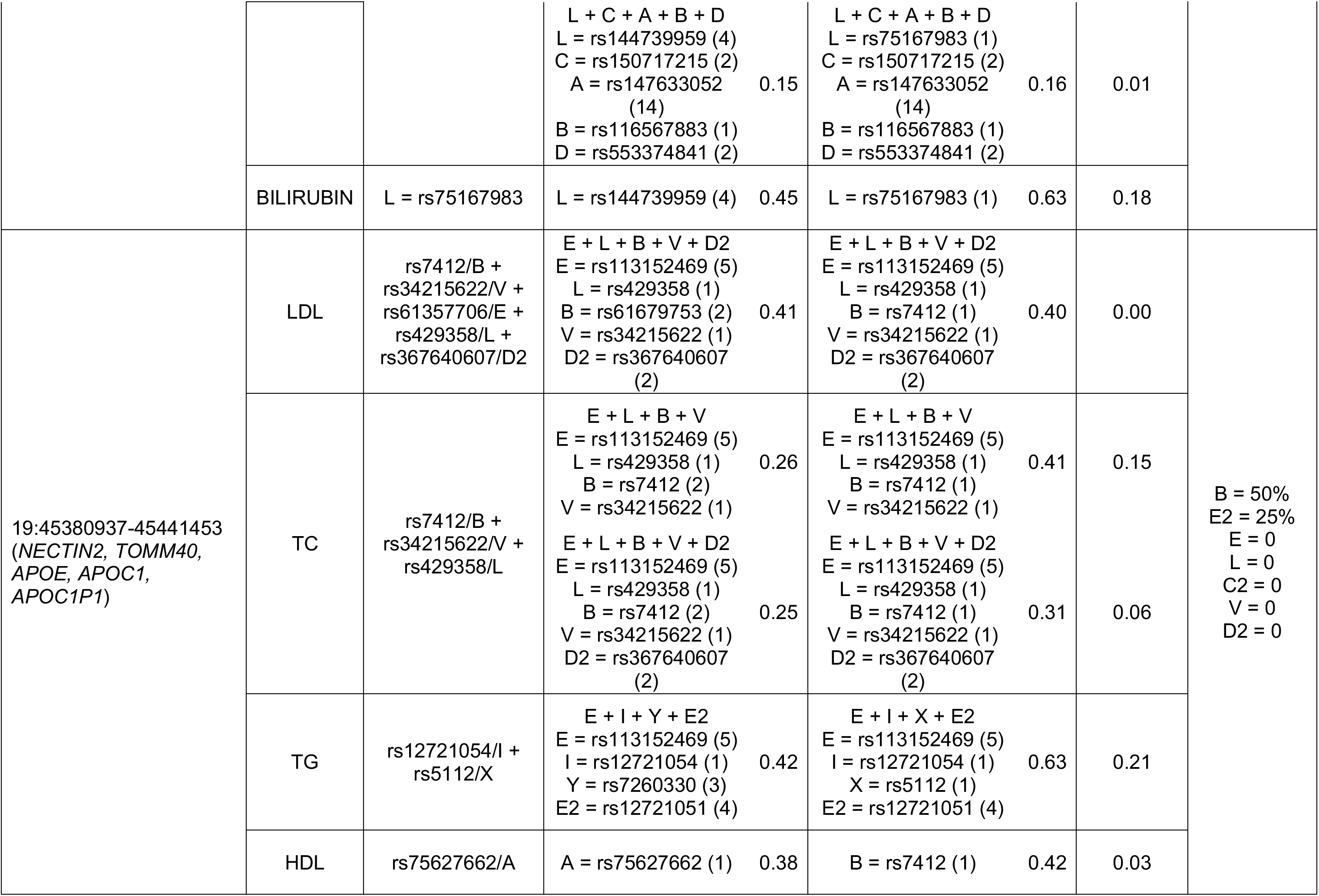
Regions with top models chosen by stepwise (SW), Independent and Flashfm where there is a noticeable reduction in SNP group sizes and/or PP of top model. Each row summarises results for a single region, defined by chromosome, start and end base-pair position and nearby gene(s). Each cell lists the SNP groups in a model; model A+B indicates all 2-SNP models with one SNP from group A and one SNP from group B. The number of SNPs in each group is given in brackets below each model. Here we list a representative SNP from each group; rs IDs are from build 37/hg19. The SNPs belonging to each group and their functional annotations are given in Supplementary Data 1.12.

Of particular interest is region 19:45380937-45441453 (*PVLR2, TOMM40, APOE, APOC1, APOC1P1*), which shows association to multiple lipid traits: LDL, TC, TG and HDL (Figure 2). This example highlights the utility of borrowing information between related traits when fine mapping weaker signals. Previous studies have concluded that the LDL association in *APOE* is explained by the missense variants rs7412 and rs429358, which together define *APOE* ε-alleles^18,19,20^. Both of these SNPs appear in top LDL and TC models by FINEMAP and flashfm. These two methods identify rs429358 as the only SNP in group L in models for LDL and TC. Flashfm isolates rs7412 as a single SNP in group B, whereas FINEMAP includes an additional SNP in group B, demonstrating that flashfm refines the SNP group of FINEMAP to the correct causal variant.

HDL shows a much weaker association than the other traits (min p ~ 10^−9^), and FINEMAP selects its top SNP rs75627662 as its best candidate. The second strongest SNP in the region for HDL is rs7412 (r^2^ with rs75627662 is 0.56) and flashfm levers information on LDL and TC to select this SNP as its best candidate for HDL. rs75627662 is an intergenic SNP upstream of *APOE*, while rs7412 encodes a missense variant (Arg158Cys, which is also known as the *APOE* e2 allele). Homozygous carriers for this variant have the condition Type III hyperlipoproteinemia^21^ (OMIM 617347), which is characterised by hyperlipidaemia and affected individuals are susceptible to severe coronary artery disease. ApoE is a 34 kDa glycoprotein, initially noted as a component of plasma VLDL and HDL. rs7412 is known to have an effect on the efficacy of Atorvastatin, a statin medication used to prevent cardiovascular disease in those at high risk (ClinVar^22^, accession RCV000211178.1); Atorvastatin decreases LDL and triglycerides in the blood and increases HDL. These observations highlight the complex interrelation among LDL, HDL and triglycerides and their regulation and the value of using approaches that capture information from multiple traits.

**FIGURE 2.**
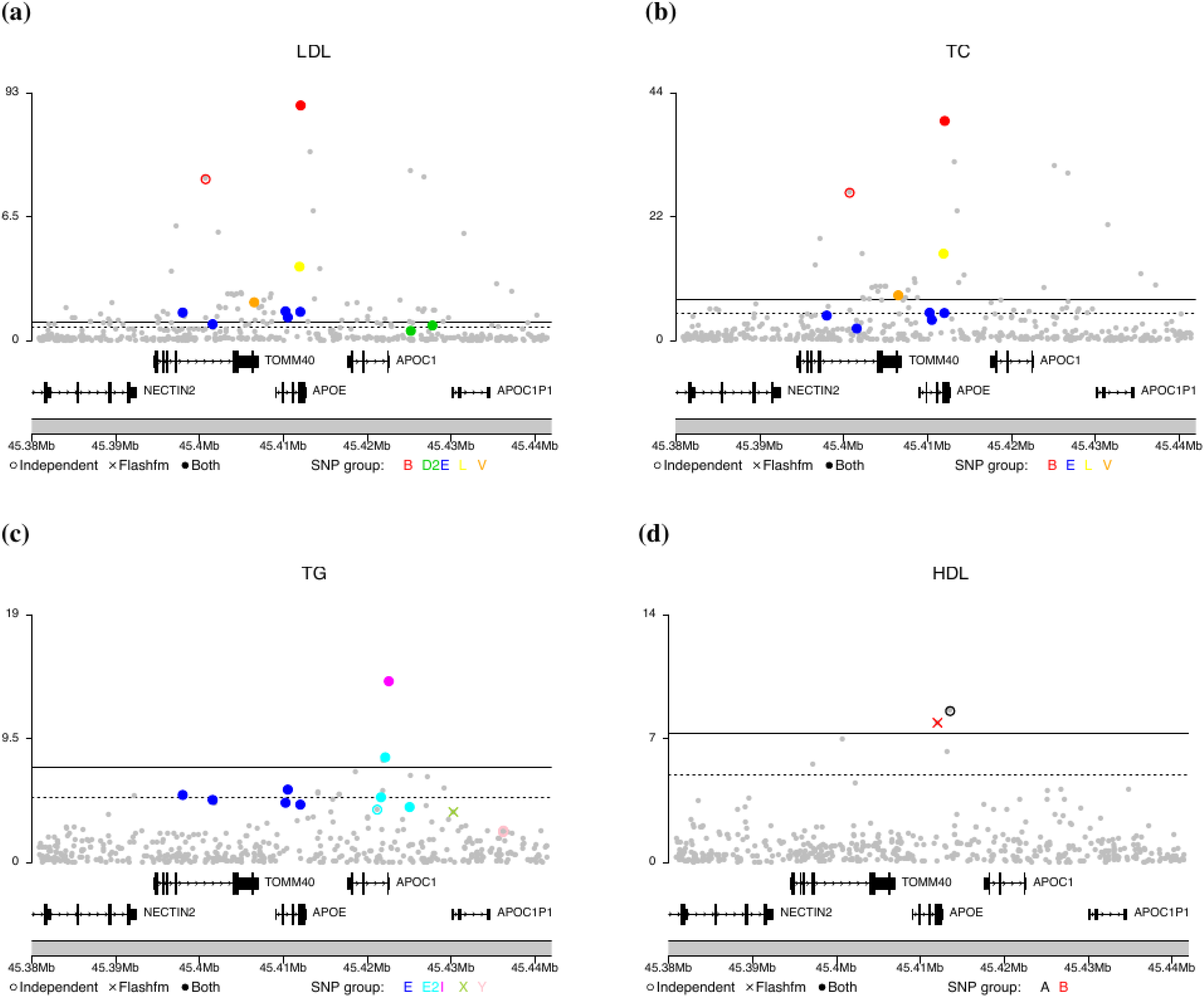
Fine-mapping of signals for four lipid traits in region 19:45380937-45441453. The −log_10_p for SNPs in the top SNP groups for (a) LDL; (b) total cholesterol (TC); (c) triglycerides (TG); (d) HDL are shown for both FINEMAP and flashfm. The two methods agree on a 5-SNP model for LDL (a) and a 4-SNP model for TC (b). The top model for TG (c) has 4 SNPs under both methods but differ in one SNP group; FINEMAP prefers 3-SNP group Y (very near one another so appear as one) and flashfm selected single SNP group X (mean r^2^ of SNPs in Y with X is 0.315). For HDL (d), a different single-SNP model was selected by the two methods; FINEMAP favoured group A, whereas flashfm selected group B. The solid coloured circles show SNPs that belong to the SNP groups constructed by both methods, the empty coloured circles represent SNPs that are only in the FINEMAP SNP group;; solid grey circles show all other SNPs in the region. In (c) and (d) an X represents a SNP that appeared in a top model for flashfm and not FINEMAP and empty circles indicate SNPs that appeared in top models for FINEMAP and not flashfm. Position is given according to hg19/build 37. Some of the genes in this region include *APOE, APOC1* and *TOMM40.*

## Discussion

Simultaneous fine-mapping of multiple traits helps in understanding pleiotropic associations by identifying sets of shared potential causal variants that underlie multi-trait associations in the same locus. Jointly fine-mapping traits with flashfm leads to improvements in both accuracy and precision when there are shared causal variants between traits, and leads to similar results to independent fine-mapping when there is no such sharing among traits. Importantly, the flashfm SNP groups are typically a subset of the FINEMAP SNP groups, suggesting that flashfm leverages the information between traits to refine the sets of potential causal variants. This approach only requires GWAS summary statistics, an estimate of the trait covariance matrix, either a genotype reference panel or the covariance matrix and MAFs from a reference panel, and single-trait fine-mapping results (PP for each model). GWAS summary statistics could be used to approximate the effective sample size for each trait. The input of flashfm makes it readily applicable to meta-analysis results from multiple traits, and the large sample sizes do not increase computational time as only summary-level data is needed.

Flashfm is flexible to accept output from any single-trait fine-mapping approach and, for convenience, we include an expanded version of JAM^4^ to obtain single-trait fine-mapping results in the flashfm R package. When using flashfm for the Uganda GWAS of cardiometabolic traits, we used single-trait fine-mapping results from both expanded JAM and FINEMAP^5^, which generally resulted in concordant results.

Our simulation studies demonstrated that flashfm tends to give greater improvement in resolution when the traits have a common causal variant and a moderate/low correlation between them, though even with highly correlated traits, there is some resolution gain over single-trait fine-mapping. Likewise, the greatest refinement in resolution within the Uganda data tended to be for sets of traits that included at least one trait with a correlation below 0.9 and single-trait fine-mapping top trait models with common SNP groups, which were refined by flashfm. Replication in another data set and downstream functional validation experiments are needed to confirm that the SNPs excluded by flashfm are indeed unlikely to be causal variants. These results promote the joint fine-mapping of traits that are relevant to certain disorders (and have genetic association in the same regions), but not usually considered together; flashfm could not only refine the sets of potential causal variants for the traits, but also reveal new shared causal variants between them.

There are clear advantages to multi-trait fine-mapping with flashfm, and the inclusion of diverse ancestries could lead to further improvements in resolution; where ancestries have common signals, the differences in LD between the ancestries could help pinpoint the causal variant(s)^23^. A multi-ancestry version of flashfm is in progress as a crucial extension to this software, that will allow for multiple reference panels from the different ancestries.

## Methods

### Expanded JAM

The Joint analysis of marginal summary statistics (JAM)^4^ facilitates fine-mapping from marginal summary statistics. JAM requires a thinned reference panel such that the genotype matrix does not contain SNPs in high LD and then infers joint LD-adjusted multi-SNP models, highlighting the best SNPs, and combinations of SNPs, via a Bayesian sparse regression framework. Included in the flashfm software, we developed an expanded version of JAM, JAMexpanded.multi (https://github.com/jennasimit/flashfm/blob/master/R/prep.R), where each of these joint multi-SNP models with PP > 0.005 is expanded by considering all the possible models formed by all the combinations of SNPs in the JAM model, in the same manner as the fine-mapping approach GUESSFM^24^ (see Supplementary Material for further details). Then, each of the expanded models is evaluated individually via approximate bayes factors.

### SNP Groups

Rather than reporting results in terms of SNPs, we construct SNP groups using an algorithm based on the group.multi function in GUESSFM (https://github.com/chr1swallace/GUESSFM/blob/master/R/groups.R). SNPs with marginal posterior probability of inclusion > 0.001 were grouped such that SNPs in the same group are in LD - high *r*^2^ - and rarely selected together in models (model selection correlation (*r*_model_) should be negative); both *r*_model_ and *r*^2^ are used so that our SNP grouping is informed by both model posteriors and LD. This algorithm was used in MFM^9^ to construct SNP groups based on the single-disease fine-mapping posterior probabilities and these same groups were used to summarise the fine-mapping results from both single-disease and multi-disease (MFM) fine-mapping; using the same groups for both methods allows a convenient comparison of results between the two methods.

As this grouping algorithm makes use of the model posteriors, we extended this grouping algorithm such that the groups are constructed independently for single-trait and multi-trait (flashfm) fine-mapping. That is, SNP groups are constructed for both methods using the same LD information, but groups may differ due to differences in the model PPs between the methods. Since the groups are constructed independently, the new algorithm incorporates a mapping between the group labels of the two sets of groups such that if a flashfm group overlaps a single-trait fine-mapping (STFM) group, the flashfm group label takes the name of the STFM group label. Sometimes a single STFM group overlaps multiple flashfm groups due to the higher precision of flashfm, which could reduce a SFTM group or split it into smaller groups. To account for this, if the STFM group has label A, each flashfm group that intersects A is given label A.1, A.2, etc. to denote that these flashfm groups were from the same larger group in STFM. This algorithm is available as the makeSNPgroups2 function in flashfm (https://github.com/jennasimit/flashfm/blob/master/R/group.multi.R)

### Multiple traits simulations

Simulations were carried out under a realistic scenario that mimics the MAF and *r*^2^ in the *IL2RA* region (345 SNPs in chromosome 10p-6030000-6220000 (GRCh37/hg19)). This region was selected as it has been previously shown to exhibit a tagging behaviour for causal variants; when there are two causal variants (A_1_=rs61839660 and D_1_=rs62626317), sometimes a different variant (B_1_=rs2104286), that is correlated with both causal variants, is detected as a single causal variant^9^; in this region this tagging behaviour was also observed for two causal variants, A_1_=rs61839660 and C_1_=rs11594656, jointly tagged by J_1_=rs706779. For this region, we generated a population of 100,000 individuals based on the CEU 1000 Genomes Phase 3 data^25^ (ftp://ftp.1000genomes.ebi.ac.uk/vol1/ftp/release/20130502/) using HapGen2^26^. Details of the LD between A_1_, B_1_, C_1_, D_1_, and J_1_ and their MAFs are given in Supplementary Data 1.1. Only variants with MAF >0.005 were included.

For each replication, a random sample of *N* individuals was selected from the population of 100,000. Causal variants for each trait were selected for a certain disease model, such that trait 1 has two causal variants (A_1_ and D_1_) and trait 2 either has two causal variants (A_1_ and C_1_); shared A_1_ with trait 1) or a single distinct causal variant (C_1_=rs11594656). Various values for the SNP effects were selected and a multiplicative model was assumed throughout. At each parameter configuration there are 300 replications.

For *M* traits, the measurement for trait *k* of individual *j*, *y_kj_*, is obtained from

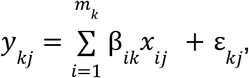

where *x_ij_* is the number of non-reference alleles of variant *i* for individual *j* (i.e. genotype score)*, β*_*ik*_ is the effect of causal variant *i* for trait *k*, *m*_k_ is the number of causal variants for trait *k*, and ε_*kj*_ is the *k^th^* element of the *j^th^* multivariate Normal distributed error variable with mean **0** and covariance Σ, which is the covariance matrix of the *M* traits.

For each replication, Expanded JAM was used for single-trait fine-mapping (SFTM) and used as input into flashfm. We used the makeSNPgroups2 function of flashfm to construct two sets of SNP groups, one set based on SFTM results and one set based on those of flashfm. Results from each method were then summarised based on their coinciding SNP groups, such that a 2-SNP model given by rs61839660 + rs62626317 = A_1_+D_1_ is represented by A+D; model A+D consists of all 2-SNP models where one SNP is from group A and the other is from group D. Group labels were assigned according to the variants contained within them, listed as in the beginning of this section for A, B, C, D, and J; if the variant is not captured by a group, then the group is given the label of the variant if it contains a variant that is in LD (r^2^>0.7) with the group variant.

Accuracy of the fine-mapping approaches was evaluated by comparing the mean group model PPs between the two methods. Precision was assessed by comparing the SNP group sizes (number of SNPs in each group) of the groups containing the causal variants for each method; within a replication, for each causal SNP group, the size of the flashfm group was subtracted from the JAM group and the median over the replications was used as a summary. To account for the possibility of a method giving a smaller group size, but not containing the causal variant, we also considered the coverage for each causal variant, defined as the proportion of simulations in which each causal variant is captured by a SNP group. In low power scenarios (e.g. small sample size) the methods may select the correct SNP groups with low PP, and as in a real analysis such SNP groups would not be followed up, we do not consider the SNP group size and coverage for a SNP group if its marginal PP < 0.1 in a replication; all replications are included in the summaries of the model PPs.

### Fine-mapping of cardiometabolic traits in a Ugandan cohort

To construct the regions for fine-mapping we used a criterion based on the centimorgan (cM) distance. In particular, for European ancestry populations, fine-mapping regions are often constructed using a boundary of +/− 0.1 cM around an association signal. When we applied this criterion to the Ugandan cohort, the constructed genome regions were excessively wide. In Park^27^ the authors show that a recombination rate of 0.1 corresponds to an LD of 0.4 for European ancestry population, meanwhile for African populations the same level of linkage disequilibrium corresponds to a recombination rate of 0.05. After investigating the dynamics between the recombination rate and the LD, we found that our data required a tighter criterion between 0.03 and 0.05 cM. In order to be conservative and aligned with the literature we selected a criterion of +/− 0.05 cM to define our fine mapping regions - for further details see Supplementary Material Section 2. For each region, we also checked a block of 200 SNPs beyond each bound of the region for any SNPs having r^2^>0.4 with the lead SNP in the region. If so, we extend the region to include such SNPs (for further details see Supplementary Material Section 2). This procedure came up with 56 regions with the number of traits by chromosome distribution shown in Supplementary Material Figure 2; 21,413,903 SNPs were considered in the analysis, (MAF>0.005). Most chromosomes had 2 trait-regions which represent 74% of the 56 regions.

Input to flashfm includes single trait fine-mapping results (model posterior probabilities) and single-SNP effect estimates for each trait from the GWAS. We consider two single trait fine-mapping algorithms that use GWAS summary statistics and a reference panel: i) FINEMAP^5^ and ii) JAM^4^. FINEMAP is a Bayesian stochastic search algorithm based on summary statistics from GWAS. As there are related samples, the effective sample size, as approximated from the GWAS summary statistics, is used for the number of measurements for each trait; this is easily calculated using the Neff function in flashfm (https://github.com/jennasimit/flashfm/blob/master/R/Neffective.R).

Besides single-trait fine-mapping results, flashfm also needs the single-SNP effect estimates, and a reference (or in-sample) genotype matrix or its covariance matrix. If GWAS summary statistics are available, but not single-trait fine-mapping results, then expanded JAM could be applied to provide input to flashfm. Only SNPs that exist in both the GWAS and the reference panel are carried forward for fine-mapping and this is automated within the flashfm software. Rather than fitting each joint model between traits, we derived an approximation to the joint Bayes’ factor (BF) that depends on the marginal BFs, single-SNP effect estimates for each trait, trait covariance matrix, and a reference panel. Based on the single-trait fine-mapping results, flashfm selects the top models from each trait, which are then carried forward for consideration in joint models between the traits. Preferably, models are selected for each trait based on the cumulative posterior probability (cpp; e.g. 0.99 by default), or a maximum number of models may be selected instead.

With exceptions to be clarified on a case-by-case basis, we consider a cpp of 0.99 in our analysis of the Uganda data. For ease of interpretation, as discussed in the SNP Groups section of Methods, main results are presented in terms of SNP groups, rather than individual SNPs; SNPs are grouped such that SNPs within each group are in high LD and rarely appear together in a model. Also, the average LD between groups is lower than 0.6.

For comparison purposes, in addition to results from single-trait fine-mapping from FINEMAP (and JAM in Supplementary Data 1.13) and flashfm (Supplementary Data 3), we also compare our results with the common approach of identifying the best model for the region, stepwise conditional regression^28,29^. In general, FINEMAP and JAM agree on the top model, though there are 6 regions for which JAM could not converge. For this reason, we focus on the more complete FINEMAP results (Supplementary Data 4).

## Data Availability

Complete results from our analysis are available in Supplementary Data 2.pdf. The GWAS results for the Uganda data have been previously published and summary results could be downloaded from https://www.ebi.ac.uk/gwas/publications/31675503#study_panel. There are no restrictions for data access and the Ugandan data may be requested from the original study authors (https://pubmed.ncbi.nlm.nih.gov/31675503/); genotype data are hosted by the European Bioinformatics Institute, under accession numbers EGAS00001001558/EGAD00010000965, EGAS00001000545/EGAD00001001639 and EGAS00001000545/EGAD00001005346.

## Code Availability

Flexible and shared information fine-mapping (flashfm) is freely available as an R library at https://jennasimit.github.io/flashfm/.

## Acknowledgements

JA is funded by the MRC (MR/R021368/1) which also supports NH. CW is funded by the Wellcome Trust (WT107881) and the MRC (MC_UU_00002/4). PN is funded by the MRC (MC UU 00002/9). NH, PN, CW, and JA are supported by the NIHR Cambridge BRC (BRC-1215-20014). This work was funded in part by an “Expanding excellence in England” award from Research England to IB. The views expressed are those of the author(s) and not necessarily those of the NHS, the NIHR or the Department of Health and Social Care. This research was funded in whole, or in part, by the Wellcome Trust [WT107881]. For the purpose of Open Access, the author has applied a CC BY public copyright licence to any Author Accepted Manuscript version arising from this submission.

## Author Contributions

JA initiated and coordinated the project, as well as developed and implemented the flashfm method. CW and PN contributed to the flashfm method implementation. MS provided the Uganda cardiometabolic genetics data. NH and JA performed statistical analyses. NH, MS, IB, CW, and JA interpreted results. IB contributed to expanding the scope of the flashfm method. JS cross-validated the flashfm method. JA, NH, and CW wrote the manuscript with input from all authors.

## Supplementary Data

**Supplementary Data 1: Details from simulations and summaries of the Ugandan cohort and its results.**

Simulation details and results summaries; Ugandan cohort trait summaries, trait correlation matrix, fine-mapped regions details, functional annotation of top SNPs in particular regions, top models for two single-trait approaches and coinciding flashfm results.

**Supplementary Data 2: Summary of single and multi-trait fine-mapping results for 56 regions in the Ugandan cohort.**

These regions have genetic associations for at least two of the 33 cardiometabolic traits. Both single-trait fine-mapping and flashfm results are given for the top models (with model PP), as well as number of SNPs in each SNP group, and the LD between SNP groups (for both fine-mapping methods).

**Supplementary Data 3: Flashfm results for all 56 regions that met criteria for fine-mapping.**

For each region, SNPs from each flashfm SNP group are listed, together with base pairs position (GRCh37/hg19 assembly), alleles, and RAF in Ugandan samples.

**Supplementary Data 4: FINEMAP results for all 56 regions that met criteria for fine-mapping.**

For each region, SNPs from each FINEMAP SNP group are listed, together with base pairs position (GRCh37/hg19 assembly), alleles, and RAF in Ugandan samples.

