## Supplementary Material for "Flashfm: A Flexible and Shared Information Fine-mapping Approach for Multiple Quantitative Traits"

### flashfm – Supplementary Material

#### Supplementary Section 1: FLexible And Shared information Fine-Mapping (flashfm) model description

##### 1.1 The multiple traits joint ABF is a function of marginal ABFs

We first suppose that we observe $N$ individuals, each with measurements for $M$ quantitative traits (transformed so that Normally distributed); later we relax this so that a subset of individuals may have missing measurements for some of the traits. Here, we find expressions for the ABF of causal SNP models for joint and marginal models and show that the information from single trait analyses could be used to evaluate the joint ABF.

To find expressions of the log(ABF) for each of the joint and marginal models we use the approximation based on BICs from the null and causal models ($\mathrm{BIC}_0$ and $\mathrm{BIC}_1$, respectively). The log(ABF) approximation $(\mathrm{BIC}_0 - \mathrm{BIC}_1)/2$, is expressed in terms of log likelihoods as

$$\log(ABF) \doteq l_1 - l_0 - k\log(N)/2, \tag{1}$$

where $k$ is the number of causal SNPs in the model and $l_1$ and $l_0$ are the log likelihoods of the causal and null models, evaluated at the maximum likelihood estimates.

An expression for the log(ABF) of a causal SNP model for a single trait is found after finding the log likelihoods for the relevant models in a Gaussian framework. Let $\mathbf{y}_j, j = 1,\ldots,M$ denote the vector of $N$ measurements for trait $j$, $\gamma_j$ represent a particular model with $k_j$ SNPs for trait $j$ and $\mathbf{X}_{\gamma_j}$ be a $N \times k_j$ matrix of genotypes scores for $k_j$ SNPs that are present in the model $\gamma_j$ for trait $j$. Under model $\gamma_j$ with causal SNPs $\mathbf{X}_{\gamma_j}$, the log-likelihood of a single trait $y_j$ is given by

$$
\begin{aligned}
l_1 &= -\frac{1}{2\hat{V}}(\mathbf{y}_j - \mathbf{X}_{\gamma_j}\hat{\boldsymbol{\beta}}_j)^T(\mathbf{y}_j - \mathbf{X}_{\gamma_j}\hat{\boldsymbol{\beta}}_j) - \frac{N}{2}\log(2\pi\hat{V}) \\
&= -\frac{N}{2}\left(1 + \log(2\pi) + \log\left(\frac{(\mathbf{y}_j - \mathbf{X}_{\gamma_j}\hat{\boldsymbol{\beta}}_j)^T(\mathbf{y}_j - \mathbf{X}_{\gamma_j}\hat{\boldsymbol{\beta}}_j)}{N}\right)\right),
\end{aligned}
\tag{2}
$$

where $\hat{V} = \frac{1}{N}(\mathbf{y}_i - \mathbf{X}_{\gamma_j}\hat{\boldsymbol{\beta}})_j^T(\mathbf{y}_i - \mathbf{X}_{\gamma_j}\hat{\boldsymbol{\beta}}_j)$ is the mle variance of the residuals from the fitted model.

Likewise, under the null model of no SNP associations and, without loss of generality, assuming mean 0 for the trait, the log likelihood is

$$
l_0 = -\frac{N}{2}\left(1 + \log(2\pi) + \log\left(\frac{\mathbf{y}_j^T\mathbf{y}_j)}{N}\right)\right). \tag{3}
$$

Then, using (2) and (3) in (1), the log(ABF) for a single trait $j$ is

$$
\begin{aligned}
\log(ABF_j) &= -\frac{N}{2}\log\left(\frac{(\mathbf{y}_j - \mathbf{X}_{\gamma_j}\hat{\boldsymbol{\beta}}_j)^T(\mathbf{y}_j - \mathbf{X}_{\gamma_j}\hat{\boldsymbol{\beta}}_j)}{\mathbf{y}_j^T\mathbf{y}_j}\right) - \frac{k_j}{2}\log(N) \\
&= -\frac{N}{2}\log\left(\frac{\hat{V}_{\gamma_j}}{\hat{V}_j}\right) - \frac{k_j}{2}\log(N),
\end{aligned}
\tag{4}
$$

where $V_j$ is the variance of trait $j$ and $\hat{V}_{\gamma_j}$ is the residual variance from model $\gamma_j$.

Next, consider $M$ traits that each have a possible model with possibly overlapping causal SNPs $\mathbf{X}_{\gamma_j}$ for trait $j$. Let $\mathbf{Y}$ be the $N \times M$ matrix of phenotypes and denote its rows by $\mathbf{y}_{i\cdot}$ ($M$-vector of trait values for individual $i$) and columns by $\mathbf{y}_j$ ($N$-vector of trait $j$ values). Under the null model for all traits, the joint log likelihood for $M$ traits is

$$
\begin{aligned}
l_0^M &= -\frac{1}{2}\sum_{i=1}^{N}\mathbf{y}_{i\cdot}^T\hat{\Sigma}_0^{-1}\mathbf{y}_{i\cdot} - \frac{N}{2}\log((2\pi)^M|\hat{\Sigma}_0|) \\
&= -\frac{N}{2}(M + M\log(2\pi) + \log(|\hat{\Sigma}_0|),
\end{aligned}
\tag{5}
$$

where $\hat{\Sigma}_0 = \frac{1}{N}\sum_{i=1}^{N}\mathbf{y}_{i\cdot}\mathbf{y}_{i\cdot}^T$ is the mle of the covariance matrix under the null model, having element $(i,j)$ given by $\frac{1}{N}\mathbf{y}_i^T\mathbf{y}_j$ (assuming mean 0 for all traits), and $\frac{NM}{2}$ is obtained by using properties of a scalar and the trace of a matrix.

Likewise, under model $\gamma_j$ with $k_j$ causal SNPs $\mathbf{X}_{\gamma_j}$ for trait $j$ the joint log likelihood is

$$l_1^M = -\frac{N}{2}(M + M\log(2\pi) + \log(|\hat{\Sigma}_1|), \tag{6}$$

where $\hat{\Sigma}_1$ is the mle of the covariance matrix under this model (covariance matrix of residuals) with element $(i, j)$ given by $\frac{1}{N}(\mathbf{y}_i - \mathbf{X}_{\gamma_i}\hat{\boldsymbol{\beta}}_i)^T(\mathbf{y}_j - \mathbf{X}_{\gamma_j}\hat{\boldsymbol{\beta}}_j)$. Then it follows from (1) with (5) and (6) that the log(ABF) of the joint model containing models $\gamma_j$ for trait $j$ is

$$\log(ABF^M) \quad = \quad -\frac{N}{2}\log(|\hat{\Sigma}_1\hat{\Sigma}_0^{-1}|) - \frac{K}{2}\log(N), \tag{7}$$

where $K = \sum_{i=1}^{M} k_i$ is the total number of SNP effects in the joint model. When there are two traits

$$\begin{aligned} |\hat{\Sigma}_1\hat{\Sigma}_0^{-1}| &= \begin{vmatrix} (\mathbf{y}_1 - \mathbf{X}_{\gamma_1}\hat{\boldsymbol{\beta}}_1)^T(\mathbf{y}_1 - \mathbf{X}_{\gamma_1}\hat{\boldsymbol{\beta}}_1) & (\mathbf{y}_1 - \mathbf{X}_{\gamma_1}\hat{\boldsymbol{\beta}}_1)^T(\mathbf{y}_2 - \mathbf{X}_{\gamma_2}\hat{\boldsymbol{\beta}}_2) \\ (\mathbf{y}_1 - \mathbf{X}_{\gamma_1}\hat{\boldsymbol{\beta}}_1)^T(\mathbf{y}_2 - \mathbf{X}_{\gamma_2}\hat{\boldsymbol{\beta}}_2) & (\mathbf{y}_2 - \mathbf{X}_{\gamma_2}\hat{\boldsymbol{\beta}}_2)^T(\mathbf{y}_2 - \mathbf{X}_{\gamma_2}\hat{\boldsymbol{\beta}}_2) \end{vmatrix} \begin{vmatrix} \mathbf{y}_1^T\mathbf{y}_1 & \mathbf{y}_1^T\mathbf{y}_2 \\ \mathbf{y}_1^T\mathbf{y}_2 & \mathbf{y}_2^T\mathbf{y}_2 \end{vmatrix}^{-1} \\ &= \begin{vmatrix} g_1 & h_{12} \\ h_{12} & g_2 \end{vmatrix} \begin{vmatrix} V_1 & C_{12} \\ C_{12} & V_2 \end{vmatrix}^{-1} \end{aligned}$$

where, $g_i = \frac{1}{N}(\mathbf{y}_i - \mathbf{X}_{\gamma_i}\hat{\boldsymbol{\beta}}_i)^T(\mathbf{y}_i - \mathbf{X}_{\gamma_i}\hat{\boldsymbol{\beta}}_i)$ is the residual variance of trait $i$, $h_{12} = \frac{1}{N}(\mathbf{y}_i - \mathbf{X}_{\gamma_i}\hat{\boldsymbol{\beta}}_i)^T(\mathbf{y}_j - \mathbf{X}_{\gamma_j}\hat{\boldsymbol{\beta}}_j)$ is the residual covariance for traits $i$ and $j$, and $C_{12}$ is the sample (unbiased) covariance between traits 1 and 2. Using (4), we obtain an expression for $g_i$ that is a function of log(ABF), sample size $N$, and the number of SNPs in model $\gamma_j$, $k_j$

$$g_j = \exp(-\frac{2}{N}\left(\log(ABF_j) + \frac{k_j}{2}\log(N)\right)V_j. \tag{8}$$

Then, using (4), the sum of the log(ABF) for $M$ single traits simplifies to

$$\begin{aligned}
\sum_{j=1}^{M} \log(ABF_j) &= -\frac{N}{2}\log\left(\frac{\prod_{j=1}^{M} g_j}{\prod_{j=1}^{M} V_j}\right) - \frac{K}{2}\log(N) \\
&= -\frac{N}{2}\log\left(\begin{vmatrix} g_1 & 0 & \cdots & 0 \\ 0 & g_2 & \cdots & 0 \\ \vdots & 0 & \ddots & \vdots \\ 0 & 0 & \cdots & g_M \end{vmatrix} \begin{vmatrix} V_1 & 0 & \cdots & 0 \\ 0 & V_2 & \cdots & 0 \\ \vdots & 0 & \ddots & \vdots \\ 0 & 0 & \cdots & V_M \end{vmatrix}^{-1}\right)
\end{aligned}$$

and it follows that the difference between the joint log(ABF) and the sum of the marginals is

$$\begin{aligned}
D_M &= \log(ABF^M) - \sum_{j=1}^{M} \log(ABF_j) \\
&= -\frac{N}{2}\log\left(\begin{vmatrix} g_1 & h_{12} & \cdots & h_{1M} \\ h_{12} & g_2 & \cdots & h_{2M} \\ \vdots & h_{i2} & \ddots & \vdots \\ h_{1M} & h_{2M} & \cdots & g_M \end{vmatrix} \begin{vmatrix} g_1 & 0 & \cdots & 0 \\ 0 & g_2 & \cdots & 0 \\ \vdots & 0 & \ddots & \vdots \\ 0 & 0 & \cdots & g_M \end{vmatrix}^{-1}\right) \\
&\quad + \frac{N}{2}\log\left(\begin{vmatrix} V_1 & C_{12} & \cdots & C_{1M} \\ C_{12} & V_2 & \cdots & C_{2M} \\ \vdots & C_{i2} & \ddots & \vdots \\ C_{1M} & C_{2M} & \cdots & V_M \end{vmatrix} \begin{vmatrix} V_1 & 0 & \cdots & 0 \\ 0 & V_2 & \cdots & 0 \\ \vdots & 0 & \ddots & \vdots \\ 0 & 0 & \cdots & V_M \end{vmatrix}^{-1}\right) \\
&= -\frac{N}{2}\left(\log\begin{vmatrix} 1 & \frac{h_{12}}{g_2} & \cdots & \frac{h_{1M}}{g_M} \\ \frac{h_{12}}{g_1} & 1 & \cdots & \frac{h_{2M}}{g_M} \\ \vdots & \frac{h_{i2}}{g_2} & \ddots & \vdots \\ \frac{h_{1M}}{g_1} & \frac{h_{2M}}{g_2} & \cdots & 1 \end{vmatrix} - \log\begin{vmatrix} 1 & \frac{C_{12}}{V_2} & \cdots & \frac{C_{1M}}{V_M} \\ \frac{C_{12}}{V_1} & 1 & \cdots & \frac{C_{2M}}{V_M} \\ \vdots & \frac{C_{i2}}{V_2} & \ddots & \vdots \\ \frac{C_{1M}}{V_1} & \frac{C_{2M}}{V_2} & \cdots & 1 \end{vmatrix}\right). \qquad (9)
\end{aligned}$$

where the second determinant is a constant $C$ with respect to the samples. Thus, the joint ABF is proportional to the product of the the marginal BFs and a function of the sample sizes and residual variances and covariances. Residual variances are approximated from the log(BF) for the coinciding model and trait, and residual covariances $h_{ij}$ are approximated as described below. If traits are standardised to have mean 0 and variance 1 and trait summary statistics are unavailable, we may use an estimate of the correlation matrix based on the GWAS summary statistics from the LD score regression approach (Buik-Sullivan et al. 2015).

We have $h_{ij} = \mathbf{y}_i^T \mathbf{y}_j - \mathbf{y}_i^T \mathbf{X}_{\gamma_j} \hat{\boldsymbol{\beta}}_j - \mathbf{y}_j^T \mathbf{X}_{\gamma_i} \hat{\boldsymbol{\beta}}_i + \hat{\boldsymbol{\beta}}^{*_i T} \mathbf{X}_{\gamma_i \cup \gamma_j}^T \mathbf{X}_{\gamma_i \cup \gamma_j} \hat{\boldsymbol{\beta}}_j^*$, where $\mathbf{X}_{\gamma_i \cup \gamma_j}$ is the genotype matrix with the SNPs contained in models $\gamma_i$ and/or $\gamma_j$ (i.e. union of model SNPs) and $\hat{\boldsymbol{\beta}}_i^*$ has the same effect estimates as $\hat{\boldsymbol{\beta}}_i$ at the SNPs in $\gamma_i$ and has 0 at SNPs that are only in in $\gamma_j$. The term $\mathbf{y}_i^T \mathbf{y}_j$ may be estimated from trait summary statistics. Terms of the form $\mathbf{y}_i^T \mathbf{X}_{\gamma_j} \hat{\boldsymbol{\beta}}_j = \sum_{k \in \gamma_j} S_{x_j y_i} \hat{\boldsymbol{\beta}}_{jk}$, where $S_{x_j y_i} = \mathrm{x}_j^T \mathbf{y}_i$ and $\hat{\beta}_{jk}$ is the trait $j$ effect estimate for the $k$th SNP in $\gamma_j$, and $S_{x_j y_i} = \sum_{k=1}^N x_{jk} y_{ik}$ is calculated from the the single-SNP effect estimates of the $k$th SNP from the trait $j$ model for trait $i$ (i.e. $S_{x_j y_i} = \hat{\beta}_{x_j}(N-1) V_{x_j} V_{y_i} + 2 * RAF_{x_j} * \hat{\mu_{Y_i}} N$, where $RAF_{x_j}$ is the reference allele frequency of $x_j$). The last term $\hat{\boldsymbol{\beta}}_i^T \mathbf{X}^T \mathbf{X} \hat{\boldsymbol{\beta}}_j$ relies on the effect estimates from the two trait models and $\mathbf{X}^T \mathbf{X}$ may be approximated from either the genotype or a suitable reference panel, as element $(i, j)$ of the matrix is

$$(\mathbf{X}^T \mathbf{X})_{ij} \doteq N(Cov(X_i, X_j) - E(X_i) E(X_j))$$

Thus, all quantities in the final expression for $D_M$ could be obtained from marginal analyses of the traits and summary information of the traits.

#### 1.2 Traits not measured for all samples

It is common for only a subset of individuals to have measurements for all traits. For simplicity, consider two traits that are both measured in $N$ individuals and let $N_{i-j}$ be the number of individuals with trait $i$ measured, but not trait $j$; the number of individuals with trait $i$ measured is $N_i = N + N_{i-j}$. Using all $N_i$

samples for trait $i$ we obtain the marginal $\log(ABF_i)$ as in (4), we then have

$$
\begin{aligned}
\log(ABF_1)+\log(ABF_2) &= -\frac{(N+N_{1-2})}{2}\log\left(\frac{\hat{V}_{\gamma_1}}{V_1}\right)-\frac{(N+N_{2-1})}{2}\log\left(\frac{\hat{V}_{\gamma_2}}{V_2}\right) \\
&\quad -\frac{k_1}{2}\log(N_1)-\frac{k_2}{2}\log(N_2) \\
&= -\frac{N}{2}\log\left(\frac{\hat{V}_{\gamma_1}\hat{V}_{\gamma_2}}{V_1V_2}\right) \\
&\quad +\frac{N_{1-2}}{N_1}\left(\log(ABF_1)+\frac{k_1}{2}\log(N_1)\right) \\
&\quad +\frac{N_{2-1}}{N_2}\left(\log(ABF_2)+\frac{k_2}{2}\log(N_2)\right) \\
&\quad -\frac{k_1}{2}\log(N_1)-\frac{k_2}{2}\log(N_2) \qquad (10)
\end{aligned}
$$

The joint ABF is obtained in a similar manner, where extra terms are needed to account for the individuals with measurements for only one of the two traits. Let $\mathbf{y}_{i-j}$ denote the trait $i$ measurements for individuals that do not have trait $j$ measured. It follows that for the likelihood under the null, we have

$$
\begin{aligned}
l_0 &= -N-\frac{N}{2}\log(V_1V_2-cov^2(y_1,y_2))-N\log(2\pi) \\
&\quad -\frac{N_{1-2}}{2}(1+\log(2\pi)+\log\left(\frac{\mathbf{y}_{1-2}^T\mathbf{y}_{1-2}}{N_{1-2}}\right)) \\
&\quad -\frac{N_{2-1}}{2}(1+\log(2\pi)+\log\left(\frac{\mathbf{y}_{2-1}^T\mathbf{y}_{2-1}}{N_{2-1}}\right)) \qquad (11)
\end{aligned}
$$

and under the models $\gamma_1,\gamma_2$, for traits 1 and 2, where $\hat{\beta}_i$ is based on all $N_i$ samples with measurements and $\hat{\beta}_{i-j}$ is based on the $N_{i-j}$ samples with trait $i$ measured and not trait $j$ , we have

$$
\begin{aligned}
l_1 &= -N-N\log(2\pi)-\frac{N}{2}\log(\hat{V}_{\gamma_1}\hat{V}_{\gamma_2}-\hat{C}^2_{\gamma_1,\gamma_2}) \\
&\quad -\frac{N_{1-2}}{2}(1+\log(2\pi)+\log\left(\frac{\mathbf{y}_{1-2}^T\mathbf{y}_{1-2}-\mathbf{y}_{1-2}^T\mathbf{X}_{\gamma_1}\hat{\beta}_{1-2}}{N_{1-2}}\right)) \\
&\quad -\frac{N_{2-1}}{2}(1+\log(2\pi)+\log\left(\frac{\mathbf{y}_{2-1}^T\mathbf{y}_{2-1}-\mathbf{y}_{2-1}^T\mathbf{X}_{\gamma_2}\hat{\beta}_{2-1}}{N_{2-1}}\right)), \qquad (12)
\end{aligned}
$$

From (11) and (12) we get

$$l_1 - l_0 = -\frac{N}{2}\log\left(\frac{\hat{V}_{\gamma_1}\hat{V}_{\gamma_2} - \hat{C}^2_{\gamma_1,\gamma_2}}{V_1V_2 - C^2_{1,2}}\right) - \frac{N_{1-2}}{2}\log\left(\frac{\mathbf{y}^T_{1-2}\mathbf{y}_{1-2} - \mathbf{y}^T_{1-2}\mathbf{X}_{\gamma_1}\hat{\boldsymbol{\beta}}_{1-2}}{\mathbf{y}^T_{1-2}\mathbf{y}_{1-2}}\right) - \frac{N_{2-1}}{2}\log\left(\frac{\mathbf{y}^T_{2-1}\mathbf{y}_{2-1} - \mathbf{y}^T_{2-1}\mathbf{X}_{\gamma_2}\hat{\boldsymbol{\beta}}_{2-1}}{\mathbf{y}^T_{2-1}\mathbf{y}_{2-1}}\right) \quad (13)$$

Notice that $\log\left(\frac{\mathbf{y}^T_{1-2}\mathbf{y}_{1-2} - \mathbf{y}^T_{1-2}\mathbf{X}_{\gamma_1}\hat{\boldsymbol{\beta}}_{1-2}}{\mathbf{y}^T_{1-2}\mathbf{y}_{1-2}}\right)$ is the same form as the log ratio in the marginal $\log(ABF_1)$ expression in (4) and is based on a subset of size $N_{1-2}$ from the $N_1$ samples. Treating this log ratio based on the $N_{1-2}$ samples as an approximation to that based on all $N_1$ samples, we have

$$\log\left(\frac{\mathbf{y}^T_{1-2}\mathbf{y}_{1-2} - \mathbf{y}^T_{1-2}\mathbf{X}_{\gamma_1}\hat{\boldsymbol{\beta}}_{1-2}}{\mathbf{y}^T_{1-2}\mathbf{y}_{1-2}}\right) \doteq -\frac{2}{N_1}(\log(ABF_1) + \frac{k_1}{2}\log(N_1))$$

Likewise for trait 2.

Now, using $l_1 - l_0$ in (13), $\log(ABF^M) = l_1 - l_0 - \frac{k_1}{2}\log(N_1) - \frac{k_2}{2}\log(N_2)$, where $N_j, j = 1,2$ are used, being the number of samples in the estimates for $\hat{\boldsymbol{\beta}}_j$. So,

$$\begin{aligned}\log(ABF^M) &= -\frac{N}{2}\log\left(\frac{\hat{V}_{\gamma_1}\hat{V}_{\gamma_2} - \hat{C}^2_{\gamma_1,\gamma_2}}{V_1V_2 - C^2_{1,2}}\right) \\ &\quad + \frac{N_{1-2}}{N_1}(\log(ABF_1) + \frac{k_1}{2}\log(N_1)) \\ &\quad + \frac{N_{2-1}}{N_2}(\log(ABF_2) + \frac{k_2}{2}\log(N_2)) \\ &\quad - \frac{k_1}{2}\log(N_1) - \frac{k_2}{2}\log(N_2) \\ &= -\frac{N}{2}\log\left(\begin{vmatrix} g_1 & h_{12} \\ h_{12} & g_2 \end{vmatrix}\begin{vmatrix} V_1 & C_{12} \\ C_{12} & V_2 \end{vmatrix}^{-1}\right) \\ &\quad + \frac{N_{1-2}}{N_1}\log(ABF_1) + \frac{N_{2-1}}{N_2}\log(ABF_2) \\ &\quad - \frac{N_1 - N_{1-2}}{N_1}\frac{k_1}{2}\log(N_1) - \frac{N_2 - N_{2-1}}{N_2}\frac{k_2}{2}\log(N_2) \end{aligned} \quad (14)$$

Then, when finding the difference between the joint ABF and sum of marginal ABFs in this setting, the additional terms accounting for missing measurements for one of the traits cancel out between the joint and sum of marginal ABFs, giving a similar form to when all traits are measured.

$$
\begin{aligned}
D_M &= \log(ABF^M) - \sum_{j=1}^{2} \log(ABF_j) \\
&= -\frac{N}{2}\left(\log\begin{vmatrix} 1 & \frac{h_{12}}{g_2} \\ \frac{h_{12}}{g_1} & 1 \end{vmatrix} - \log\begin{vmatrix} 1 & \frac{C_{12}}{V_2} \\ \frac{C_{12}}{V_1} & 1 \end{vmatrix}\right)
\end{aligned} \tag{15}
$$

.

In general, for $M$ traits, the joint $\log(ABF)$ is expressed as

$$
\log(ABF^M) = \sum_{j=1}^{M} \log(ABF_j) + D_M,
$$

where $D_M$ for two traits is as in (16) and for $M = 3,4,5$, expressions follow.

When there are more than two traits and some have missing data, additional terms to account for missing measurements are present in the expression for the $\log(ABF^M)$. The derivations for 3-6 traits generalise from the two trait scenario and we use the notation $\delta_{ijk}$ to represent the term given in (9) for traits $i, j, k$ and analogously for a larger number of traits. In addition, $N_{ijk}$ is the number of individuals with traits $i, j$ and $k$ all measured, $N_{ij-k}$ denotes the number of individuals with both traits $i, j$ measured and not trait $k$, $N_{i-jk}$ denotes the number of individuals with trait $i$ measured, but neither of traits $j$ and $k$, and analogous notation is used for more/different combinations of traits.

**3 traits:**

$$
D_M = \delta_{123} + \frac{N_{12-3}}{N_{12}}\delta_{12} + \frac{N_{13-2}}{N_{13}}\delta_{13} + \frac{N_{23-1}}{N_{23}}\delta_{23},
$$

where

$$
N_{jk-l} = N_{jk} - N_{jkl};
$$

$$
N_{j-kl} = N_j - N_{jk} - N_{jl} + N_{123};
$$

**4 traits**, denote $\mathscr{M} = \{1,2,3,4\}$:

$$D_M = \delta_{1234} + \sum_{j=1;k,l,m\in\mathscr{M}\setminus\{j\}}^{4} \frac{N_{klm-j}}{N_{klm}} \delta_{klm} + \sum_{j=1;k\neq j;l,m\in\mathscr{M}\setminus\{j,k\}}^{4} \frac{N_{jk-lm}}{N_{jk}} \delta_{jk},$$

where

$$N_{jkl-m} = N_{jkl} - N_{jklm};$$

$$N_{jk-lm} = N_{jk} - N_{jkl} - N_{jkm} + N_{1234};$$

**5 traits**, denote $\mathscr{M} = \{1,2,3,4,5\}$:

$$D_M = \delta_{12345} + \sum_{i=1;j,k,l,m\in\mathscr{M}\setminus\{j\}}^{5} \frac{N_{jklm-i}}{N_{jklm}} \delta_{jklm-i} + \sum_{i=1;j\neq i;k,l,m\in\mathscr{M}\setminus\{i,j\}}^{5} \frac{N_{ij-klm}}{N_{ij}} \delta_{ij} + \sum_{i=1;j,k\neq i;l,m\in\mathscr{M}\setminus\{i,j,k\}}^{5} \frac{N_{ijk-lm}}{N_{ijk}} \delta_{ijk}$$

where

$$N_{ijkl-m} = N_{ijkl} - N_{ijklm};$$

$$N_{ijk-lm} = N_{ijk} - N_{ijkl} - N_{ijkm} + N_{12345};$$

$$N_{ij-klm} = N_{ij} - N_{ijk} - N_{ijl} - N_{ijm} + N_{ijkl} + N_{ijkm} + N_{ijlm} - N_{12345};$$

In our flashfm software we include a "fastapprox" option that gives a quicker calculation by ignoring the extra adjustment terms. This is recommended when there are not many missing trait measurements and when a quicker answer is required; by default `fastapprox=FALSE`, but for 6 traits, only `fastapprox=TRUE` is available.

The prior probability for the joint models includes a term that gives more weight to joint models that have a shared causal variant between the traits; this term $\kappa$ is derived in a combinatorial manner and is identical to that used in MFM (Asimit et al. 2019). As in MFM, a correction term $\tau$ is also included to ensure that the prior probability of a certain number of SNPs in a model is the same for any value of $\kappa$. When $\kappa = 1$, there is no weight for joint models with shared causal variants and the flashfm PP for each model for a given trait is the same to what one would obtain from single-trait fine-mapping, which we also refer to as independent fine-mapping, as it does not make use of data from other traits.

##### 1.3 Implementation

There are two options for implementing flashfm. If single-trait fine-mapping results have not already been obtained, they may be generated within flashfm using an extended version of JAM (Joint Analysis of Marginal summary statistics; Newcombe, Conti, Richardson, 2016) - this requires GWAS summary statistics and either a genotype matrix or both a genotype covariance matrix and MAF vector from a reference panel or in-sample study. Alternatively, any single-trait fine-mapping approach that output model PPs, such as FINEMAP (Benner et al. 2016) could be used as input - this still requires GWAS summary statistics and either a genotype matrix or both a genotype covariance matrix and MAF vector from a reference panel or in-sample study.

JAM assesses the joint effect of multiple SNPs on a trait in an integrated Bayesian penalized regression framework, outputting the posterior probabilities (PP) for the multi-SNP models. This allows us to identify the models with non-negligible evidence that should be the focus when assessing joint models between multiple traits. As JAM operates on a set of tag SNPs due to colinearity issues, we have extended it such that all models with PP $> 0.005$ are expanded by their tag SNPs in the same manner as GUESSFM (https://github.com/chr1swallace/GUESSFM; Wallace et al. 2015). This is done by substituting each tag SNP in a model by each of the SNPs that it tags so that if SNPs 1 and 2 are in a model and they each tag $t_1$ and $t_2$ SNPS, respectively, the model expands into $(t_1+1)(t_2+1)$ models, for which ABFs are found using (4; $\hat{\beta}$ for multi-SNP models are obtained from the single-SNP $\hat{\beta}$ and the genotype matrix (or reference panel) of the SNPs in the model.

Using a binomial prior distribution we may then find PPs for all of the expanded (and original) models. For ease of interpretation, we also construct SNP groups (using the snp.picker function of GUESSFM) such that SNPs in the same group could be substituted for one another; SNPs in the same group are in high LD and are rarely selected together in models. The results are then summarised in terms of SNP group PPs by summing over SNP models that fall into each SNP group model; the PP for the SNP group model $A+B$ is the sum over PPs from all models with one SNP from $A$ and one SNP from $B$.

The posterior probability of model $\gamma_1$ for trait 1 is proportional to a sum of the posterior probabilities of all configurations $C_{1,j},\ j=1,\ldots,n$. Let $I_{i,j}$ be an indicator function, taking the value 1 if $\gamma_i \cap \gamma_j \neq \varnothing$ and 0 otherwise, and let $\delta_{ij} = \exp D_{ij}$ Then

$$
\begin{aligned}
Pr(\gamma_i \text{ for trait } 1|\text{Data}) &\propto \sum_j p_i p_j ABF_i ABF_j \times \delta_{ij} \kappa^{I_{i,j}} \tau_{ij} \\
&= PP_i \left( \sum_{j:I_{i,j}=0} \delta_{ij}\tau_{ij}PP_j + \kappa \sum_{j:I_{i,j}=i} \tau_{ij}\delta_{ij}PP_j \right) \\
&= PP_i \left( \sum_j \delta_{ij}\tau_{ij}PP_j + (\kappa-1) \sum_{j:I_{i,j}=i} \delta_{ij}\tau_{ij}PP_j \right) \\
&= PP_i \left( 1 + (\kappa-1) \frac{\sum_{j:I_{i,j}=1} \delta_{ij}\tau_{ij}PP_j}{\sum_j \delta_{ij}\tau_{ij}PP_j} \right)
\end{aligned}
$$

Rather than considering all model combinations, we reduce the model space by setting a cumulative posterior probability threshold (e.g. cpp=0.99). For each trait, we use the single-trait fine-mapping results to order the models by PP and retain those for which the sum of their PPs is below 0.99. As these $\delta_{ij}$ terms depend on the SNPs that are included in each model, a loop over the model combinations is required to make these small calculations.

#### 1.4 Related Individuals Implementation

The above derivations are based on a sample of unrelated individuals. If the proportion of related individuals is relatively large such that their removal would be a noticeable loss in data, rather than excluding related samples, an alternative approach is considered. First, single-SNP mixed linear models that account for relatedness are fit for each trait using GEMMA (Genome-wide Efficient Mixed Model Association; Zhou and Stephens, 2013). The output from GEMMA includes the relatedness-adjusted effect estimates $\hat{\beta}$ of each SNP for one trait, which may then be used as input to JAM (Newcombe, Conti, Richardson, 2016) or FINEMAP (Benner et al. 2016), as above, to identify the models with non-negligible evidence. As the single-SNP effect estimates are adjusted for relatedness, they may be used together with the genotype matrix of unrelated samples (or reference panel) as above to obtain $\hat{\boldsymbol{\beta}}$ for multi-SNP models, which are needed to get $\log(ABF)$ as in (4); as these effect estimates are adjusted for relatedness, they may be treated as if obtained from an unrelated sample. The effective sample size is used as $N$ in the $\log(ABF)$ calculation.

#### Supplementary Section 2: Region construction for fine-mapping in the Ugandan cohort

In order to obtain more precision in the construction of the fine mapping regions we consider the centimorgan (cM) genetic distance between SNPs. Approximately 80% of the SNPs (hg19/build 37) in the Ugandan data set do not map to a cM (reference panel) position so missing values were imputed using linear interpolation.

We then considered the GWAS for each of the 33 traits and selected the SNPs using a p-value threshold of $1 \times 10^{-6}$.

Next, we sorted the p-values of the selected SNPs from all trait GWAS in descending order and removed any duplicated SNPs. Finally, regions were constructed using the following steps:

1. For the most significant SNP, construct a region $\pm 0.05cM$ around the SNP

2. Block correlation check: for a block of 200 SNPs on each side of the region we compute the $r^2$ (LD) between the 200 SNPs block and the lead SNP in the region. We extend the region until the SNP of the block with highest $r^2$ (considering a minimum of 0.4).

3. Repeat the procedure for subsequent SNPs, checking first if the SNP belongs to any previously constructed region.

Applying this procedure to the Ugandan data set we obtained 56 regions detailed in Supplementary Figure 2.

### Supplementary Section 3: Figures

Supplementary Figure 1: **Comparison of flashfm and single trait fine-mapping when there is a shared causal variant.** Causal variants were simulated for two traits: trait 1 has causal variants A+C, while trait 2 has A+D causal variants, where $\beta_A = log(1.4)$ for both traits and $\beta_C = \beta_D = log(1.25)$. Both panels show the mean posterior probability throughout a variation of the sample size from 1000 to 5000, with no missing data. Source data are provided in Supplementary Data 1 Tables 1-5.

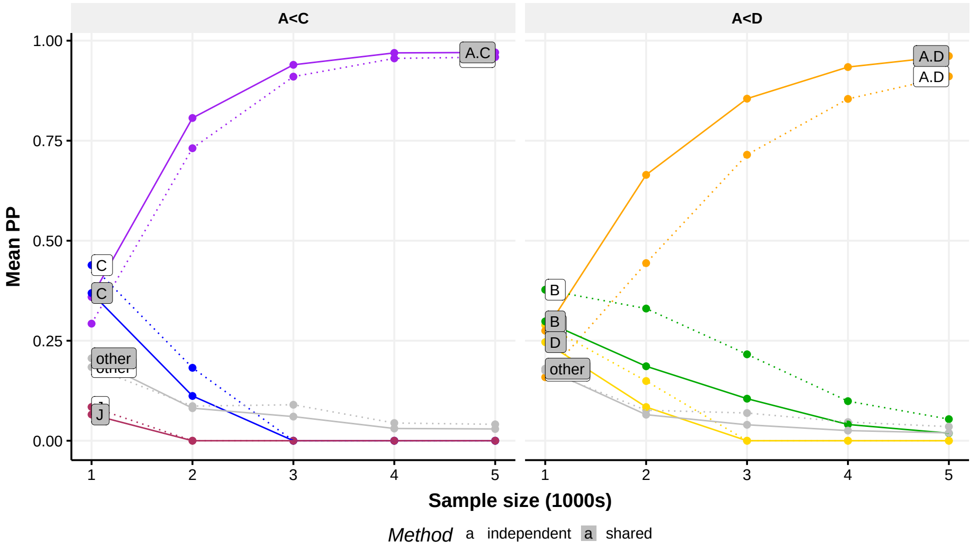

Supplementary Figure 2: **Distribution of regions by traits and chromosome.** Most of the regions are formed by 2 traits, while chromosomes 16 and 19 contain some regions with signals from 4-6 traits.

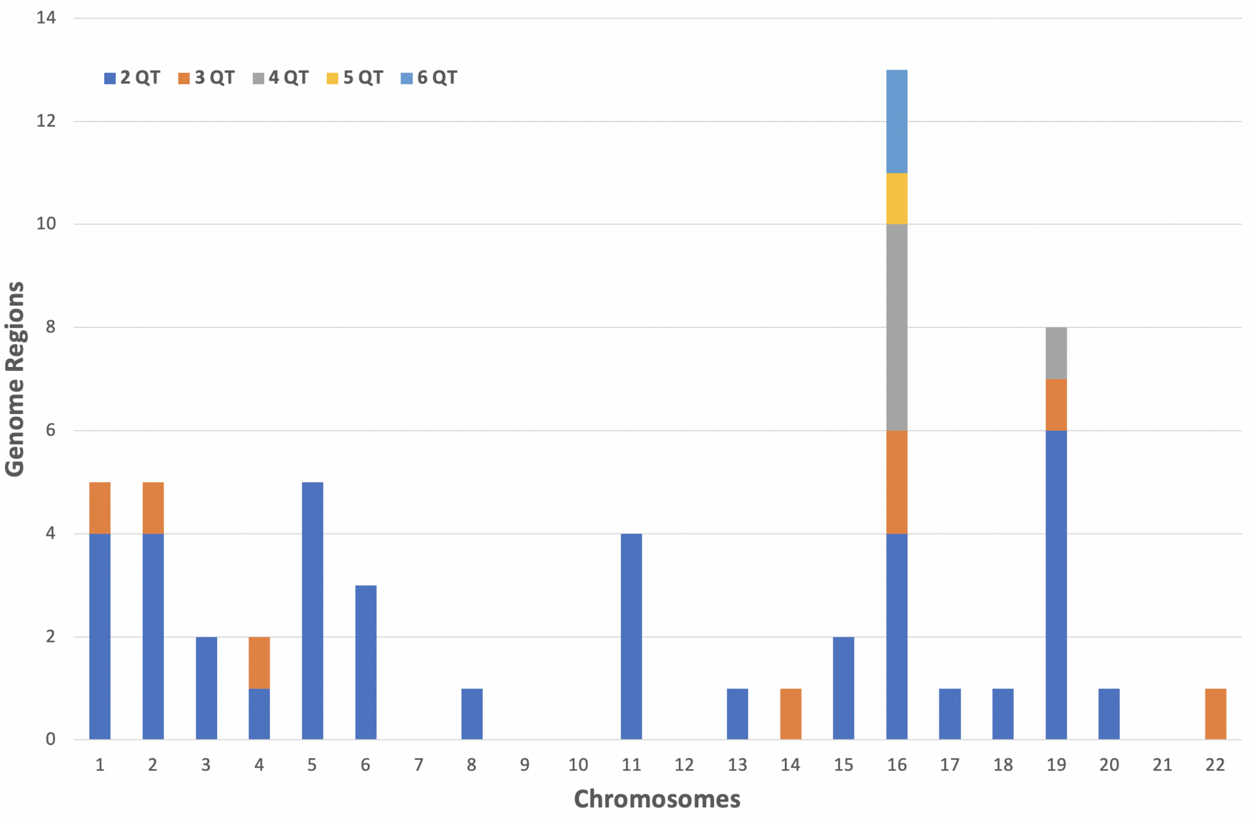

Supplementary Figure 3: **Quantitative traits correlation heatmap**. Correlation blocks can be observed between different traits. There is one correlation block of anthropometric traits as height, weight, body mass index, etc. There is another block that relates mature red blood cell traits such as mchc, mch, hct, mcv, hgb, rbc. Finally there is a last clear correlation block for lipid traits such as lowdlipo, highdlipo, cholesterol and triglycerides.

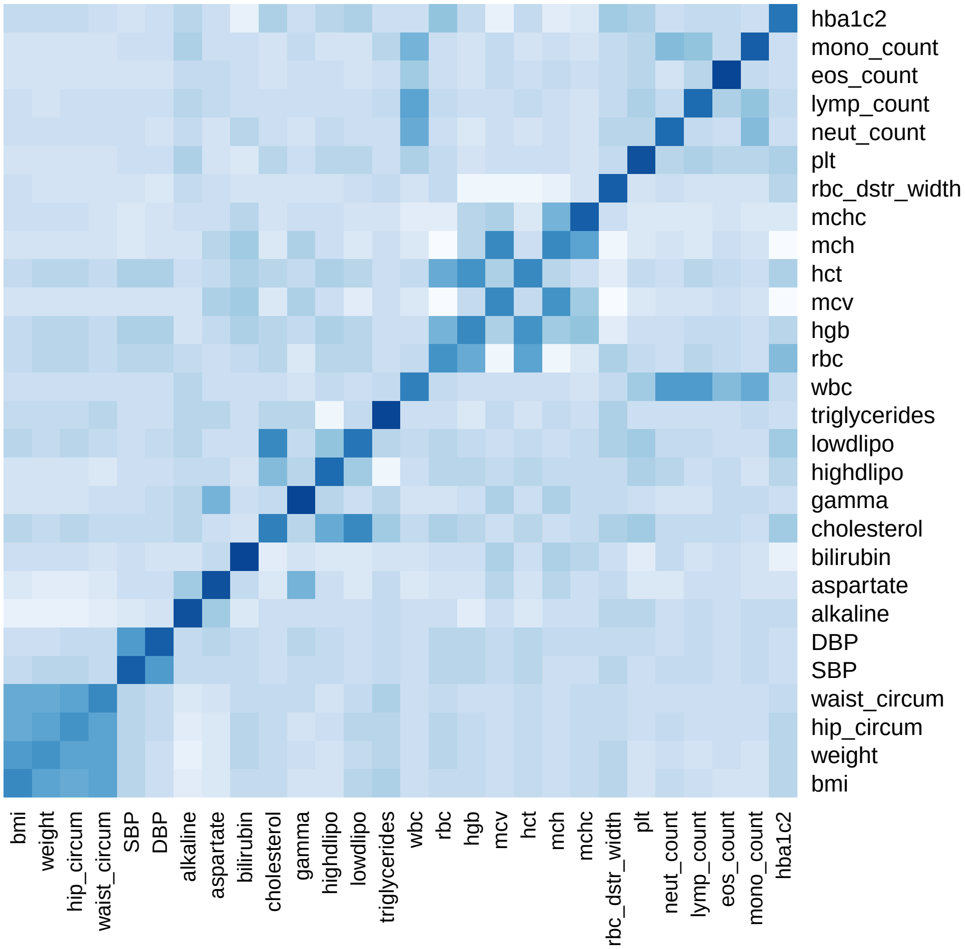
