## Supplementary Data 2 for "Flashfm: A Flexible and Shared Information Fine-mapping Approach for Multiple Quantitative Traits"

### Flashfm – Supplementary Data 2: Uganda fine-mapping results

Here we present the complete Uganda fine-mapping results from flashfm and independent fine-mapping. For all regions the independent fine-mapping method is FINEMAP, except for one region (16:302161-440165) where we used JAM; this region has 6 traits with multi-SNP models that was computationally quicker with JAM. For all the regions that involve 2 or 3 traits we consider a cumulative posterior probability (cpp) of 0.99. For regions with more than 3 traits we consider a cpp of 0.95 unless otherwise stated.

For each region we present two figures that show (i) the average LD between SNP groups constructed under single-trait fine-mapping and those constructed under flashfm; (ii) a pair of plots, each showing the average LD between SNP groups within each method. If independent fine-mapping and flashfm both favour the same single SNP model, a Venn diagram of the SNP group is presented. This illustrates how many SNPs from the flashfm group are a subset of the independent fine-mapping group.

Sometimes an independent fine-mapping group is split into two groups by flashfm. This is shown, for example by group A for independent fine-mapping and groups A.1 and A.2 from flashfm.

| | Stepwise | | Independent | | Flashfm | |
|---|---|---|---|---|---|---|
| Traits | SNP/Model | P-value | Model | PP | Model | PP |
| lowdlipo | 1:109817590/A | 2.63893e-19 | A | 0.722 | A | 0.797 |
| cholesterol | 1:109817590/A | 4.98465e-14 | A | 0.692 | A | 0.774 |

Supplementary Table 1: Fine-mapping results for 1:109811755-109825591 by independent fine-mapping and Flashfm. Stepwise results are also given.

| | SNP Group |
|---|---|
| | A |
| Independent | 2 |
| | 1:109817590 |
| Flashfm | 2 |
| | 1:109817590 |

Supplementary Table 2: SNP group sizes and representative variant for groups constructed by independent fine-mapping and by Flashfm for region 1:109811755-109825591.

Supplementary Figure 1: Venn diagram of SNP group constructed under indepenedent fine-mapping compared to that under flashfm for region 1:109811755-109825591. This shows that the flashfm group is a subset of, or the same as, the group form independent fine-mapping.

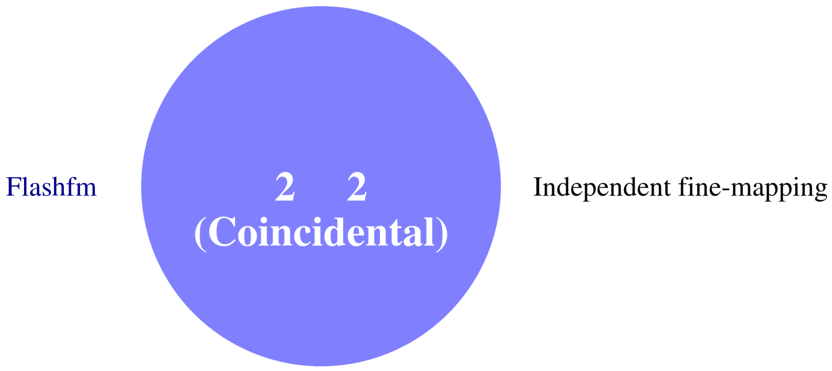

| | Stepwise | | Independent | | Flashfm | |
|---|---|---|---|---|---|---|
| Traits | SNP/Model | P-value | Model | PP | Model | PP |
| lowdlipo | 1:55520445/A | 1.05319e-14 | A | 0.496 | A | 0.628 |
| | – | – | A+C | 0.092 | A+C | 0.061 |
| cholesterol | 1:55520445/A | 2.37584e-09 | A | 0.639 | A | 0.758 |

Supplementary Table 3: Fine-mapping results for 1:55517883-55674945 by independent fine-mapping and Flashfm. Stepwise results are also given.

| | SNP Group | |
|---|---|---|
| | A | C |
| Independent | 4 | 1 |
| | 1:55518622 | 1:55522083 |
| Flashfm | 3 | 1 |
| | 1:55518622 | 1:55522083 |

Supplementary Table 4: SNP group sizes and representative variant for groups constructed by independent fine-mapping and by Flashfm for region 1:55517883-55674945.

Supplementary Figure 2: Average LD between SNP groups constructed under independent fine-mapping with those from flashfm for region 1:55517883-55674945.

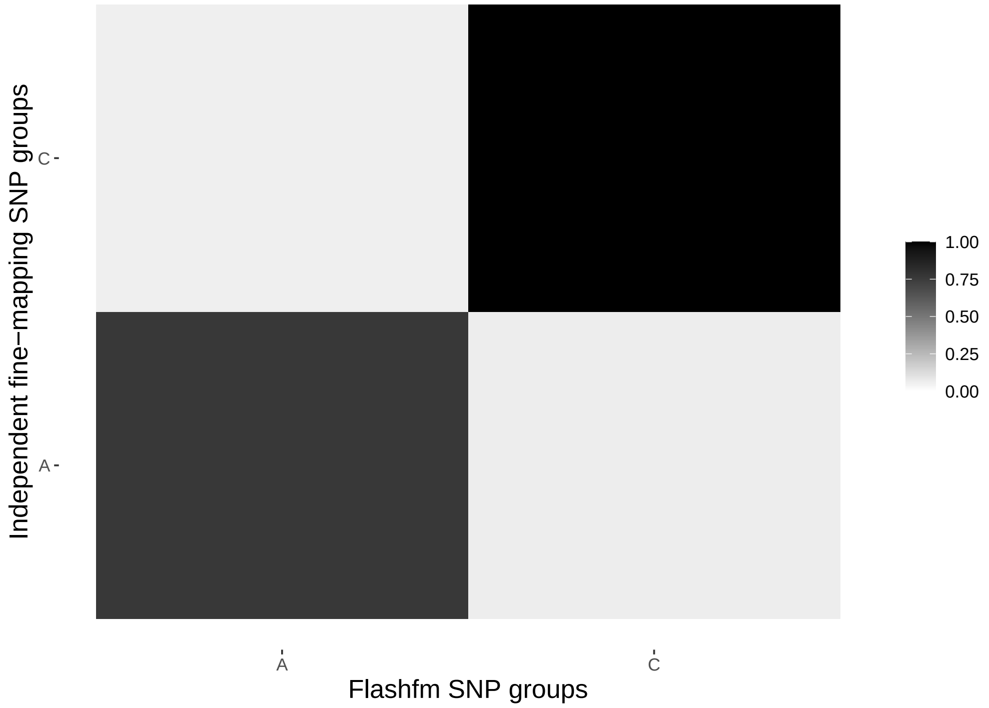

Supplementary Figure 3: Average LD between SNP groups constructed under independent fine-mapping (left) and Flashfm (right) for region 1:55517883-55674945.

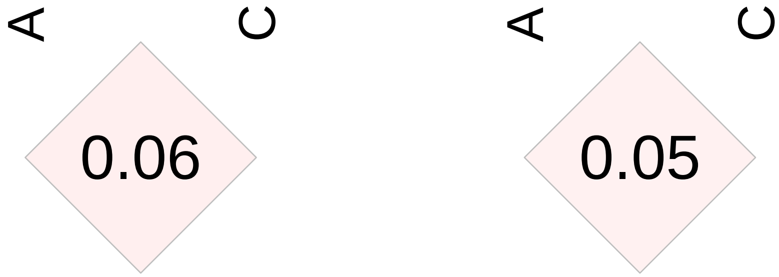

| | Stepwise | | Independent | | Flashfm | |
|---|---|---|---|---|---|---|
| Traits | SNP/Model | P-value | Model | PP | Model | PP |
| cholesterol | 1:17784317/D | 4.24918e-08 | D | 0.279 | D | 0.496 |
| | – | – | B+D | 0.274 | B+D | 0.18 |
| | – | – | A+D | 0.085 | A+D | 0.069 |
| | – | – | A+B+D | 0.057 | – | – |
| lowdlipo | 1:17784317/D | 5.54811e-07 | B+D | 0.252 | B+D | 0.277 |
| | – | – | A+B+D | 0.160 | D | 0.154 |
| | – | – | A+D | 0.068 | A+B+D | 0.089 |
| | – | – | D | 0.068 | A+D | 0.069 |
| | – | – | B+C+D | 0.051 | – | – |

Supplementary Table 5: Fine-mapping results for 1:17700017-17873294 by independent fine-mapping and Flashfm. Stepwise results are also given.

| | SNP Group | | | |
|---|---|---|---|---|
| | A | B | C | D |
| Independent | 29 | 1 | 14 | 1 |
| | 1:17730238 | 1:17765792 | 1:17843968 | 1:17784317 |
| Flashfm | 25 | 1 | 9 | 1 |
| | 1:17781513 | 1:17765792 | 1:17847630 | 1:17784317 |

Supplementary Table 6: SNP group sizes and representative variant for groups constructed by independent fine-mapping and by Flashfm for region 1:17700017-17873294.

Supplementary Figure 4: Average LD between SNP groups constructed under independent fine-mapping with those from flashfm for region 1:17700017-17873294.

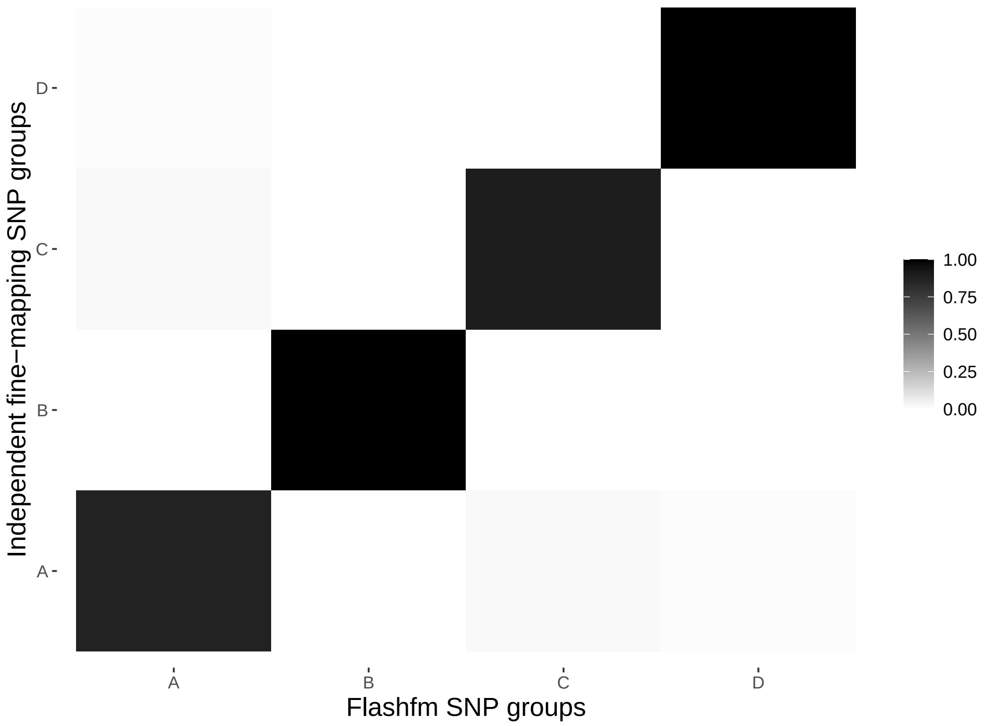

Supplementary Figure 5: Average LD between SNP groups constructed under independent fine-mapping (left) and Flashfm (right) for region 1:17700017-17873294.

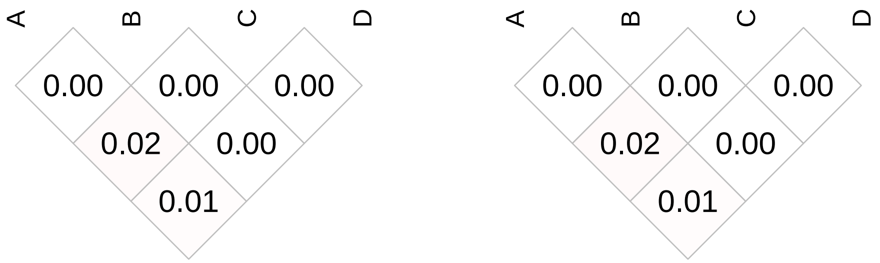

| | Stepwise | | Independent | | Flashfm | |
|---|---|---|---|---|---|---|
| Traits | SNP/Model | P-value | Model | PP | Model | PP |
| hip_circum | 1:180143575/C | 8.6265e-08 | C | 0.463 | C | 0.495 |
| | – | – | D | 0.297 | D | 0.293 |
| waist_circum | 1:180171741/D | 1.8039e-07 | D | 0.606 | D | 0.68 |
| | – | – | C | 0.086 | C | 0.055 |
| weight | 1:180143575/C | 5.30306e-07 | C | 0.635 | C | 0.68 |
| | – | – | A+C | 0.053 | – | – |

Supplementary Table 7: Fine-mapping results for 1:180122985-180192543 by independent fine-mapping and Flashfm. Stepwise results are also given.

| | SNP Group | | |
|---|---|---|---|
| | A | C | D |
| Independent | 9 | 15 | 2 |
| | 1:180123815 | 1:180187342 | 1:180171741 |
| Flashfm | 9 | 14 | 2 |
| | 1:180123815 | 1:180187342 | 1:180171741 |

Supplementary Table 8: SNP group sizes and representative variant for groups constructed by independent fine-mapping and by Flashfm for region 1:180122985-180192543.

Supplementary Figure 6: Average LD between SNP groups constructed under independent fine-mapping with those from flashfm for region 1:180122985-180192543.

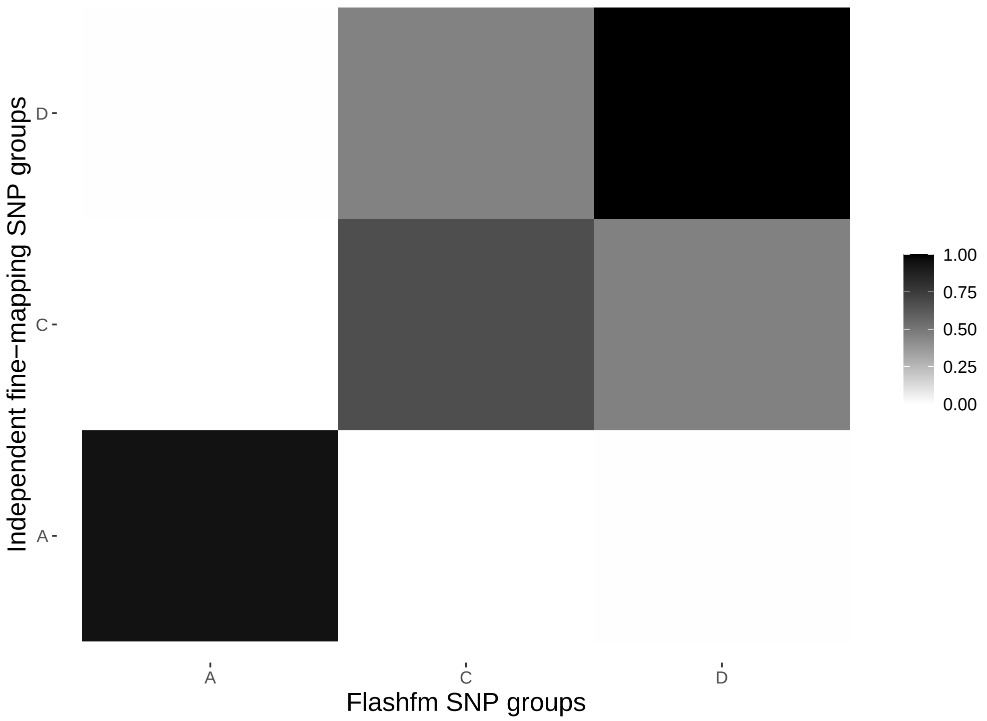

Supplementary Figure 7: Average LD between SNP groups constructed under independent fine-mapping (left) and Flashfm (right) for region 1:180122985-180192543.

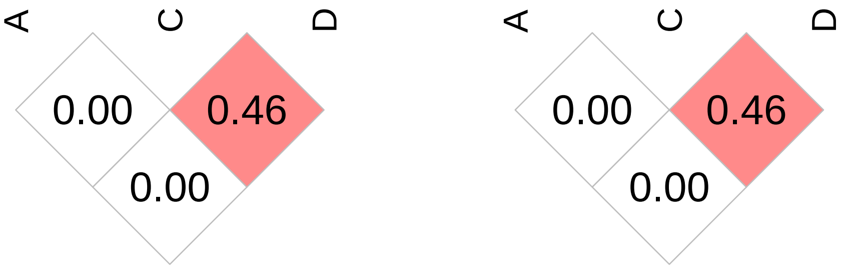

| | Stepwise | | Independent | | Flashfm | |
|---|---|---|---|---|---|---|
| Traits | SNP/Model | P-value | Model | PP | Model | PP |
| wbc | 1:220425014/B | 3.13673e-07 | B | 0.579 | B | 0.505 |
| | – | – | – | – | A | 0.236 |
| mono_count | 1:220360197/A | 5.88953e-07 | A | 0.575 | A | 0.655 |

Supplementary Table 9: Fine-mapping results for 1:220212231-220479374 by independent fine-mapping and Flashfm. Stepwise results are also given.

| | SNP Group | |
|---|---|---|
| | A | B |
| Independent | 1 | 1 |
| | 1:220360197 | 1:220425014 |
| Flashfm | 1 | 1 |
| | 1:220360197 | 1:220425014 |

Supplementary Table 10: SNP group sizes and representative variant for groups constructed by independent fine-mapping and by Flashfm for region 1:220212231-220479374.

Supplementary Figure 8: Average LD between SNP groups constructed under independent fine-mapping with those from flashfm for region 1:220212231-220479374.

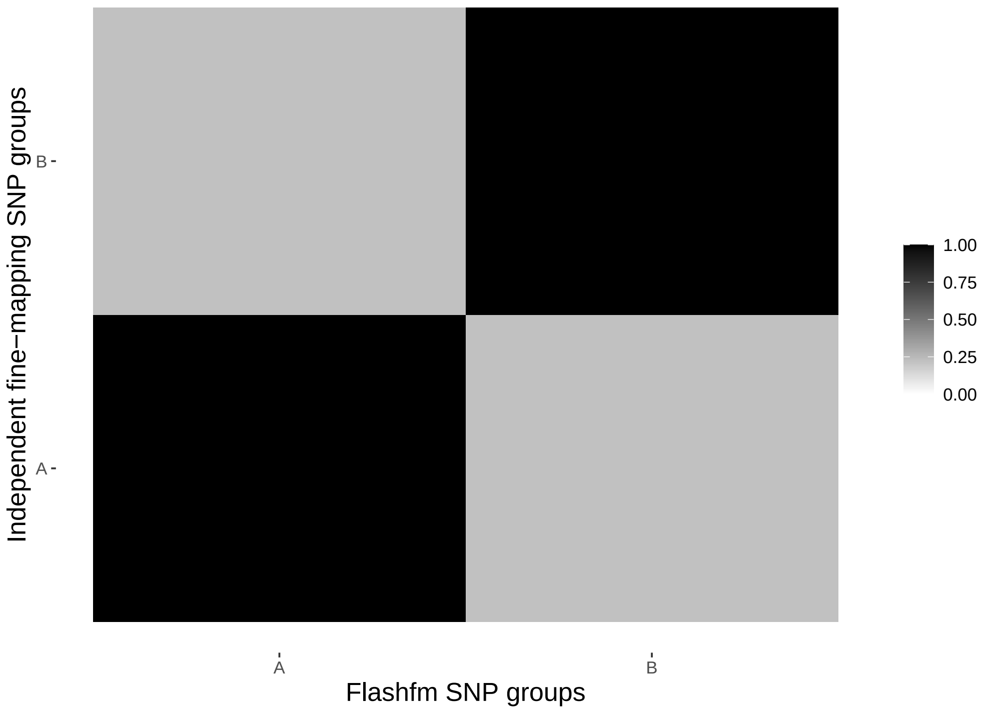

Supplementary Figure 9: Average LD between SNP groups constructed under independent fine-mapping (left) and Flashfm (right) for region 1:220212231-220479374.

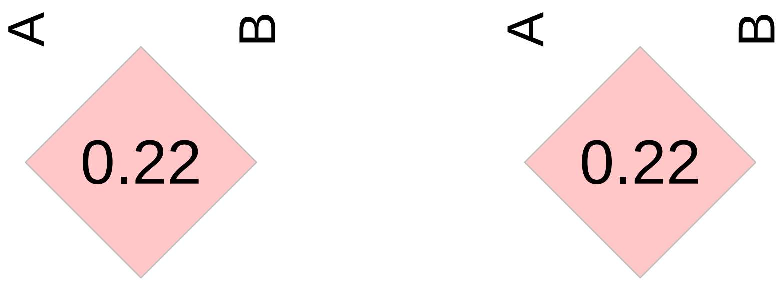

| | Stepwise | | Independent | | Flashfm | |
|---|---|---|---|---|---|---|
| Traits | SNP/Model | P-value | Model | PP | Model | PP |
| neut_count | 2:136485657/D | 1.05224e-11 | D | 0.861 | D | 0.866 |
| rbc | 2:135784852/A | 8.51555e-09 | A+B | 0.086 | A+B | 0.085 |

Supplementary Table 11: Fine-mapping results for 2:135758116-136787730 by independent fine-mapping and Flashfm. Stepwise results are also given.

| | SNP Group | | |
|---|---|---|---|
| | A | B | D |
| Independent | 1 | 5 | 45 |
| | 2:135784852 | 2:136353460 | 2:136493550 |
| Flashfm | 1 | 5 | 45 |
| | 2:135784852 | 2:136353460 | 2:136493550 |

Supplementary Table 12: SNP group sizes and representative variant for groups constructed by independent fine-mapping and by Flashfm for region 2:135758116-136787730.

Supplementary Figure 10: Average LD between SNP groups constructed under independent fine-mapping with those from flashfm for region 2:135758116-136787730.

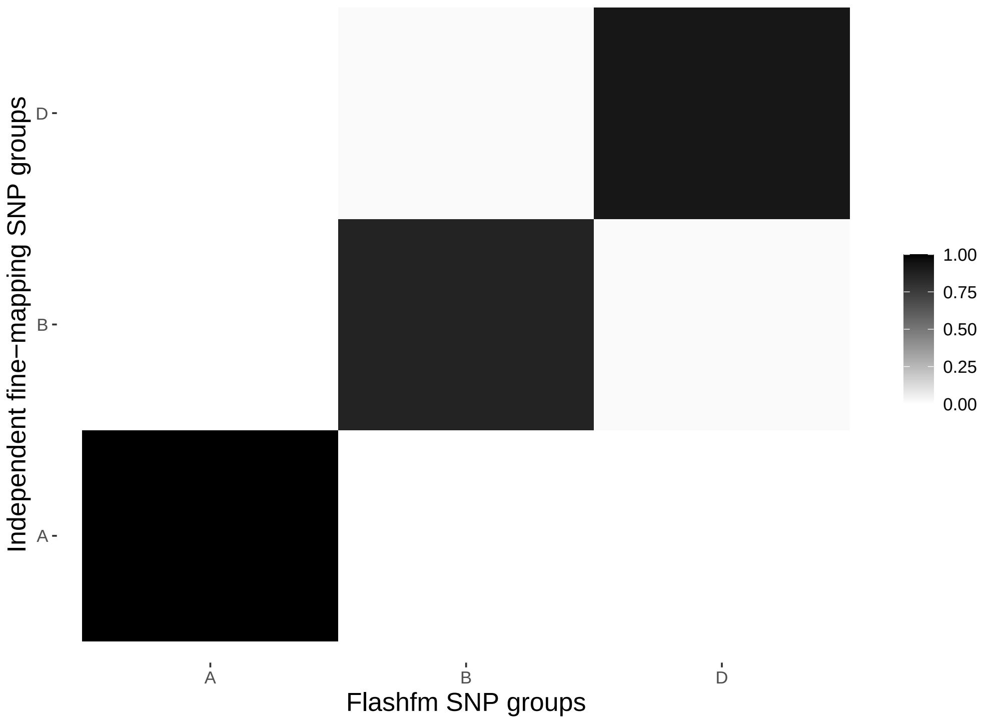

Supplementary Figure 11: Average LD between SNP groups constructed under independent fine-mapping (left) and Flashfm (right) for region 2:135758116-136787730.

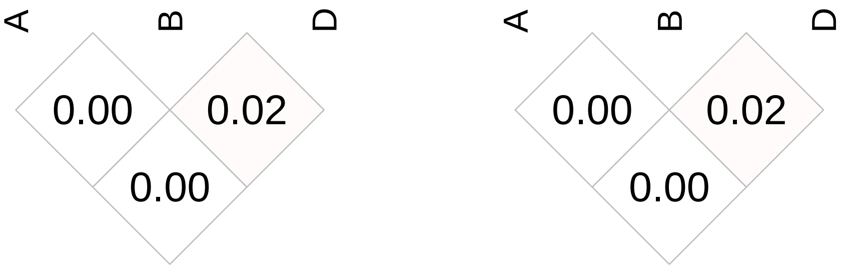

| | Stepwise | | Independent | | Flashfm | |
|---|---|---|---|---|---|---|
| Traits | SNP/Model | P-value | Model | PP | Model | PP |
| hba1c2 | 2:189377509/I | 6.51918e-10 | A+B+G+H+I+K | 0.622 | A+B+G+H+I+K | 0.595 |
| | 2:189593139/G | 8.20728e-07 | A+B+G+H+K | 0.214 | A+B+G+H+K | 0.230 |
| | – | – | – | – | A+B+G+H | 0.050 |
| weight | 2:189382844/D | 2.28506e-07 | D | 0.536 | D | 0.436 |
| | – | – | – | – | J | 0.131 |
| | – | – | – | – | D+J | 0.124 |
| gamma | 2:189185755/J | 3.11663e-07 | J | 0.893 | J | 0.672 |
| | – | – | – | – | B+J | 0.187 |

Supplementary Table 13: Fine-mapping results for 2:189071956-189635041 by independent fine-mapping and Flashfm. Stepwise results are also given.

| | SNP Group | | | | | |
|---|---|---|---|---|---|---|
| | A | B | D | G | H | I |
| Independent | 2 | 1 | 5 | 1 | 17 | 5 |
| | 2:189436024 | 2:189461890 | 2:189350336 | 2:189593139 | 2:189629716 | 2:189377509 |
| Flashfm | 2 | 1 | 5 | 1 | 17 | 4 |
| | 2:189436024 | 2:189461890 | 2:189350336 | 2:189593139 | 2:189629716 | 2:189413894 |
| | J | K | | | | |
| Independent | 20 | 15 | | | | |
| | 2:189194029 | 2:189443782 | | | | |
| Flashfm | 17 | 15 | | | | |
| | 2:189106677 | 2:189443782 | | | | |

Supplementary Table 14: SNP group sizes and representative variant for groups constructed by independent fine-mapping and by Flashfm for region 2:189071956-189635041.

Supplementary Figure 12: Average LD between SNP groups constructed under independent fine-mapping with those from flashfm for region 2:189071956-189635041.

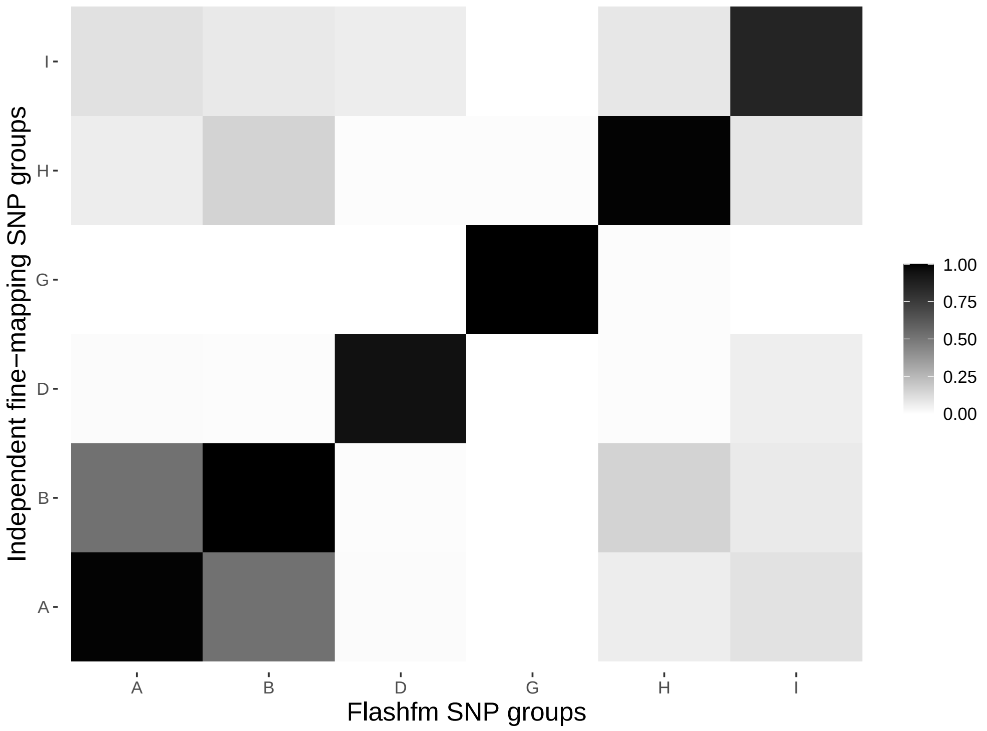

Supplementary Figure 13: Average LD between SNP groups constructed under independent fine-mapping (left) and Flashfm (right) for region 2:189071956-189635041.

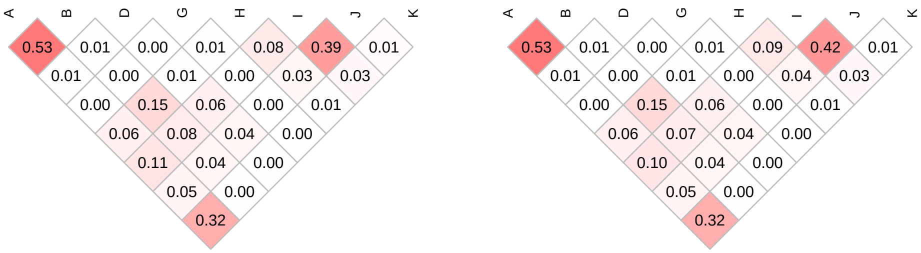

| | Stepwise | | Independent | | Flashfm | |
|---|---|---|---|---|---|---|
| Traits | SNP/Model | P-value | Model | PP | Model | PP |
| alkaline | 2:62760476/B | 5.9724e-08 | B | 0.657 | B | 0.731 |
| plt | 2:62742534/J | 4.9984e-07 | J | 0.462 | B+J | 0.623 |
| | – | – | – | – | J | 0.156 |
| | – | – | – | – | B | 0.078 |

Supplementary Table 15: Fine-mapping results for 2:62716187-62887884 by independent fine-mapping and Flashfm. Stepwise results are also given.

| | SNP Group | |
|---|---|---|
| | B | J |
| Independent | 8 | 1 |
| | 2:62781499 | 2:62742534 |
| Flashfm | 5 | 1 |
| | 2:62761855 | 2:62742534 |

Supplementary Table 16: SNP group sizes and representative variant for groups constructed by independent fine-mapping and by Flashfm for region 2:62716187-62887884.

Supplementary Figure 14: Average LD between SNP groups constructed under independent fine-mapping with those from flashfm for region 2:62716187-62887884.

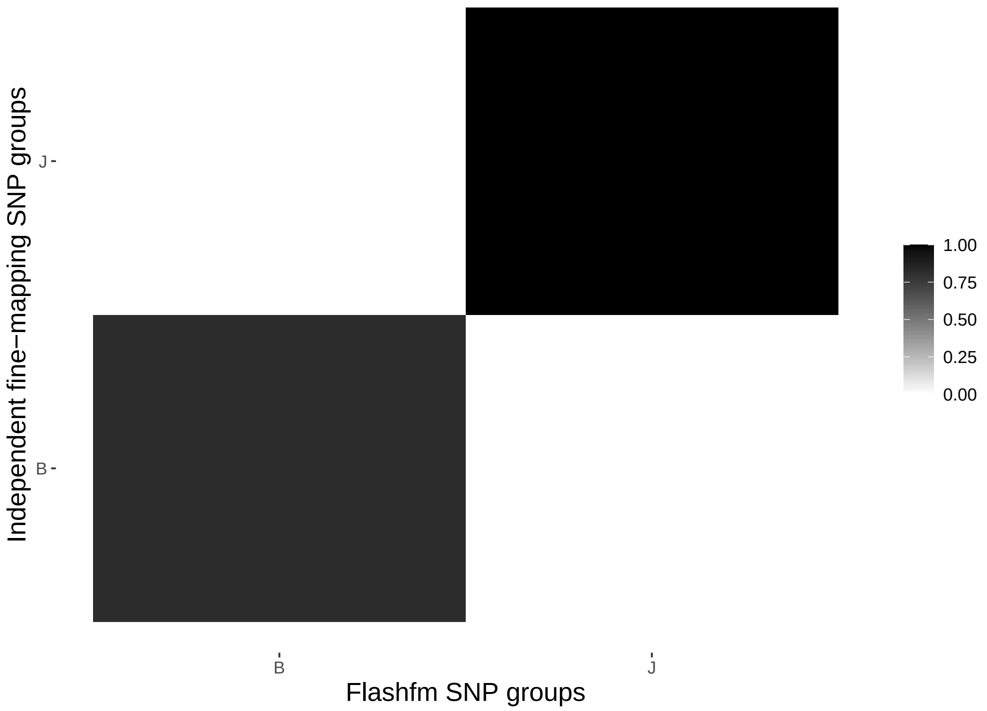

Supplementary Figure 15: Average LD between SNP groups constructed under independent fine-mapping (left) and Flashfm (right) for region 2:62716187-62887884.

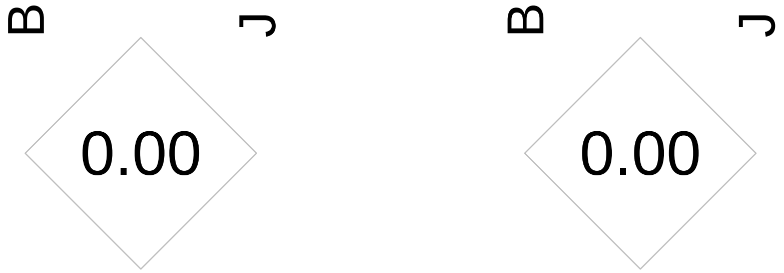

| | Stepwise | | Independent | | Flashfm | |
|---|---|---|---|---|---|---|
| Traits | SNP/Model | P-value | Model | PP | Model | PP |
| aspartate | 2:85387799/F | 2.62681e-07 | F | 0.66 | F | 0.676 |
| | – | – | – | – | E+F | 0.058 |
| rbc_dstr_width | 2:85382730/G | 7.46467e-07 | G | 0.318 | F+G | 0.325 |
| | – | – | E | 0.231 | G | 0.184 |
| | – | – | A+G | 0.119 | E | 0.142 |
| | – | – | A+E | 0.058 | E+F | 0.113 |
| | – | – | – | – | A+G | 0.054 |

Supplementary Table 17: Fine-mapping results for 2:85364419-85412152 by independent fine-mapping and Flashfm. Stepwise results are also given.

| | SNP Group | | | |
|---|---|---|---|---|
| | A | E | F | G |
| Independent | 13 | 21 | 24 | 2 |
| | 2:85400920 | 2:85379656 | 2:85410632 | 2:85382730 |
| Flashfm | 13 | 21 | 13 | 2 |
| | 2:85400920 | 2:85379656 | 2:85387799 | 2:85382730 |

Supplementary Table 18: SNP group sizes and representative variant for groups constructed by independent fine-mapping and by Flashfm for region 2:85364419-85412152.

Supplementary Figure 16: Average LD between SNP groups constructed under independent fine-mapping with those from flashfm for region 2:85364419-85412152.

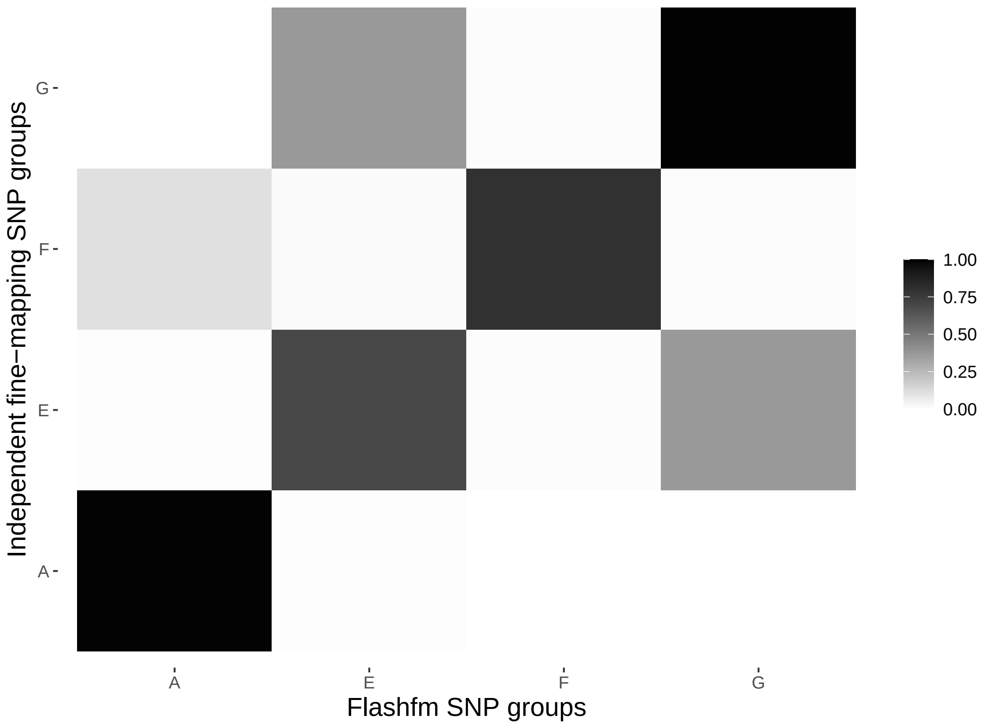

Supplementary Figure 17: Average LD between SNP groups constructed under independent fine-mapping (left) and Flashfm (right) for region 2:85364419-85412152.

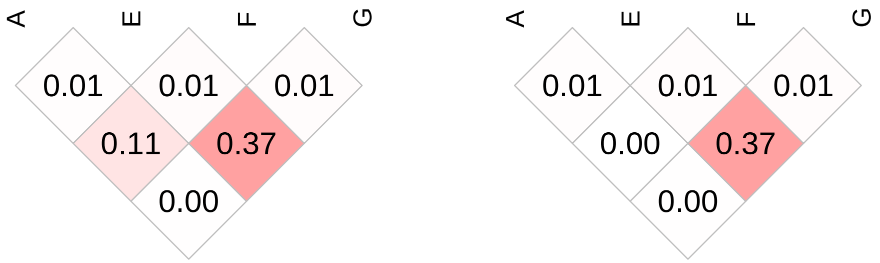

| | Stepwise | | Independent | | Flashfm | |
|---|---|---|---|---|---|---|
| Traits | SNP/Model | P-value | Model | PP | Model | PP |
| hct | 2:18682721/B | 6.1164e-07 | B | 0.547 | B | 0.511 |
| | – | – | B+C | 0.085 | A | 0.133 |
| | – | – | A | 0.075 | B+C | 0.097 |
| hgb | 2:18682721/B | 7.50734e-07 | B | 0.677 | B | 0.65 |
| | – | – | A | 0.068 | A | 0.127 |
| | – | – | B+F | 0.055 | – | – |

Supplementary Table 19: Fine-mapping results for 2:18651577-18703005 by independent fine-mapping and Flashfm. Stepwise results are also given.

| | SNP Group | | | |
|---|---|---|---|---|
| | A | B | C | F |
| Independent | 5 | 11 | 2 | 13 |
| | 2:18703005 | 2:18675213 | 2:18701221 | 2:18691613 |
| Flashfm | 5 | 11 | 2 | 13 |
| | 2:18703005 | 2:18675213 | 2:18701221 | 2:18691613 |

Supplementary Table 20: SNP group sizes and representative variant for groups constructed by independent fine-mapping and by Flashfm for region 2:18651577-18703005.

Supplementary Figure 18: Average LD between SNP groups constructed under independent fine-mapping with those from flashfm for region 2:18651577-18703005.

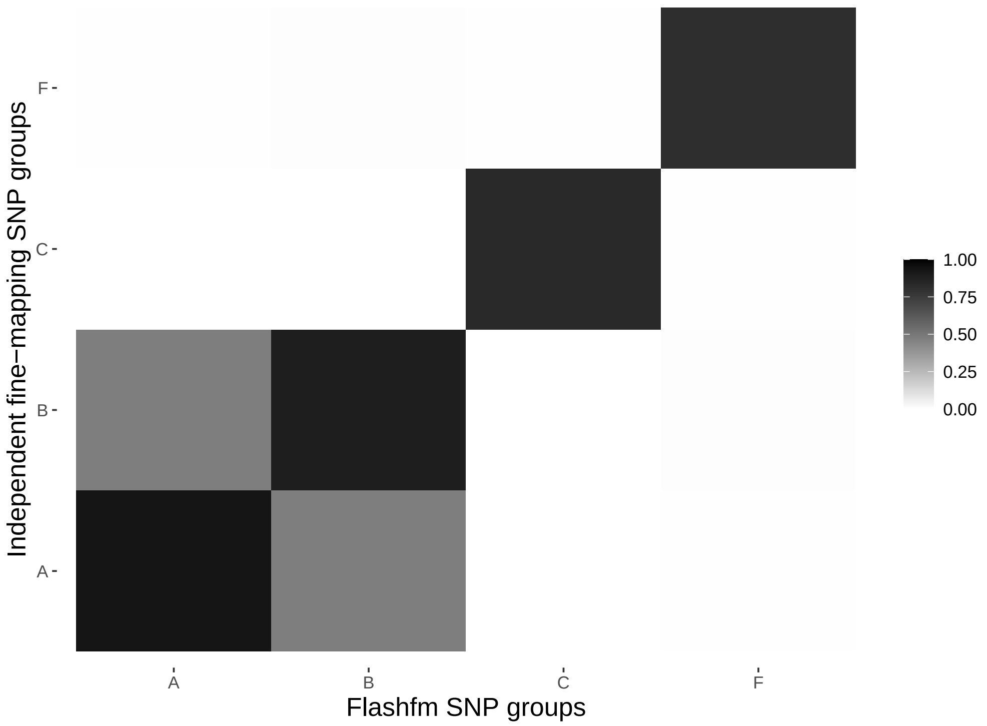

Supplementary Figure 19: Average LD between SNP groups constructed under independent fine-mapping (left) and Flashfm (right) for region 2:18651577-18703005.

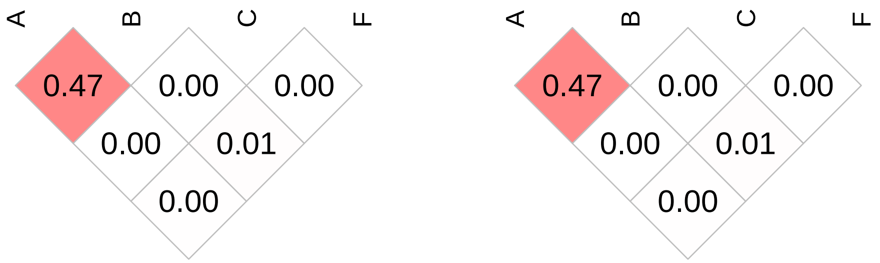

| | Stepwise | | Independent | | Flashfm | |
|---|---|---|---|---|---|---|
| Traits | SNP/Model | P-value | Model | PP | Model | PP |
| hgb | 3:48547711/A | 1.02557e-07 | B | 0.613 | B.1 | 0.623 |
| | – | – | A | 0.222 | A | 0.266 |
| hct | 3:48547711/A | 1.81636e-07 | A | 0.468 | A | 0.550 |
| | – | – | B | 0.323 | B.1 | 0.290 |

Supplementary Table 21: Fine-mapping results for 3:48378500-48938297 by independent fine-mapping and Flashfm. Stepwise results are also given.

| | SNP Group | | |
|---|---|---|---|
| | A | B | B.1 |
| Independent | 8 | 40 | 0 |
| | 3:48721571 | 3:48626557 | – |
| Flashfm | 8 | 0 | 29 |
| | 3:48721571 | – | 3:48626557 |

Supplementary Table 22: SNP group sizes and representative variant for groups constructed by independent fine-mapping and by Flashfm for region 3:48378500-48938297.

Supplementary Figure 20: Average LD between SNP groups constructed under independent fine-mapping with those from flashfm for region 3:48378500-48938297.

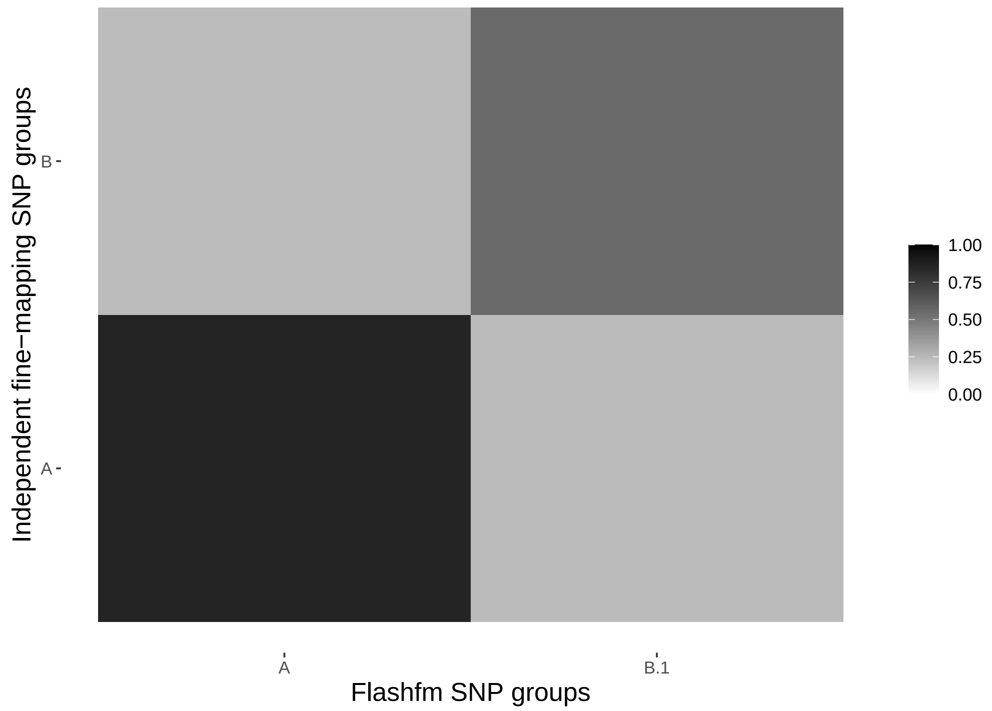

Supplementary Figure 21: Average LD between SNP groups constructed under independent fine-mapping (left) and Flashfm (right) for region 3:48378500-48938297.

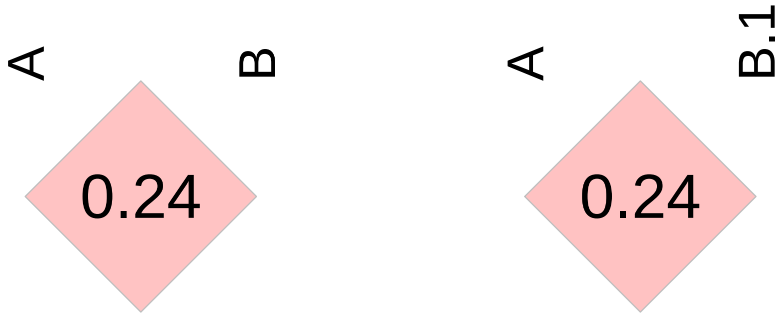

| | Stepwise | | Independent | | Flashfm | |
|---|---|---|---|---|---|---|
| Traits | SNP/Model | P-value | Model | PP | Model | PP |
| SBP | 3:135663939/A | 4.25088e-07 | A | 0.861 | A.2 | 0.762 |
| | – | – | A+C | 0.054 | A.1 | 0.165 |
| DBP | 3:135673880/A | 6.06157e-07 | A+B | 0.643 | A.2+B | 0.555 |
| | – | – | A | 0.335 | A.2 | 0.433 |

Supplementary Table 23: Fine-mapping results for 3:135548414-135986942 by independent fine-mapping and Flashfm. Stepwise results are also given.

| | SNP Group | | | | |
|---|---|---|---|---|---|
| | A | B | C | A.1 | A.2 |
| Independent | 98 | 54 | 21 | 0 | 0 |
| | 3:135650060 | 3:135589895 | 3:135692426 | – | – |
| Flashfm | 0 | 54 | 14 | 2 | 36 |
| | – | 3:135589895 | 3:135658784 | 3:135650060 | 3:135676132 |

Supplementary Table 24: SNP group sizes and representative variant for groups constructed by independent fine-mapping and by Flashfm for region 3:135548414-135986942.

Supplementary Figure 22: Average LD between SNP groups constructed under independent fine-mapping with those from flashfm for region 3:135548414-135986942.

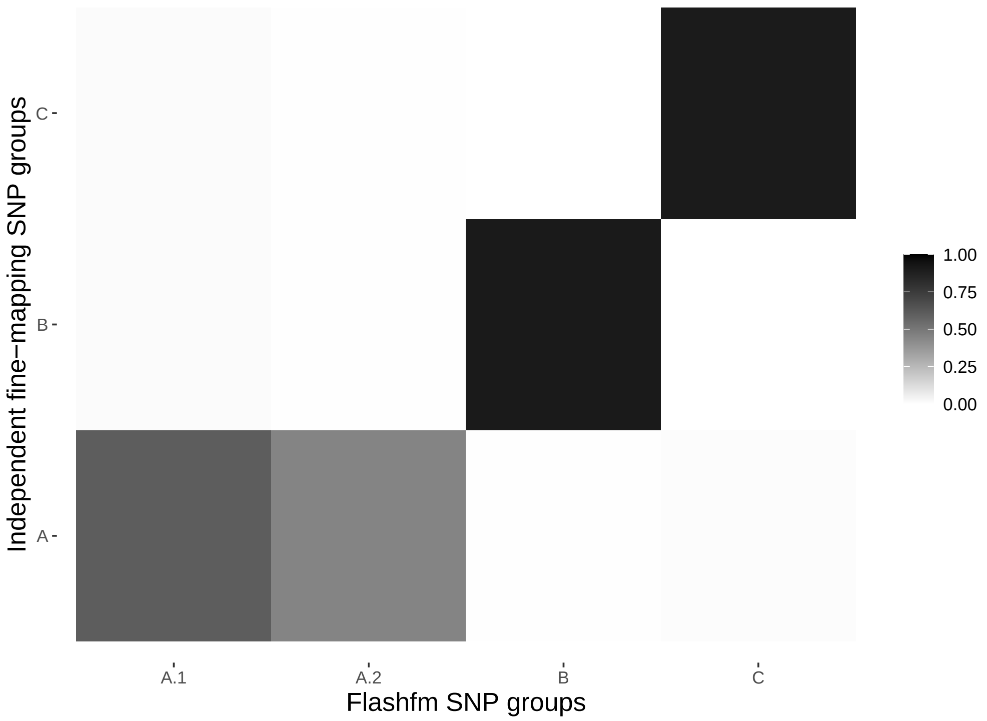

Supplementary Figure 23: Average LD between SNP groups constructed under independent fine-mapping (left) and Flashfm (right) for region 3:135548414-135986942.

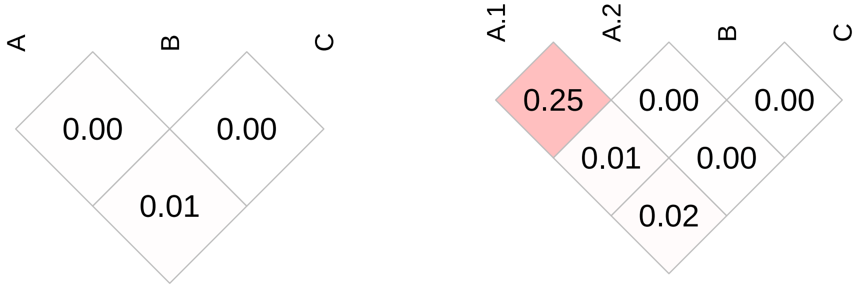

| | Stepwise | | Independent | | Flashfm | |
|---|---|---|---|---|---|---|
| Traits | SNP/Model | P-value | Model | PP | Model | PP |
| lymp_count | 4:53095312/B | 5.56795e-08 | B | 0.789 | B | 0.832 |
| weight | 4:53083377/A | 4.82996e-07 | A | 0.665 | A | 0.718 |

Supplementary Table 25: Fine-mapping results for 4:49648062-53304218 by independent fine-mapping and Flashfm. Stepwise results are also given.

| | SNP Group | |
|---|---|---|
| | A | B |
| Independent | 1 | 2 |
| | 4:53083377 | 4:53060664 |
| Flashfm | 1 | 2 |
| | 4:53083377 | 4:53060664 |

Supplementary Table 26: SNP group sizes and representative variant for groups constructed by independent fine-mapping and by Flashfm for region 4:49648062-53304218.

Supplementary Figure 24: Average LD between SNP groups constructed under independent fine-mapping with those from flashfm for region 4:49648062-53304218.

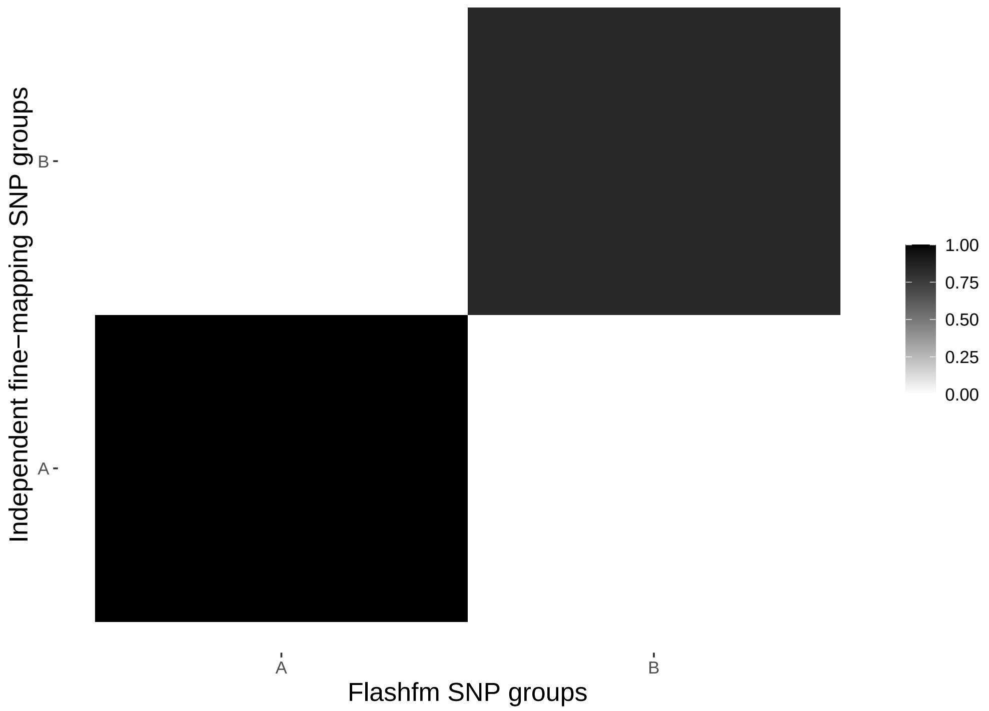

Supplementary Figure 25: Average LD between SNP groups constructed under independent fine-mapping (left) and Flashfm (right) for region 4:49648062-53304218.

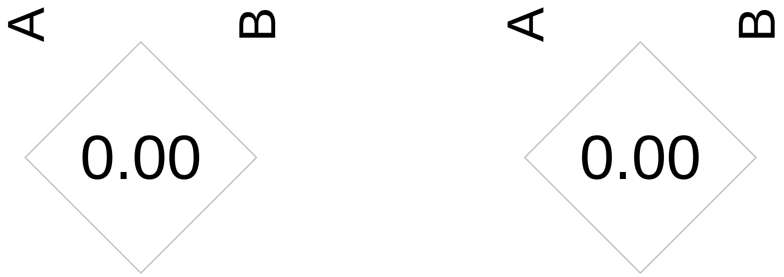

| | Stepwise | | Independent | | Flashfm | |
|---|---|---|---|---|---|---|
| Traits | SNP/Model | P-value | Model | PP | Model | PP |
| lymp_count | 4:19765235/G | 7.2126e-08 | G | 0.403 | G | 0.49 |
| | – | – | E+G | 0.213 | E+G | 0.184 |
| | – | – | D+G | 0.063 | D+G | 0.056 |
| wbc | 4:19765235/G | 1.62137e-07 | G | 0.714 | G | 0.794 |
| mono_count | 4:19765235/G | 7.48651e-07 | G | 0.369 | G | 0.532 |
| | – | – | C+G | 0.088 | C+G | 0.092 |
| | – | – | C | 0.061 | E+G | 0.055 |
| | – | – | E+G | 0.052 | – | – |

Supplementary Table 27: Fine-mapping results for 4:19443144-19852689 by independent fine-mapping and Flashfm. Stepwise results are also given.

| | SNP Group | | | |
|---|---|---|---|---|
| | C | D | E | G |
| Independent | 23 | 4 | 59 | 1 |
| | 4:19598292 | 4:19532959 | 4:19638561 | 4:19765235 |
| Flashfm | 23 | 4 | 52 | 1 |
| | 4:19598292 | 4:19532959 | 4:19638561 | 4:19765235 |

Supplementary Table 28: SNP group sizes and representative variant for groups constructed by independent fine-mapping and by Flashfm for region 4:19443144-19852689.

Supplementary Figure 26: Average LD between SNP groups constructed under independent fine-mapping with those from flashfm for region 4:19443144-19852689.

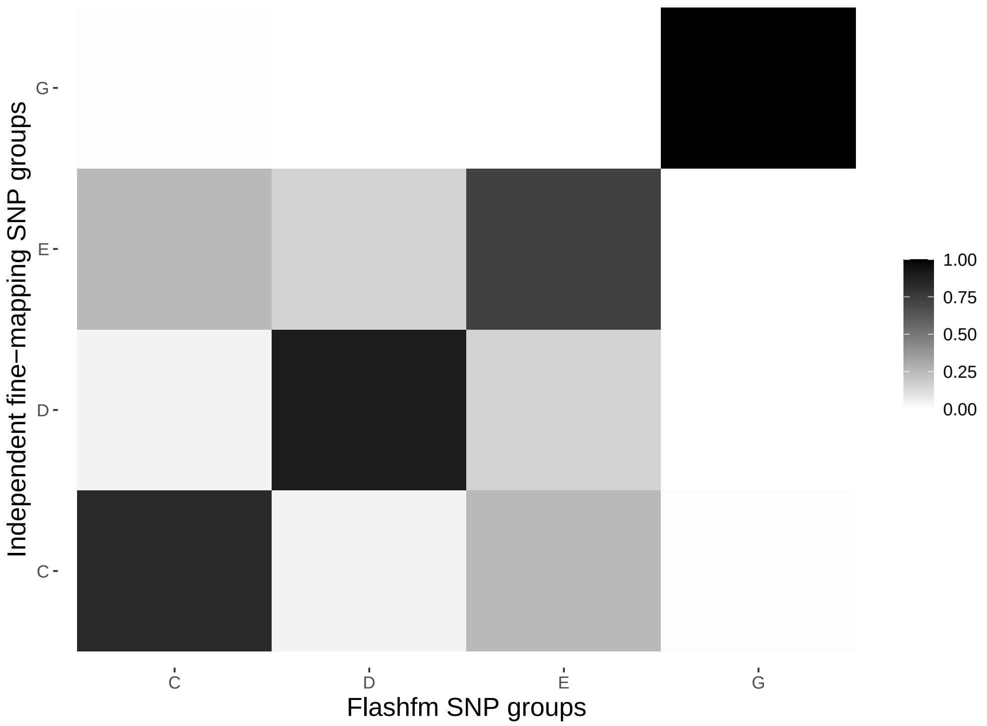

Supplementary Figure 27: Average LD between SNP groups constructed under independent fine-mapping (left) and Flashfm (right) for region 4:19443144-19852689.

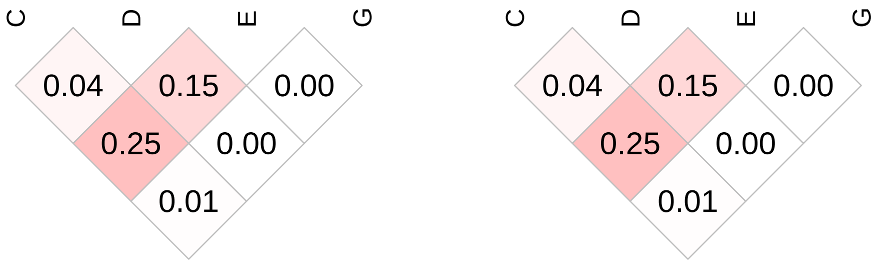

| | Stepwise | | Independent | | Flashfm | |
|---|---|---|---|---|---|---|
| Traits | SNP/Model | P-value | Model | PP | Model | PP |
| cholesterol | 5:156378584/A | 4.0245e-10 | A | 0.328 | A | 0.494 |
| | – | – | A+G | 0.288 | A+G | 0.211 |
| | – | – | A+E | 0.177 | A+E | 0.129 |
| lowdlipo | 5:156309938/A | 2.93049e-09 | A+G | 0.491 | A+G | 0.445 |
| | – | – | A | 0.136 | A | 0.253 |
| | – | – | A+E | 0.094 | A+E | 0.085 |

Supplementary Table 29: Fine-mapping results for 5:156281094-156517240 by independent fine-mapping and Flashfm. Stepwise results are also given.

| | SNP Group | | |
|---|---|---|---|
| | A | E | G |
| Independent | 5 | 37 | 47 |
| | 5:156309938 | 5:156482884 | 5:156443079 |
| Flashfm | 5 | 37 | 47 |
| | 5:156309938 | 5:156482884 | 5:156443079 |

Supplementary Table 30: SNP group sizes and representative variant for groups constructed by independent fine-mapping and by Flashfm for region 5:156281094-156517240.

Supplementary Figure 28: Average LD between SNP groups constructed under independent fine-mapping with those from flashfm for region 5:156281094-156517240.

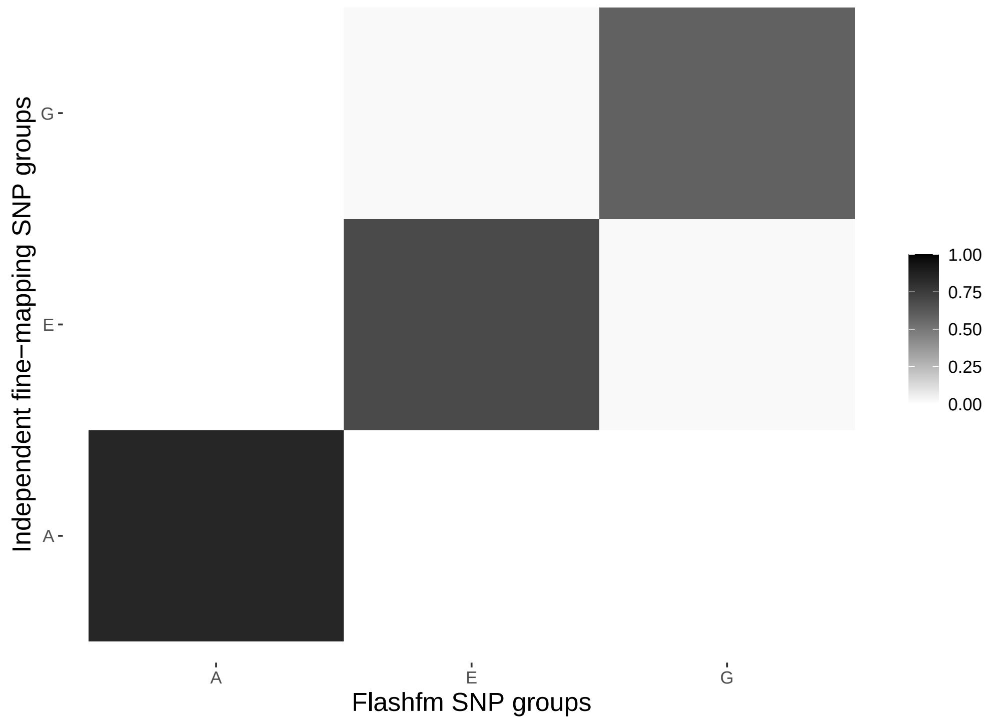

Supplementary Figure 29: Average LD between SNP groups constructed under independent fine-mapping (left) and Flashfm (right) for region 5:156281094-156517240.

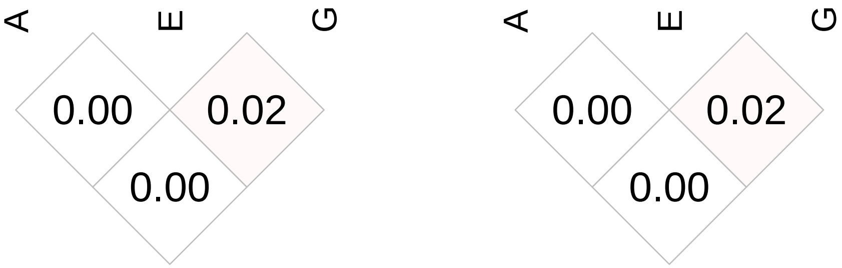

| | Stepwise | | Independent | | Flashfm | |
|---|---|---|---|---|---|---|
| Traits | SNP/Model | P-value | Model | PP | Model | PP |
| cholesterol | 5:156309938/B | 1.46167e-09 | B | 0.684 | B | 0.795 |
| | – | – | B+E | 0.051 | – | – |
| lowdlipo | 5:156309938/B | 2.93049e-09 | B | 0.436 | B | 0.601 |
| | – | – | B+E | 0.157 | B+E | 0.106 |
| | – | – | B+F | 0.062 | – | – |

Supplementary Table 31: Fine-mapping results for 5:156067961-156345403 by independent fine-mapping and Flashfm. Stepwise results are also given.

| | SNP Group | | |
|---|---|---|---|
| | B | E | F |
| Independent | 4 | 27 | 20 |
| | 5:156142225 | 5:156294885 | 5:156226108 |
| Flashfm | 4 | 27 | 17 |
| | 5:156142225 | 5:156294885 | 5:156257190 |

Supplementary Table 32: SNP group sizes and representative variant for groups constructed by independent fine-mapping and by Flashfm for region 5:156067961-156345403.

Supplementary Figure 30: Average LD between SNP groups constructed under independent fine-mapping with those from flashfm for region 5:156067961-156345403.

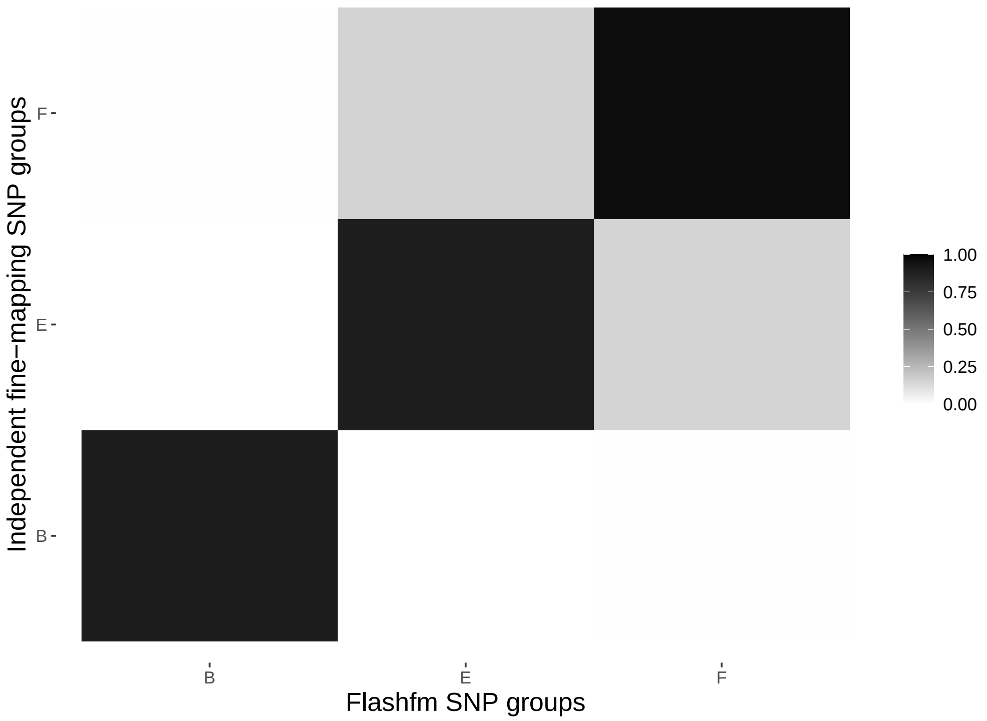

Supplementary Figure 31: Average LD between SNP groups constructed under independent fine-mapping (left) and Flashfm (right) for region 5:156067961-156345403.

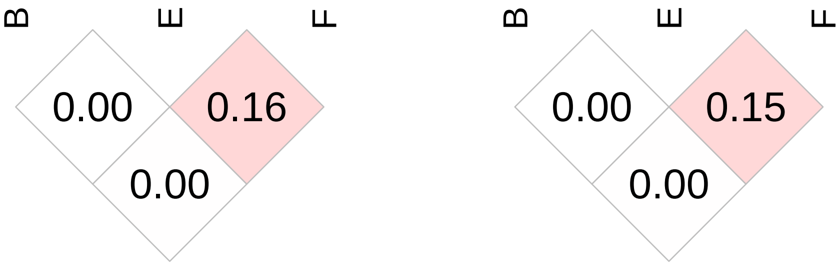

| | Stepwise | | Independent | | Flashfm | |
|---|---|---|---|---|---|---|
| Traits | SNP/Model | P-value | Model | PP | Model | PP |
| cholesterol | 5:155883245/C | 1.08914e-08 | C | 0.677 | C | 0.521 |
| | – | – | A | 0.111 | A | 0.307 |
| lowdlipo | 5:155883245/C | 4.42672e-08 | C | 0.732 | C | 0.677 |
| | – | – | – | – | B | 0.081 |

Supplementary Table 33: Fine-mapping results for 5:155765598-155972507 by independent fine-mapping and Flashfm. Stepwise results are also given.

| | SNP Group | | |
|---|---|---|---|
| | A | B | C |
| Independent | 1 | 1 | 5 |
| | 5:155960873 | 5:155972507 | 5:155916168 |
| Flashfm | 1 | 1 | 5 |
| | 5:155960873 | 5:155972507 | 5:155916168 |

Supplementary Table 34: SNP group sizes and representative variant for groups constructed by independent fine-mapping and by Flashfm for region 5:155765598-155972507.

Supplementary Figure 32: Average LD between SNP groups constructed under independent fine-mapping with those from flashfm for region 5:155765598-155972507.

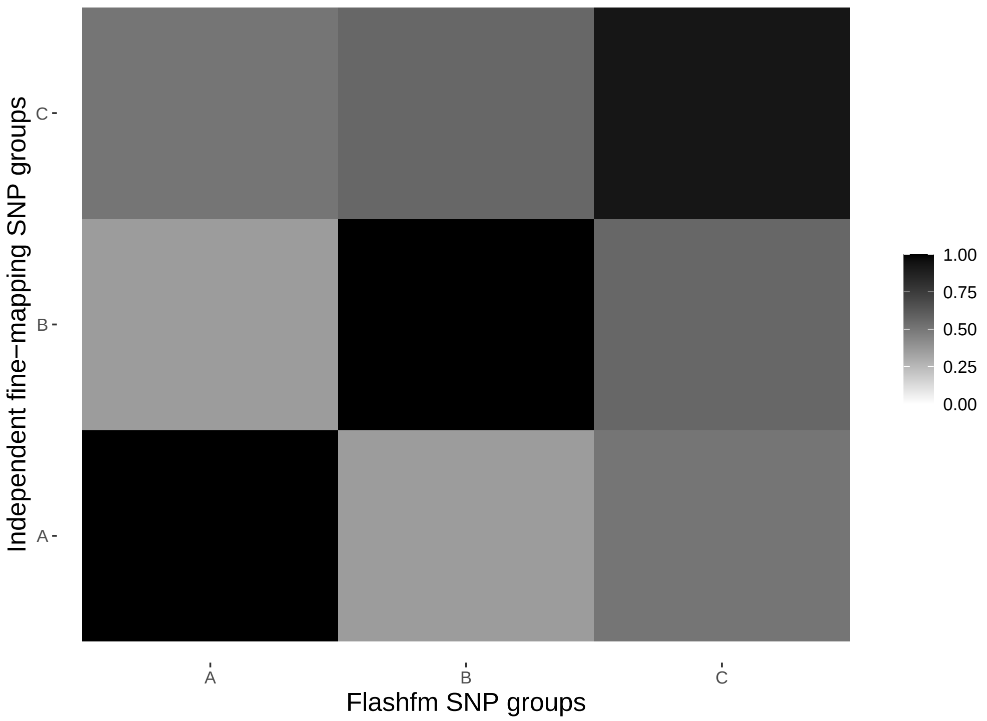

Supplementary Figure 33: Average LD between SNP groups constructed under independent fine-mapping (left) and Flashfm (right) for region 5:155765598-155972507.

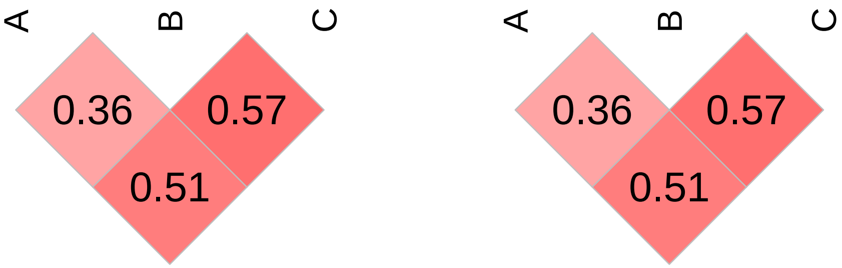

| | Stepwise | | Independent | | Flashfm | |
|---|---|---|---|---|---|---|
| Traits | SNP/Model | P-value | Model | PP | Model | PP |
| hct | 5:133872879/A | 2.41027e-08 | A | 0.551 | A | 0.657 |
| | – | – | A+C | 0.078 | A+C | 0.051 |
| hgb | 5:133881621/A | 1.31749e-07 | A | 0.574 | A | 0.676 |

Supplementary Table 35: Fine-mapping results for 5:133815324-133907415 by independent fine-mapping and Flashfm. Stepwise results are also given.

| | SNP Group | |
|---|---|---|
| | A | C |
| Independent | 3 | 8 |
| | 5:133860151 | 5:133836607 |
| Flashfm | 3 | 8 |
| | 5:133860151 | 5:133836607 |

Supplementary Table 36: SNP group sizes and representative variant for groups constructed by independent fine-mapping and by Flashfm for region 5:133815324-133907415.

Supplementary Figure 34: Average LD between SNP groups constructed under independent fine-mapping with those from flashfm for region 5:133815324-133907415.

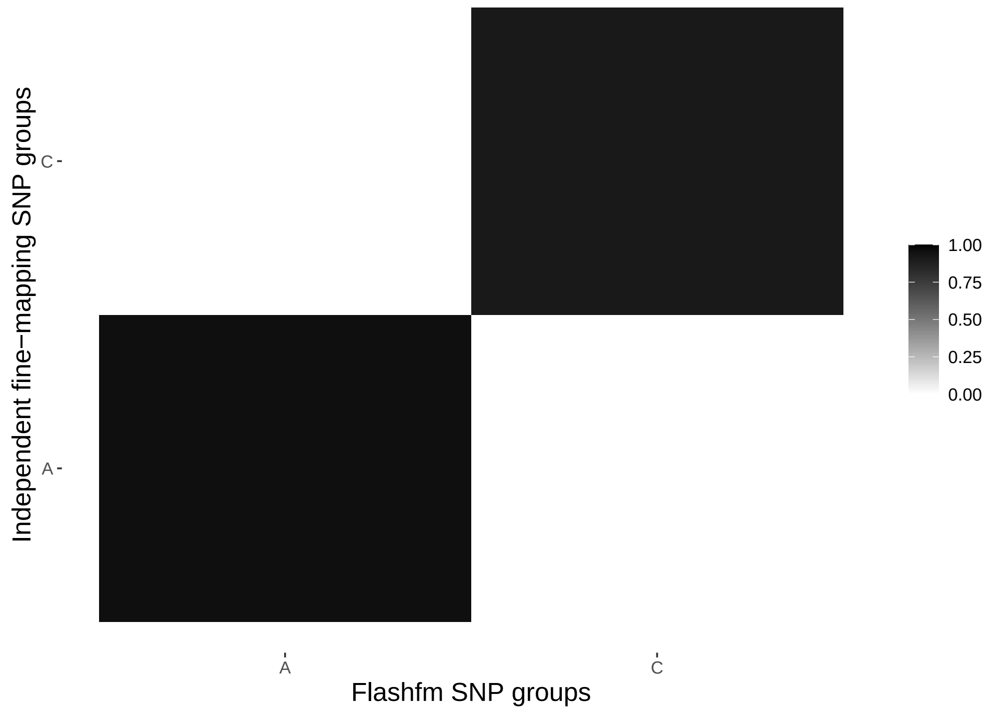

Supplementary Figure 35: Average LD between SNP groups constructed under independent fine-mapping (left) and Flashfm (right) for region 5:133815324-133907415.

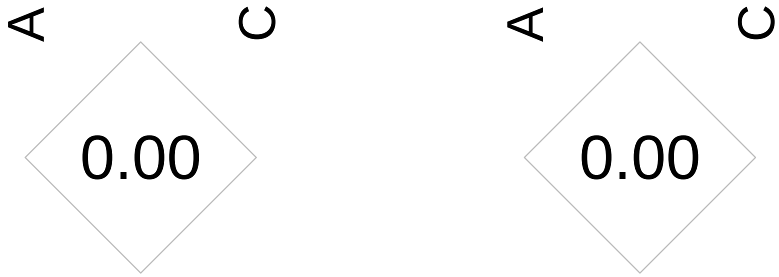

| | Stepwise | | Independent | | Flashfm | |
|---|---|---|---|---|---|---|
| Traits | SNP/Model | P-value | Model | PP | Model | PP |
| cholesterol | 5:155781587/B | 5.89497e-08 | B | 0.851 | B | 0.837 |
| lowdlipo | 5:155781587/B | 5.3179e-08 | B | 0.816 | B | 0.801 |

Supplementary Table 37: Fine-mapping results for 5:155573180-155781587 by independent fine-mapping and Flashfm. Stepwise results are also given.

| | SNP Group |
|---|---|
| | B |
| Independent | 11 |
| | 5:155781587 |
| Flashfm | 11 |
| | 5:155781587 |

Supplementary Table 38: SNP group sizes and representative variant for groups constructed by independent fine-mapping and by Flashfm for region 5:155573180-155781587.

Supplementary Figure 36: Venn diagram of SNP group constructed under indepenedent fine-mapping compared to that under flashfm for region 5:155573180-155781587. This shows that the flashfm group is a subset of, or the same as, the group form independent fine-mapping.

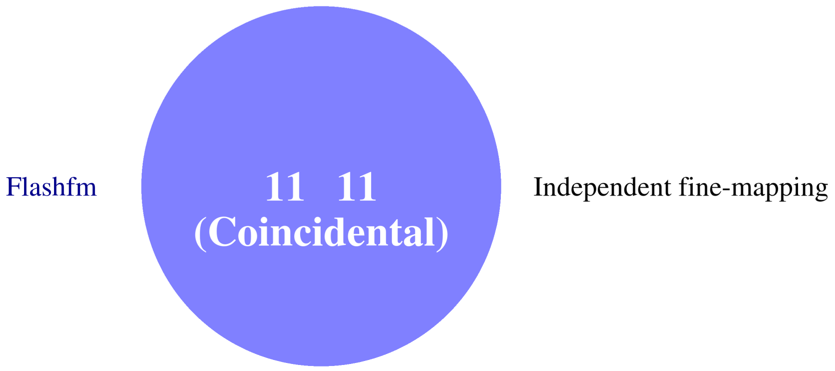

| | Stepwise | | Independent | | Flashfm | |
|---|---|---|---|---|---|---|
| Traits | SNP/Model | P-value | Model | PP | Model | PP |
| hba1c2 | 6:32609126/B | 4.1838e-08 | B+C+I | 0.34 | B+C+I+K | 0.648 |
| | 6:32601902/C | 7.53393e-07 | B+C | 0.182 | B+C+K | 0.147 |
| | 6:32665913/I | 1.38104e-07 | B | 0.113 | B+C+I | 0.059 |
| | – | – | – | – | B+C | 0.055 |
| aspartate | 6:32583830/chr6_32583830 | 1.46603e-07 | A+D+E+F+K+L | 0.516 | A+C+D+E+G+K | 0.447 |
| | – | – | A+D+E+F+K | 0.128 | A+D+E+F+K+L | 0.215 |
| | – | – | – | – | A+D+E+F+K | 0.082 |

Supplementary Table 39: Fine-mapping results for 6:32559710-32671044 by independent fine-mapping and Flashfm. Stepwise results are also given.

| | SNP Group | | | | | | |
|---|---|---|---|---|---|---|---|
| | A | B | C | D | E | F | G |
| Independent | 21<br>6:32570281 | 17<br>6:32650409 | 1<br>6:32601902 | 1<br>6:32561422 | 1<br>6:32573910 | 18<br>6:32667458 | 1<br>6:32597435 |
| Flashfm | 18<br>6:32568522 | 12<br>6:32628306 | 1<br>6:32601902 | 1<br>6:32561422 | 1<br>6:32573910 | 11<br>6:32670145 | 1<br>6:32597435 |
| | I | K | L | | | | |
| Independent | 1<br>6:32665913 | 1<br>6:32579904 | 1<br>6:32568521 | | | | |
| Flashfm | 1<br>6:32665913 | 1<br>6:32579904 | 1<br>6:32568521 | | | | |

Supplementary Table 40: SNP group sizes and representative variant for groups constructed by independent fine-mapping and by Flashfm for region 6:32559710-32671044.

Supplementary Figure 37: Average LD between SNP groups constructed under independent fine-mapping with those from flashfm for region 6:32559710-32671044.

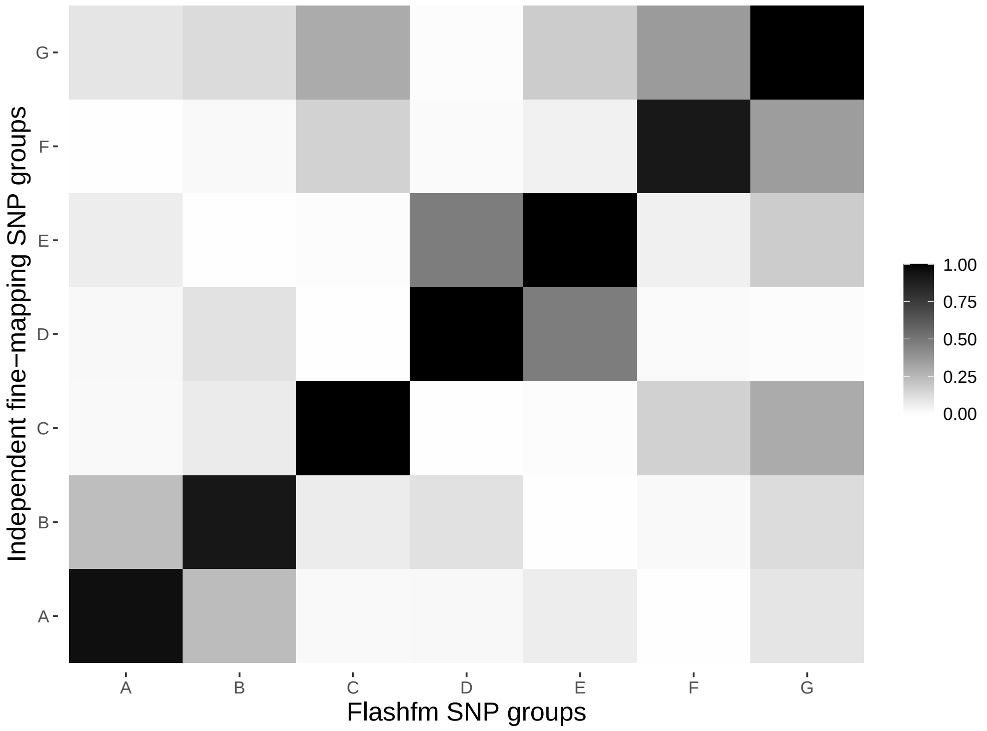

Supplementary Figure 38: Average LD between SNP groups constructed under independent fine-mapping (left) and Flashfm (right) for region 6:32559710-32671044.

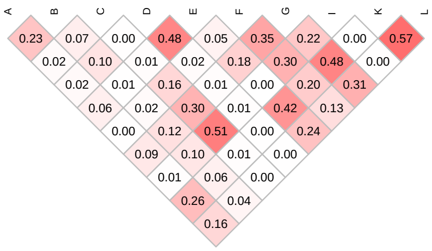


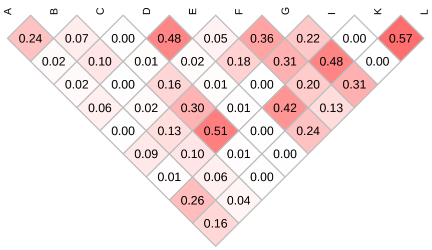

| | Stepwise | | Independent | | Flashfm | |
|---|---|---|---|---|---|---|
| Traits | SNP/Model | P-value | Model | PP | Model | PP |
| mch | 6:12369067/B | 6.21894e-08 | B | 0.352 | B | 0.513 |
| | – | – | B+I | 0.148 | B+I | 0.110 |
| mcv | 6:12369067/B | 6.72028e-07 | B | 0.326 | B | 0.451 |
| | – | – | B+I | 0.122 | L | 0.091 |
| | – | – | L | 0.063 | B+I | 0.086 |

Supplementary Table 41: Fine-mapping results for 6:12367517-12394380 by independent fine-mapping and Flashfm. Stepwise results are also given.

| | SNP Group | | |
|---|---|---|---|
| | B | I | L |
| Independent | 1 | 10 | 1 |
| | 6:12369067 | 6:12390604 | 6:12368852 |
| Flashfm | 1 | 10 | 1 |
| | 6:12369067 | 6:12390604 | 6:12368852 |

Supplementary Table 42: SNP group sizes and representative variant for groups constructed by independent fine-mapping and by Flashfm for region 6:12367517-12394380.

Supplementary Figure 39: Average LD between SNP groups constructed under independent fine-mapping with those from flashfm for region 6:12367517-12394380.

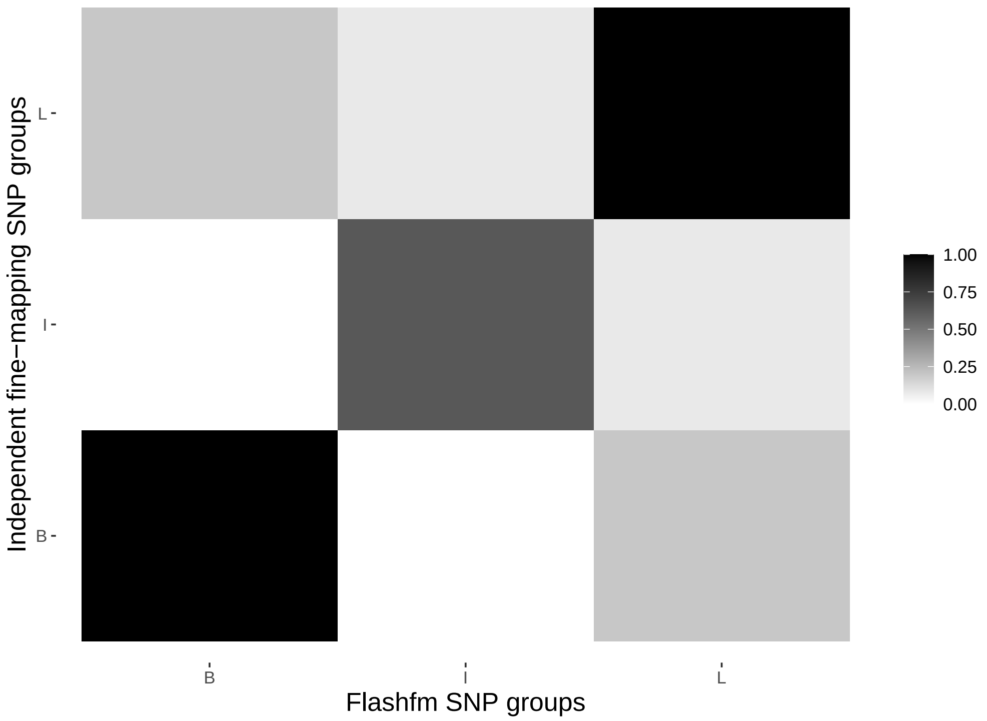

Supplementary Figure 40: Average LD between SNP groups constructed under independent fine-mapping (left) and Flashfm (right) for region 6:12367517-12394380.

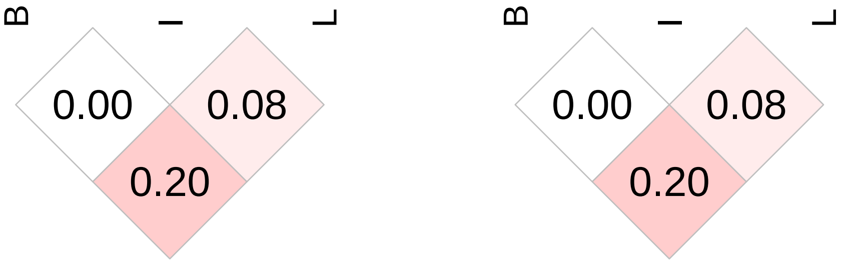

| | Stepwise | | Independent | | Flashfm | |
|---|---|---|---|---|---|---|
| Traits | SNP/Model | P-value | Model | PP | Model | PP |
| mcv | 6:167913072/A | 2.0766e-07 | A | 0.618 | A | 0.708 |
| mch | 6:167913072/A | 2.30335e-07 | A | 0.651 | A | 0.738 |

Supplementary Table 43: Fine-mapping results for 6:167874267-168050957 by independent fine-mapping and Flashfm. Stepwise results are also given.

| | SNP Group |
|---|---|
| | A |
| Independent | 3 |
| | 6:167919770 |
| Flashfm | 3 |
| | 6:167919770 |

Supplementary Table 44: SNP group sizes and representative variant for groups constructed by independent fine-mapping and by Flashfm for region 6:167874267-168050957.

Supplementary Figure 41: Venn diagram of SNP group constructed under indepenedent fine-mapping compared to that under flashfm for region 6:167874267-168050957. This shows that the flashfm group is a subset of, or the same as, the group form independent fine-mapping.

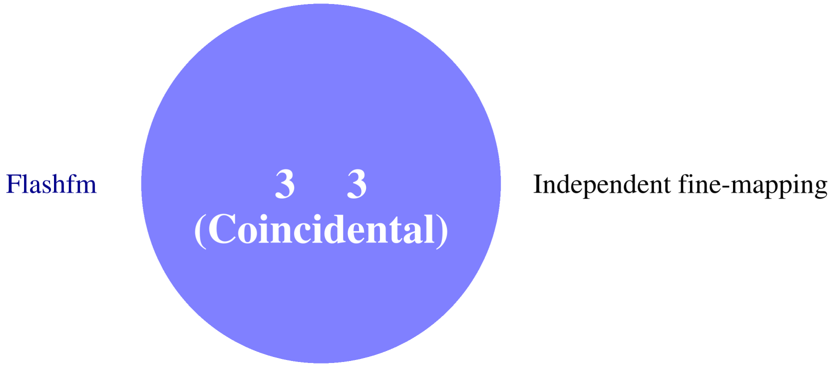

| | Stepwise | | Independent | | Flashfm | |
|---|---|---|---|---|---|---|
| Traits | SNP/Model | P-value | Model | PP | Model | PP |
| hip_circum | 8:6357093/A | 5.68796e-08 | A | 0.792 | A | 0.773 |
| weight | 8:6357093/A | 3.30175e-07 | A | 0.729 | A | 0.753 |

Supplementary Table 45: Fine-mapping results for 8:6353108-6394179 by independent fine-mapping and Flashfm. Stepwise results are also given.

| | SNP Group |
|---|---|
| | A |
| Independent | 18 |
| | 8:6357093 |
| Flashfm | 15 |
| | 8:6357093 |

Supplementary Table 46: SNP group sizes and representative variant for groups constructed by independent fine-mapping and by Flashfm for region 8:6353108-6394179.

Supplementary Figure 42: Venn diagram of SNP group constructed under indepenedent fine-mapping compared to that under flashfm for region 8:6353108-6394179. This shows that the flashfm group is a subset of, or the same as, the group form independent fine-mapping.

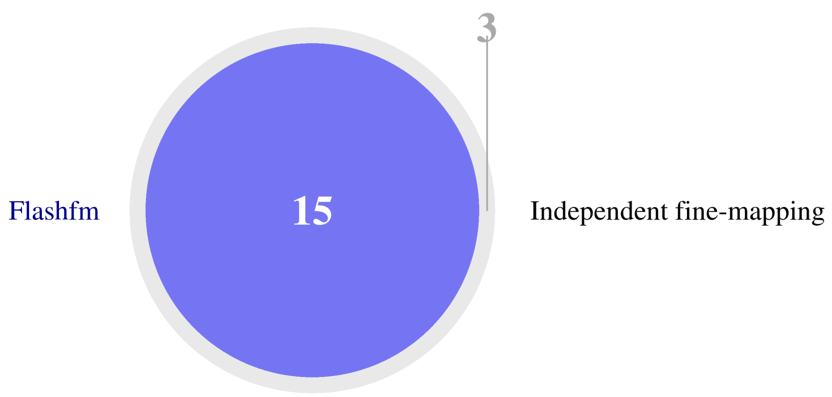

| | Stepwise | | Independent | | Flashfm | |
|---|---|---|---|---|---|---|
| Traits | SNP/Model | P-value | Model | PP | Model | PP |
| rbc_dstr_width | 11:5248232/A | 1.26439e-16 | A | 0.667 | A | 0.752 |
| | – | – | A+C | 0.087 | A+C | 0.07 |
| bilirubin | 11:5243936/A | 3.95972e-07 | A | 0.517 | A | 0.742 |
| | – | – | B | 0.192 | – | – |
| | – | – | A+B | 0.055 | – | – |

Supplementary Table 47: Fine-mapping results for 11:5211496-5252794 by independent fine-mapping and Flashfm. Stepwise results are also given.

| | SNP Group | | |
|---|---|---|---|
| | A | B | C |
| Independent | 4 | 12 | 1 |
| | 11:5252794 | 11:5236954 | 11:5230135 |
| Flashfm | 4 | 5 | 1 |
| | 11:5252794 | 11:5236954 | 11:5230135 |

Supplementary Table 48: SNP group sizes and representative variant for groups constructed by independent fine-mapping and by Flashfm for region 11:5211496-5252794.

Supplementary Figure 43: Average LD between SNP groups constructed under independent fine-mapping with those from flashfm for region 11:5211496-5252794.

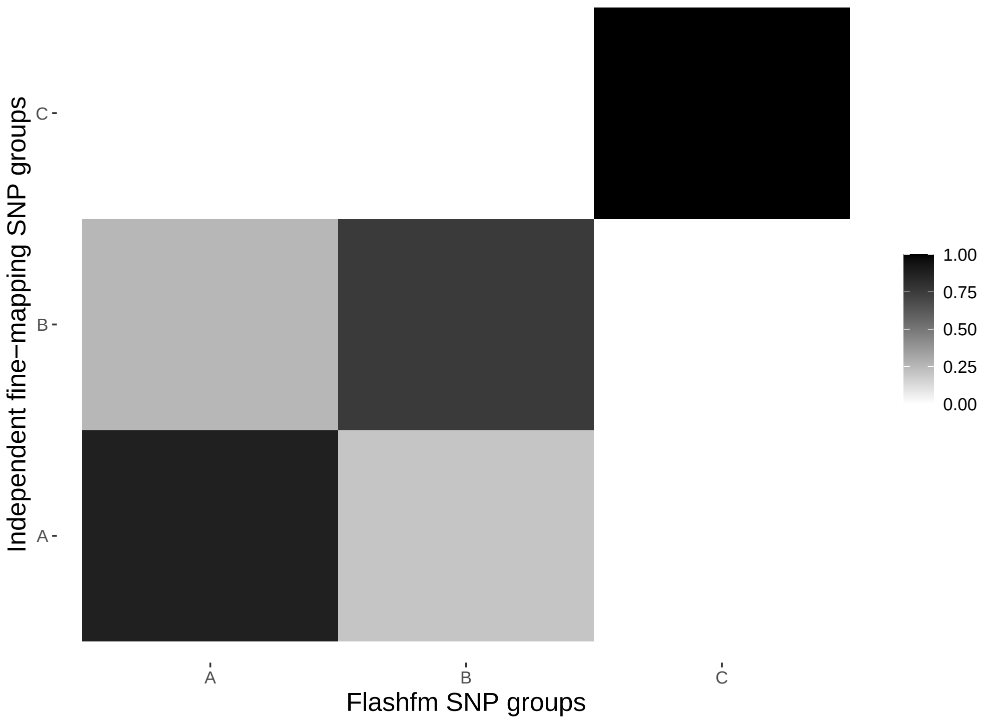

Supplementary Figure 44: Average LD between SNP groups constructed under independent fine-mapping (left) and Flashfm (right) for region 11:5211496-5252794.

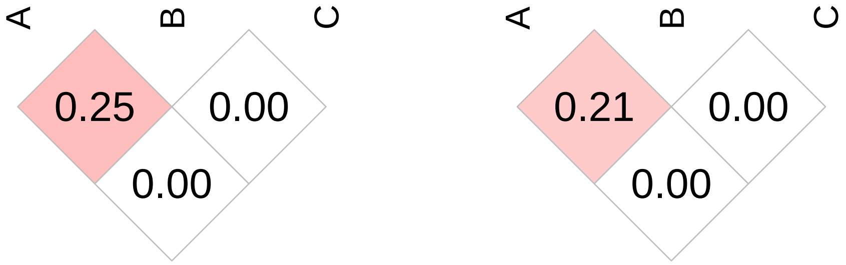

| | Stepwise | | Independent | | Flashfm | |
|---|---|---|---|---|---|---|
| Traits | SNP/Model | P-value | Model | PP | Model | PP |
| rbc_dstr_width | 11:5336134/A | 1.02917e-14 | A | 0.619 | A | 0.684 |
| bilirubin | 11:5334665/A | 3.0441e-07 | A | 0.713 | A | 0.733 |

Supplementary Table 49: Fine-mapping results for 11:5304648-5363496 by independent fine-mapping and Flashfm. Stepwise results are also given.

| | SNP Group |
|---|---|
| | A |
| Independent | 8 |
| | 11:5363496 |
| Flashfm | 3 |
| | 11:5340595 |

Supplementary Table 50: SNP group sizes and representative variant for groups constructed by independent fine-mapping and by Flashfm for region 11:5304648-5363496.

Supplementary Figure 45: Venn diagram of SNP group constructed under indepenedent fine-mapping compared to that under flashfm for region 11:5304648-5363496. This shows that the flashfm group is a subset of, or the same as, the group form independent fine-mapping.

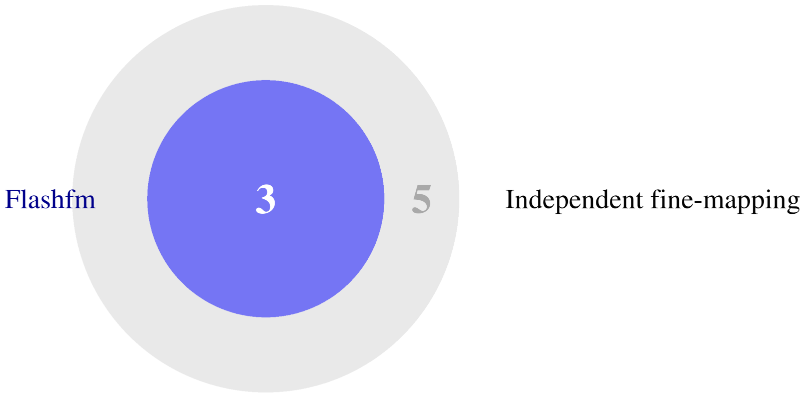

| | Stepwise | | Independent | | Flashfm | |
|---|---|---|---|---|---|---|
| Traits | SNP/Model | P-value | Model | PP | Model | PP |
| rbc_dstr_width | 11:5118578/A | 2.08951e-13 | A | 0.701 | A | 0.885 |
| mchc | 11:5190723/A | 2.34699e-07 | A | 0.593 | A | 0.724 |

Supplementary Table 51: Fine-mapping results for 11:5053165-5190723 by independent fine-mapping and Flashfm. Stepwise results are also given.

| | SNP Group |
|---|---|
| | A |
| Independent | 2 |
| | 11:5118578 |
| Flashfm | 2 |
| | 11:5190723 |

Supplementary Table 52: SNP group sizes and representative variant for groups constructed by independent fine-mapping and by Flashfm for region 11:5053165-5190723.

Supplementary Figure 46: Venn diagram of SNP group constructed under indepenedent fine-mapping compared to that under flashfm for region 11:5053165-5190723. This shows that the flashfm group is a subset of, or the same as, the group form independent fine-mapping.

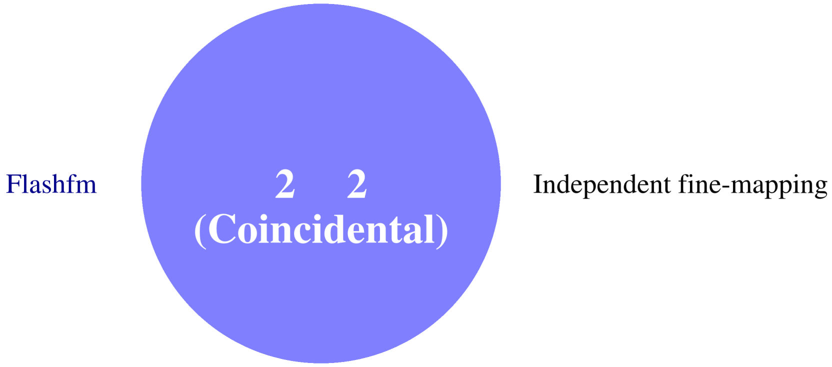

| | Stepwise | | Independent | | Flashfm | |
|---|---|---|---|---|---|---|
| Traits | SNP/Model | P-value | Model | PP | Model | PP |
| cholesterol | 11:84360985/B | 2.20999e-07 | B | 0.823 | B | 0.818 |
| lowdlipo | 11:84360985/B | 2.52085e-07 | B | 0.76 | B | 0.594 |
| | – | – | – | – | B+D | 0.078 |
| | – | – | – | – | B+C | 0.061 |
| | – | – | – | – | A | 0.059 |

Supplementary Table 53: Fine-mapping results for 11:84190922-84413365 by independent fine-mapping and Flashfm. Stepwise results are also given.

| | SNP Group | | | |
|---|---|---|---|---|
| | A | B | C | D |
| Independent | 1 | 12 | 4 | 3 |
| | 11:84356305 | 11:84342305 | 11:84262771 | 11:84349327 |
| Flashfm | 1 | 12 | 4 | 3 |
| | 11:84356305 | 11:84366910 | 11:84262771 | 11:84349327 |

Supplementary Table 54: SNP group sizes and representative variant for groups constructed by independent fine-mapping and by Flashfm for region 11:84190922-84413365.

Supplementary Figure 47: Average LD between SNP groups constructed under independent fine-mapping with those from flashfm for region 11:84190922-84413365.

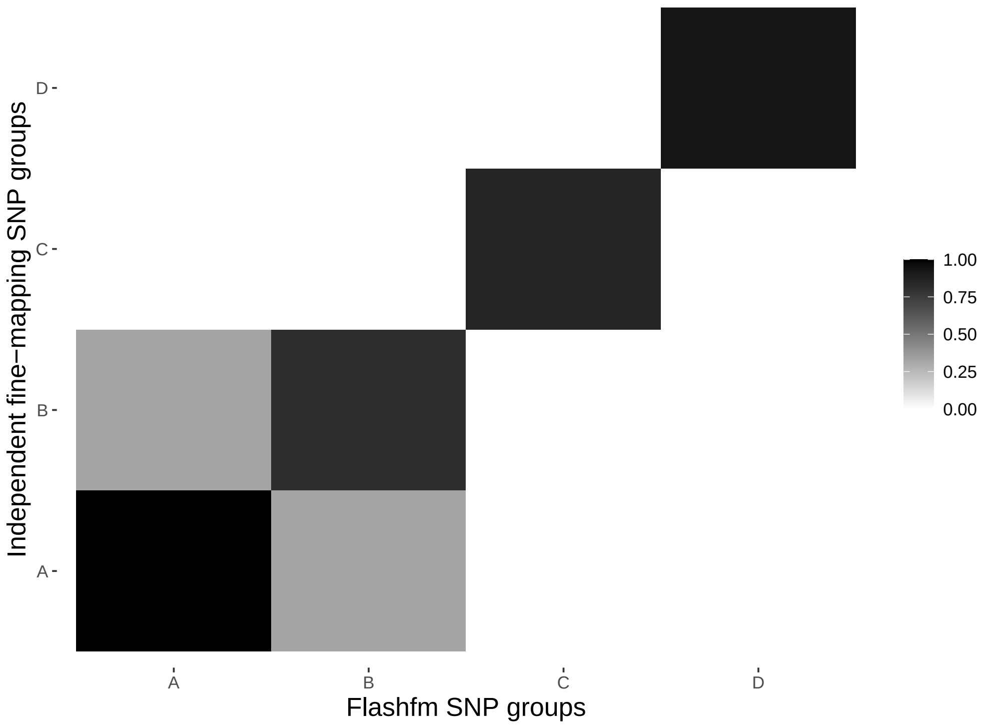

Supplementary Figure 48: Average LD between SNP groups constructed under independent fine-mapping (left) and Flashfm (right) for region 11:84190922-84413365.

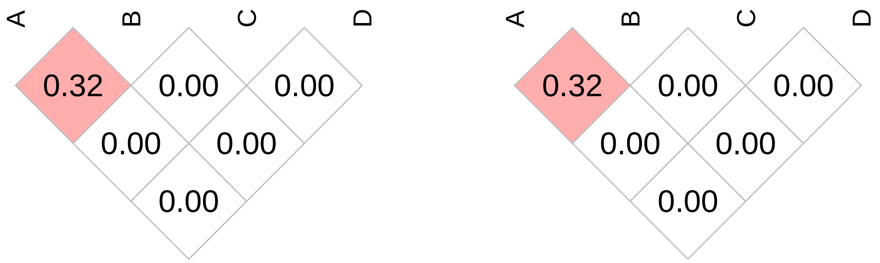

| | Stepwise | | Independent | | Flashfm | |
|---|---|---|---|---|---|---|
| Traits | SNP/Model | P-value | Model | PP | Model | PP |
| hgb | 13:23057646/A | 4.47604e-07 | A | 0.662 | A | 0.744 |
| hct | 13:23057646/A | 6.80465e-07 | A | 0.679 | A | 0.760 |

Supplementary Table 55: Fine-mapping results for 13:23033928-23144347 by independent fine-mapping and Flashfm. Stepwise results are also given.

| | SNP Group |
|---|---|
| | A |
| Independent | 4 |
| | 13:23065854 |
| Flashfm | 4 |
| | 13:23065854 |

Supplementary Table 56: SNP group sizes and representative variant for groups constructed by independent fine-mapping and by Flashfm for region 13:23033928-23144347.

Supplementary Figure 49: Venn diagram of SNP group constructed under indepenedent fine-mapping compared to that under flashfm for region 13:23033928-23144347. This shows that the flashfm group is a subset of, or the same as, the group form independent fine-mapping.

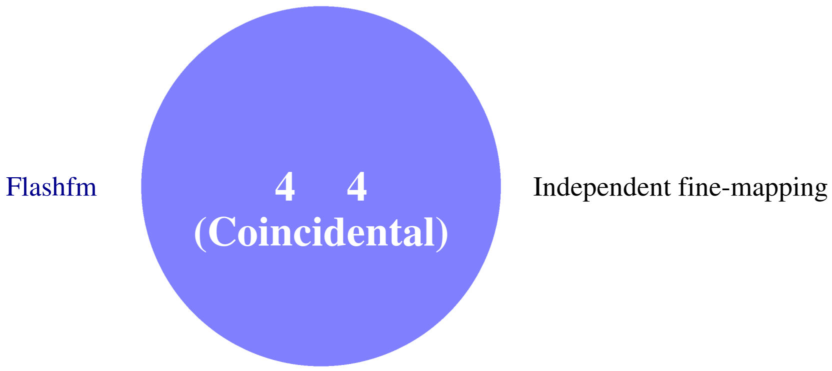

| Traits | Stepwise SNP/Model | Stepwise P-value | Independent Model | Independent PP | Flashfm Model | Flashfm PP |
|---|---|---|---|---|---|---|
| weight | 14:21748646/A | 7.00214e-08 | A | 0.722 | A | 0.763 |
| | – | – | A+F | 0.080 | A+F | 0.057 |
| waist_circum | 14:21748646/A | 1.49727e-07 | A | 0.756 | A | 0.786 |
| bmi | 14:21748646/A | 2.63504e-07 | A | 0.764 | A | 0.803 |
| | – | – | A+D | 0.068 | – | – |

Supplementary Table 57: Fine-mapping results for 14:21673702-21797029 by independent fine-mapping and Flashfm. Stepwise results are also given.

| | SNP Group A | SNP Group D | SNP Group F |
|---|---|---|---|
| Independent | 32 | 12 | 4 |
| | 14:21686690 | 14:21743242 | 14:21692999 |
| Flashfm | 8 | 12 | 4 |
| | 14:21744483 | 14:21743242 | 14:21692999 |

Supplementary Table 58: SNP group sizes and representative variant for groups constructed by independent fine-mapping and by Flashfm for region 14:21673702-21797029.

Supplementary Figure 50: Average LD between SNP groups constructed under independent fine-mapping with those from flashfm for region 14:21673702-21797029.

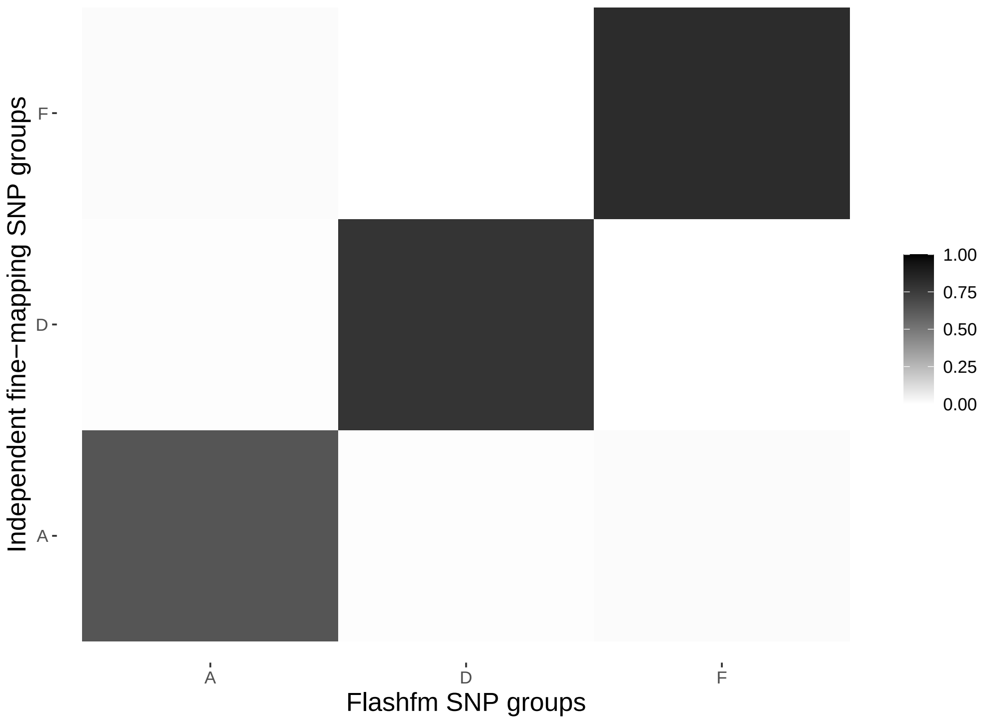

Supplementary Figure 51: Average LD between SNP groups constructed under independent fine-mapping (left) and Flashfm (right) for region 14:21673702-21797029.

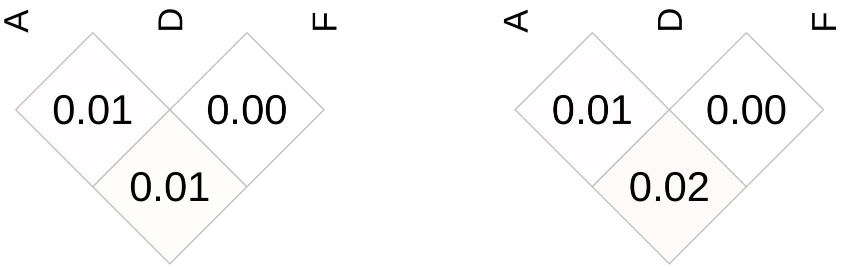

| | Stepwise | | Independent | | Flashfm | |
|---|---|---|---|---|---|---|
| Traits | SNP/Model | P-value | Model | PP | Model | PP |
| highdlipo | 15:58723675/G | 5.53742e-14 | G | 0.427 | G | 0.523 |
| | – | – | G+J | 0.091 | G+J | 0.09 |
| | – | – | C+G | 0.076 | C+G | 0.064 |
| | – | – | G+K | 0.051 | – | – |
| triglycerides | 15:58723426/G | 9.90278e-08 | G | 0.560 | G | 0.658 |

Supplementary Table 59: Fine-mapping results for 15:58718136-58742605 by independent fine-mapping and Flashfm. Stepwise results are also given.

| | SNP Group | | | |
|---|---|---|---|---|
| | C | G | J | K |
| Independent | 6 | 5 | 2 | 9 |
| | 15:58731665 | 15:58724842 | 15:58736623 | 15:58737382 |
| Flashfm | 6 | 4 | 2 | 7 |
| | 15:58731665 | 15:58723675 | 15:58736623 | 15:58735027 |

Supplementary Table 60: SNP group sizes and representative variant for groups constructed by independent fine-mapping and by Flashfm for region 15:58718136-58742605.

Supplementary Figure 52: Average LD between SNP groups constructed under independent fine-mapping with those from flashfm for region 15:58718136-58742605.

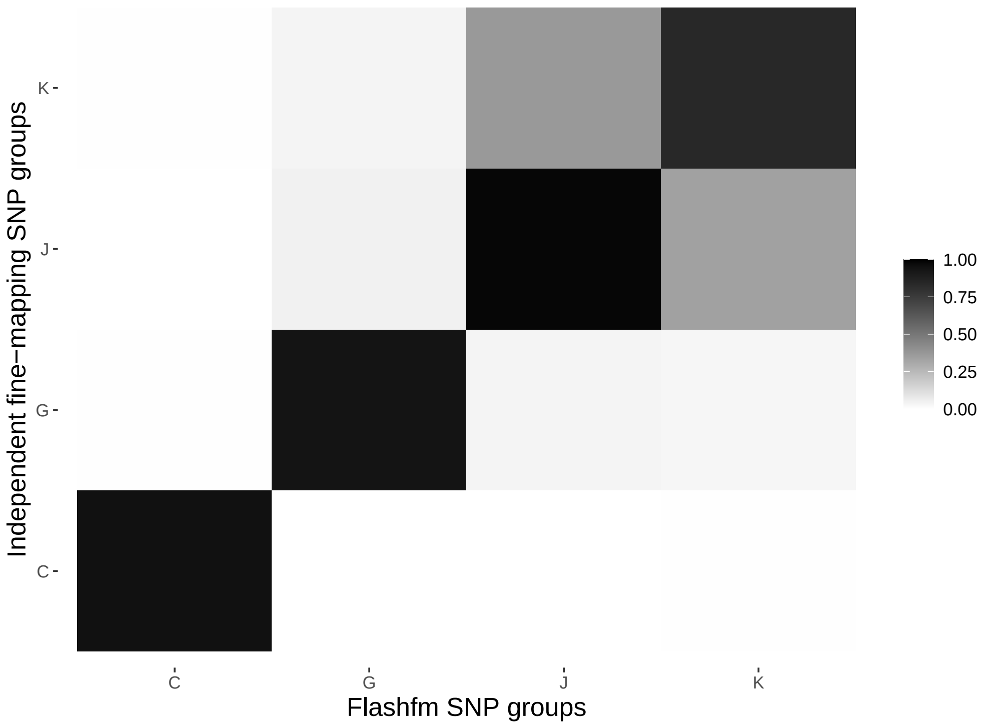

Supplementary Figure 53: Average LD between SNP groups constructed under independent fine-mapping (left) and Flashfm (right) for region 15:58718136-58742605.

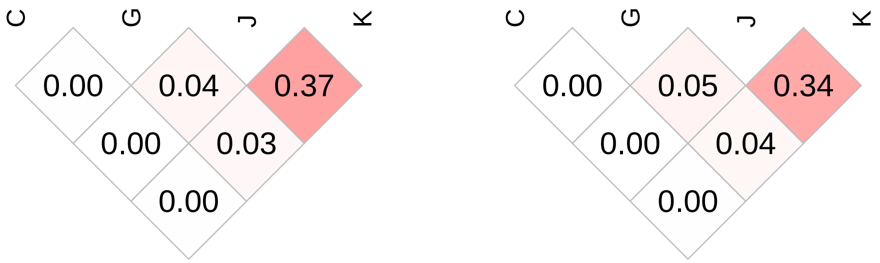

| | Stepwise | | Independent | | Flashfm | |
|---|---|---|---|---|---|---|
| Traits | SNP/Model | P-value | Model | PP | Model | PP |
| cholesterol | 15:55705675/A | 1.71224e-07 | A | 0.365 | A | 0.545 |
| | – | – | A+G | 0.364 | A+G | 0.266 |
| lowdlipo | 15:55705675/A | 4.20627e-07 | A | 0.581 | A | 0.738 |
| | – | – | A+G | 0.227 | A+G | 0.140 |

Supplementary Table 61: Fine-mapping results for 15:55547292-55822875 by independent fine-mapping and Flashfm. Stepwise results are also given.

| | SNP Group | |
|---|---|---|
| | A | G |
| Independent | 8 | 36 |
| | 15:55745354 | 15:55734203 |
| Flashfm | 8 | 36 |
| | 15:55745354 | 15:55734203 |

Supplementary Table 62: SNP group sizes and representative variant for groups constructed by independent fine-mapping and by Flashfm for region 15:55547292-55822875.

Supplementary Figure 54: Average LD between SNP groups constructed under independent fine-mapping with those from flashfm for region 15:55547292-55822875.

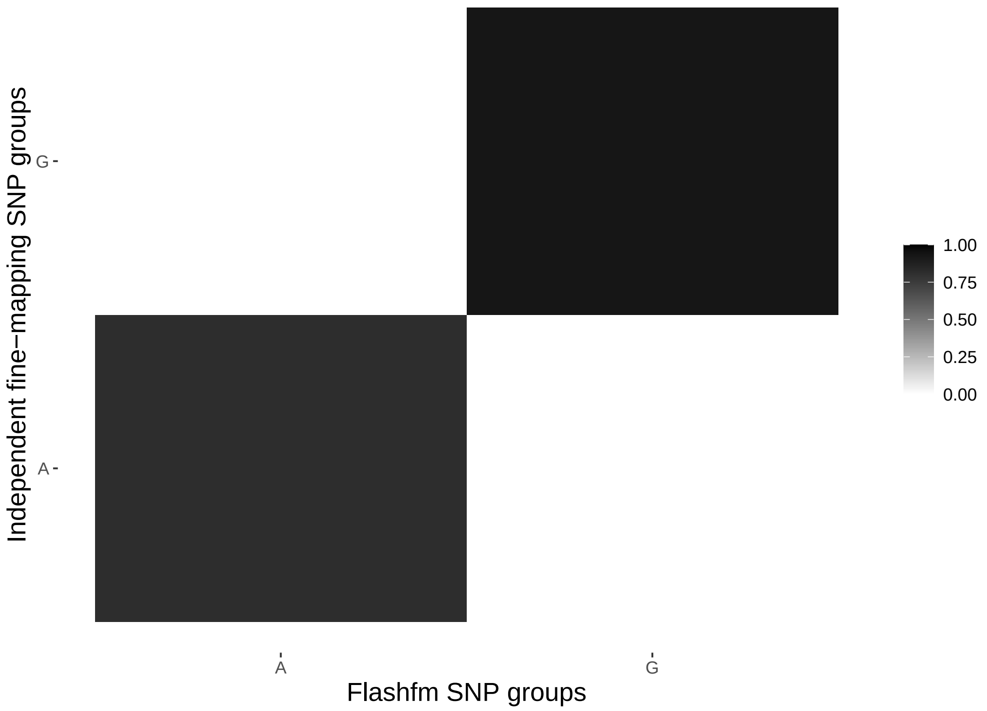

Supplementary Figure 55: Average LD between SNP groups constructed under independent fine-mapping (left) and Flashfm (right) for region 15:55547292-55822875.

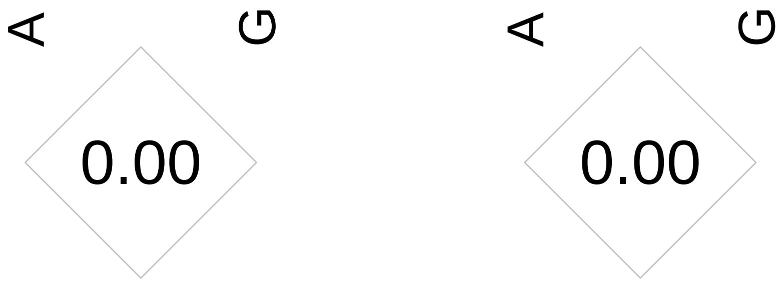

| | Stepwise | | Independent | | Flashfm | |
|---|---|---|---|---|---|---|
| Traits | SNP/Model | P-value | Model | PP | Model | PP |
| rbc | 16:90070/A | 1.10783e-10 | A | 0.215 | A | 0.331 |
| | – | – | A+T | 0.170 | A+T | 0.148 |
| | – | – | A+B | 0.111 | A+B | 0.096 |
| | – | – | A+B+T | 0.108 | A+B+T | 0.058 |
| | – | – | A+M | 0.059 | A+M | 0.055 |
| mchc | 16:88553/A | 1.38189e-10 | A+C | 0.413 | A+C | 0.494 |
| | – | – | A | 0.291 | A | 0.302 |
| hba1c2 | 16:90599/A | 1.72663e-08 | A+H+N | 0.427 | A+H+N | 0.393 |
| | 16:88607/H | 5.72399e-09 | A+H | 0.181 | A+H | 0.254 |
| | – | – | A+H+K | 0.074 | A+H+J | 0.067 |
| | – | – | A+H+M | 0.070 | A+H+K | 0.066 |
| | – | – | A+H+K+N | 0.064 | A+H+M | 0.066 |
| mch | 16:90070/A | 2.74381e-24 | A+C+D+G+J+U | 0.414 | A+C+D+G+J+U | 0.327 |
| | 16:102581/J | 2.01668e-09 | A+C+G+J+M | 0.159 | A+C+G+J | 0.187 |
| | 16:130429/G | 3.15454e-07 | A+C+G+J | 0.110 | A+C+G+J+M | 0.173 |
| | 16:117331/C | 1.01688e-07 | A+C+D+G+J | 0.089 | A+C+D+G+J | 0.098 |
| | – | – | A+C+G+J+U | 0.064 | A+C+G+J+U | 0.078 |

Supplementary Table 63: Fine-mapping results for 16:60647-137716 by independent fine-mapping and Flashfm. Stepwise results are also given.

| | SNP Group | | | | | | |
|---|---|---|---|---|---|---|---|
| | A | B | C | D | G | H | J |
| Independent | 4 | 1 | 2 | 3 | 1 | 5 | 8 |
| | 16:90599 | 16:104329 | 16:117050 | 16:134745 | 16:130429 | 16:88607 | 16:94853 |
| Flashfm | 4 | 1 | 2 | 3 | 1 | 5 | 8 |
| | 16:90599 | 16:104329 | 16:117331 | 16:134745 | 16:130429 | 16:88607 | 16:94853 |
| | K | M | N | T | U | | |
| Independent | 1 | 7 | 6 | 9 | 1 | | |
| | 16:130406 | 16:137634 | 16:128054 | 16:88245 | 16:103592 | | |
| Flashfm | 1 | 7 | 6 | 9 | 1 | | |
| | 16:130406 | 16:137634 | 16:128054 | 16:88245 | 16:103592 | | |

Supplementary Table 64: SNP group sizes and representative variant for groups constructed by independent fine-mapping and by Flashfm for region 16:60647-137716.

Supplementary Figure 56: Average LD between SNP groups constructed under independent fine-mapping with those from flashfm for region 16:60647-137716.

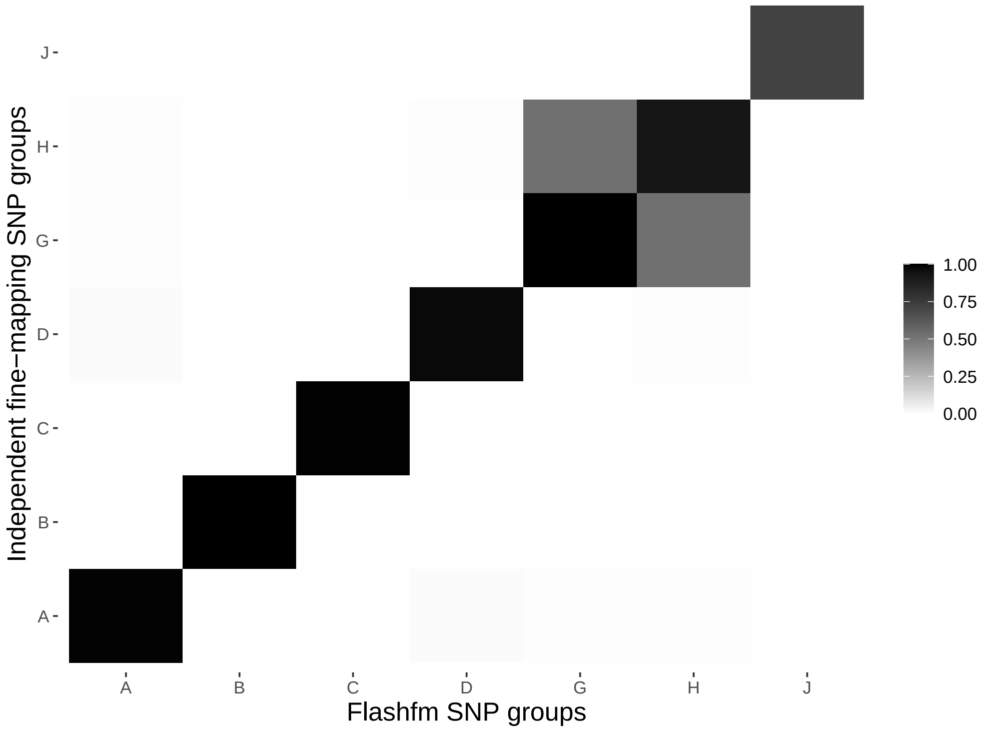

Supplementary Figure 57: Average LD between SNP groups constructed under independent fine-mapping (left) and Flashfm (right) for region 16:60647-137716.

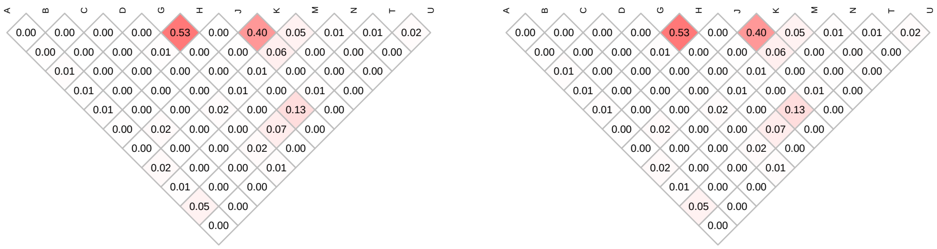

| | Stepwise | | Independent | | Flashfm | |
|---|---|---|---|---|---|---|
| Traits | SNP/Model | P-value | Model | PP | Model | PP |
| bilirubin | 16:603562/B | 1.14991e-07 | B | 0.295 | B | 0.531 |
| | – | – | B+L | 0.254 | B+L | 0.271 |
| | – | – | C | 0.142 | – | – |
| | – | – | C+L | 0.072 | – | – |
| rbc | 16:603562/B | 1.54871e-07 | B+D | 0.511 | B+D | 0.591 |
| | – | – | B+D+F | 0.086 | B | 0.094 |
| | – | – | A+D | 0.056 | B+D+F | 0.075 |
| | – | – | B | 0.053 | – | – |
| mcv | 16:603562/B | 7.55658e-14 | B+D+F | 0.519 | B+D+F | 0.599 |
| | 16:542642/F | 1.70737e-12 | B+D+F+M | 0.092 | B+D+F+M | 0.077 |
| | 16:527280/D | 7.88461e-12 | – | – | – | – |

Supplementary Table 65: Fine-mapping results for 16:498355-605776 by independent fine-mapping and Flashfm. Stepwise results are also given.

| | SNP Group | | | | | | |
|---|---|---|---|---|---|---|---|
| | A | B | C | D | F | L | M |
| Independent | 1 | 11 | 1 | 4 | 3 | 30 | 2 |
| | 16:594790 | 16:600756 | 16:594329 | 16:527280 | 16:526278 | 16:553945 | 16:553917 |
| Flashfm | 0 | 1 | 1 | 4 | 3 | 30 | 2 |
| | – | 16:603562 | 16:594329 | 16:527280 | 16:526278 | 16:553945 | 16:553917 |

Supplementary Table 66: SNP group sizes and representative variant for groups constructed by independent fine-mapping and by Flashfm for region 16:498355-605776.

Supplementary Figure 58: Average LD between SNP groups constructed under independent fine-mapping with those from flashfm for region 16:498355-605776.

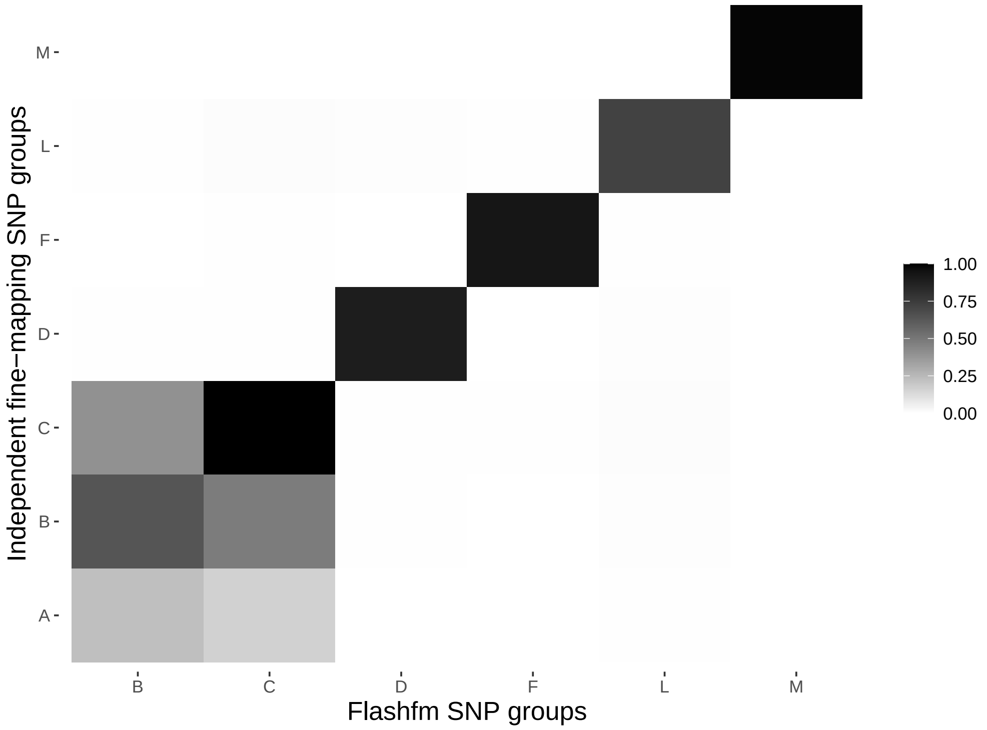

Supplementary Figure 59: Average LD between SNP groups constructed under independent fine-mapping (left) and Flashfm (right) for region 16:498355-605776.

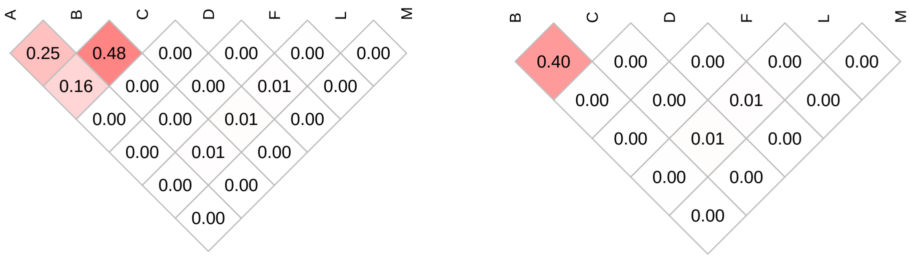

| | Stepwise | | Independent | | Flashfm | |
|---|---|---|---|---|---|---|
| Traits | SNP/Model | P-value | Model | PP | Model | PP |
| mch | 16:1027250/C | 7.09192e-10 | C | 0.584 | C | 0.722 |
| | – | – | A+C | 0.161 | A+C | 0.098 |
| mcv | 16:1027250/C | 2.93089e-09 | C | 0.414 | C | 0.583 |
| | – | – | A+C | 0.295 | A+C | 0.205 |

Supplementary Table 67: Fine-mapping results for 16:939366-1043324 by independent fine-mapping and Flashfm. Stepwise results are also given.

| | SNP Group | |
|---|---|---|
| | A | C |
| Independent | 4 | 5 |
| | 16:949809 | 16:1027250 |
| Flashfm | 4 | 5 |
| | 16:949809 | 16:1027250 |

Supplementary Table 68: SNP group sizes and representative variant for groups constructed by independent fine-mapping and by Flashfm for region 16:939366-1043324.

Supplementary Figure 60: Average LD between SNP groups constructed under independent fine-mapping with those from flashfm for region 16:939366-1043324.

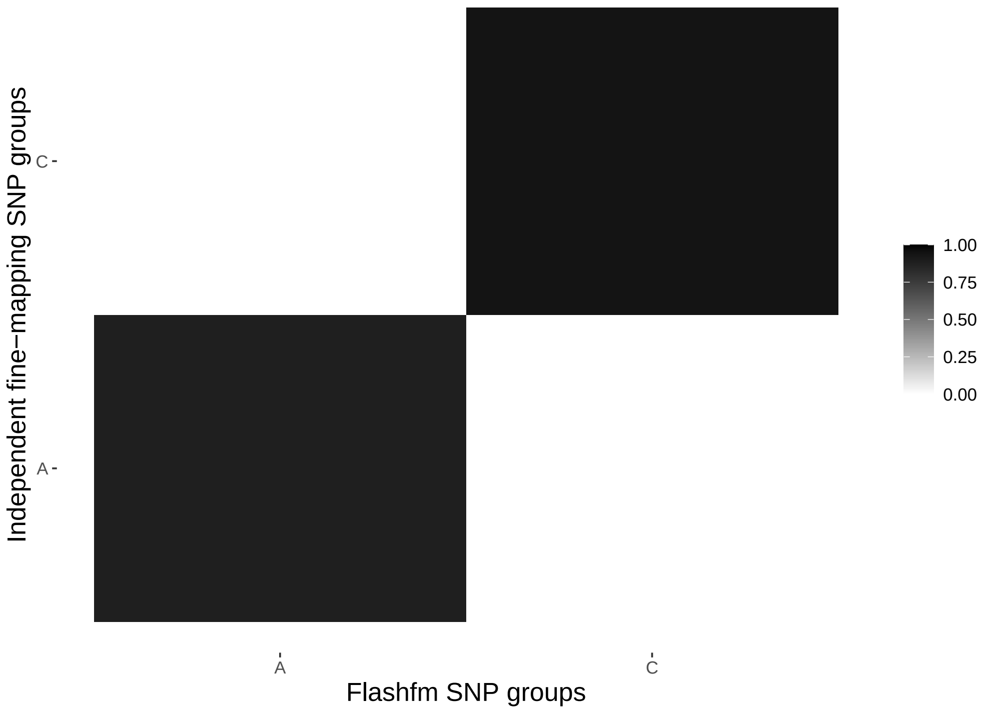

Supplementary Figure 61: Average LD between SNP groups constructed under independent fine-mapping (left) and Flashfm (right) for region 16:939366-1043324.

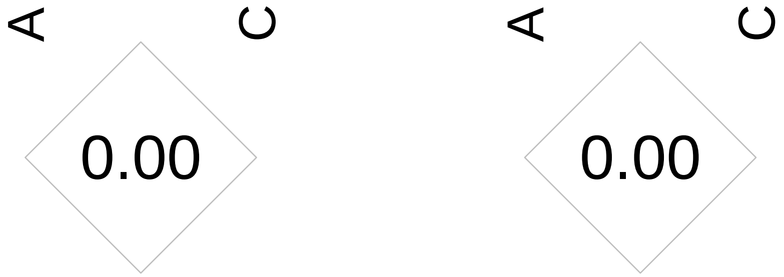

| | Stepwise | | Independent | | Flashfm | |
|---|---|---|---|---|---|---|
| Traits | SNP/Model | P-value | Model | PP | Model | PP |
| highdlipo | 16:56995935/B | 1.1908e-49 | B+D+H+M+T | 0.269 | B+E+J+S+T | 0.148 |
| | 16:56987369/M | 4.38062e-17 | B+E+J+S+T | 0.061 | B+D+H+M+T | 0.116 |
| | 16:57002092/T | 2.07704e-12 | B+D+M+T | 0.055 | B+C+J+S+T | 0.092 |
| | – | – | B+E+J+M+T | 0.053 | B+F+J+S+T | 0.087 |
| | – | – | B+E+M+T | 0.052 | B+E+S+T | 0.080 |
| cholesterol | 16:56991363/S | 6.2228e-08 | S | 0.739 | S | 0.730 |
| | – | – | – | – | S+T | 0.050 |
| triglycerides | 16:56995827/B | 1.21097e-07 | B+K | 0.55 | B+K | 0.604 |
| | – | – | B | 0.063 | B | 0.106 |
| | – | – | K | 0.056 | K+S | 0.059 |
| neut_count | 16:56987960/A | 7.75024e-07 | A | 0.562 | A | 0.322 |
| | – | – | – | – | A+S | 0.231 |
| | – | – | – | – | A+B | 0.153 |

Supplementary Table 69: Fine-mapping results for 16:56971385-57018102 by independent fine-mapping and Flashfm. Stepwise results are also given.

| | SNP Group | | | | | | |
|---|---|---|---|---|---|---|---|
| | A | B | C | D | E | F | H |
| Independent | 4 | 3 | 9 | 1 | 2 | 1 | 10 |
| | 16:56987960 | 16:56990827 | 16:57017002 | 16:57007610 | 16:57007734 | 16:57007451 | 16:56999960 |
| Flashfm | 2 | 2 | 4 | 1 | 2 | 1 | 3 |
| | 16:56987960 | 16:56995827 | 16:57002152 | 16:57007610 | 16:57007734 | 16:57007451 | 16:56998761 |
| | J | K | M | S | T | | |
| Independent | 1 | 11 | 3 | 6 | 2 | | |
| | 16:56996060 | 16:57009985 | 16:56987015 | 16:56993324 | 16:57002092 | | |
| Flashfm | 1 | 11 | 3 | 5 | 2 | | |
| | 16:56996060 | 16:57009985 | 16:56987015 | 16:56996211 | 16:57002092 | | |

Supplementary Table 70: SNP group sizes and representative variant for groups constructed by independent fine-mapping and by Flashfm for region 16:56971385-57018102.

Supplementary Figure 62: Average LD between SNP groups constructed under independent fine-mapping with those from flashfm for region 16:56971385-57018102.



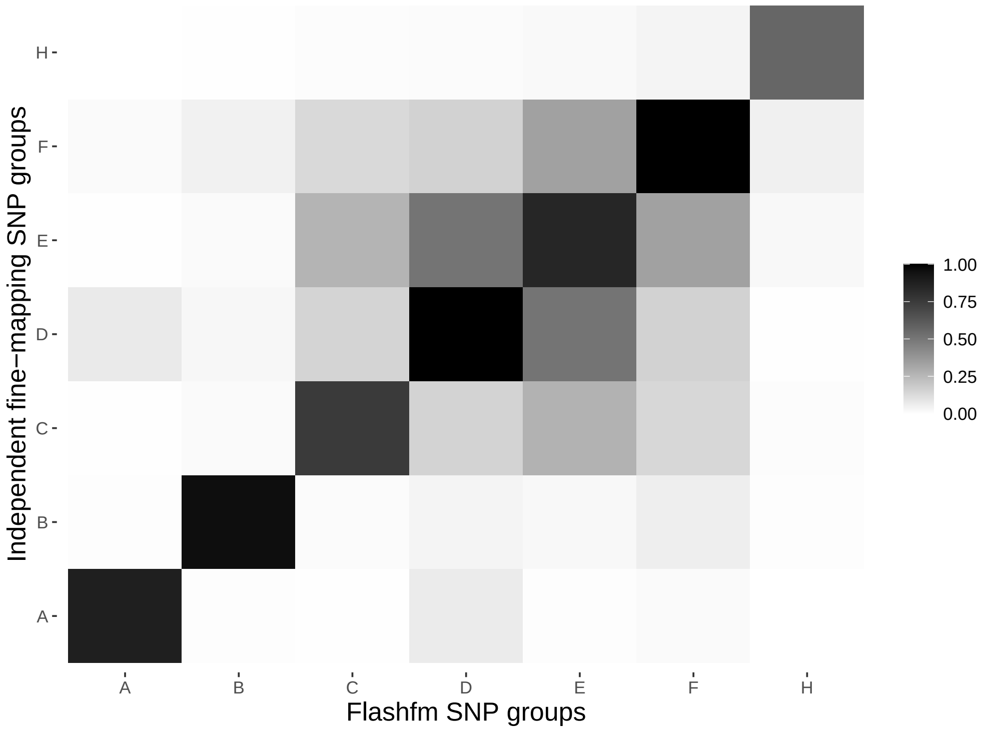

Supplementary Figure 63: Average LD between SNP groups constructed under independent fine-mapping (left) and Flashfm (right) for region 16:56971385-57018102.

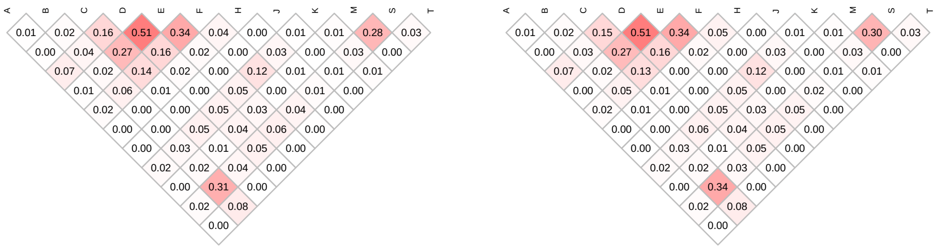

| | Stepwise | | Independent | | Flashfm | |
|---|---|---|---|---|---|---|
| Traits | SNP/Model | P-value | Model | PP | Model | PP |
| mcv | 16:167165/D | 6.25413e-19 | A2+C2+D+E+I+X | 0.202 | A2+C2+D+E+I+X | 0.429 |
| | 16:190254/E | 1.5333e-13 | A+A2+C2+D+E+I+R+X | 0.127 | A2+C2+D+E+I+R+X | 0.136 |
| | 16:172646/C2 | 5.61875e-08 | A+A2+C2+D+E+I | 0.123 | A2+C2+D+E+I+P+X | 0.09 |
| | – | – | A+A2+C2+D+E+I+X | 0.112 | A2+C2+D+E+I+Y | 0.055 |
| | – | – | A+A2+C2+D+E+I+R | 0.099 | – | – |
| | – | – | A2+C2+D+E+I+R+X | 0.076 | – | – |
| | – | – | A+A2+C2+D+E+I+Y | 0.050 | – | – |
| mchc | 16:167165/D | 5.79002e-11 | D+I | 0.362 | D+I | 0.362 |
| | – | – | D+H | 0.346 | D+H | 0.349 |
| | – | – | D | 0.087 | D | 0.148 |
| hba1c2 | 16:150764/U | 5.87248e-08 | D+F+K | 0.375 | D+F+K | 0.416 |
| | – | – | D+F+K+Q | 0.094 | D+F+K+Q | 0.057 |
| | – | – | D+F+K+W | 0.066 | D+F+K+W | 0.053 |
| rbc | 16:163626/D | 4.1364e-08 | D+E | 0.257 | D | 0.426 |
| | – | – | D | 0.192 | D+E | 0.236 |
| | – | – | D+E+Y | 0.108 | D+F | 0.09 |
| | – | – | D+F | 0.080 | D+Y | 0.054 |
| | – | – | D+Y | 0.063 | – | – |

Supplementary Table 71: Fine-mapping results for 16:132064-198333 by independent fine-mapping and Flashfm. Stepwise results are also given.

| | SNP Group | | | | | | |
|---|---|---|---|---|---|---|---|
| | A | C | D | E | F | H | I |
| Independent | 2 | 1 | 10 | 7 | 2 | 13 | 12 |
| | 16:140490 | 16:137591 | 16:167165 | 16:193088 | 16:167408 | 16:195176 | 16:149393 |
| Flashfm | 2 | 0 | 3 | 5 | 2 | 13 | 12 |
| | 16:140490 | – | 16:167165 | 16:193088 | 16:167408 | 16:195176 | 16:149393 |
| | K | P | Q | R | W | X | Y |
| Independent | 1 | 2 | 3 | 1 | 2 | 1 | 1 |
| | 16:196263 | 16:141060 | 16:153657 | 16:192741 | 16:153048 | 16:187520 | 16:173899 |
| Flashfm | 1 | 2 | 3 | 1 | 2 | 1 | 1 |
| | 16:196263 | 16:141060 | 16:153657 | 16:192741 | 16:153048 | 16:187520 | 16:173899 |
| | A2 | C2 | | | | | |
| Independent | 1 | 1 | | | | | |
| | 16:194759 | 16:172646 | | | | | |
| Flashfm | 1 | 1 | | | | | |
| | 16:194759 | 16:172646 | | | | | |



Supplementary Table 72: SNP group sizes and representative variant for groups constructed by independent fine-mapping and by Flashfm for region 16:132064-198333.

Supplementary Figure 64: Average LD between SNP groups constructed under independent fine-mapping with those from flashfm for region 16:132064-198333.

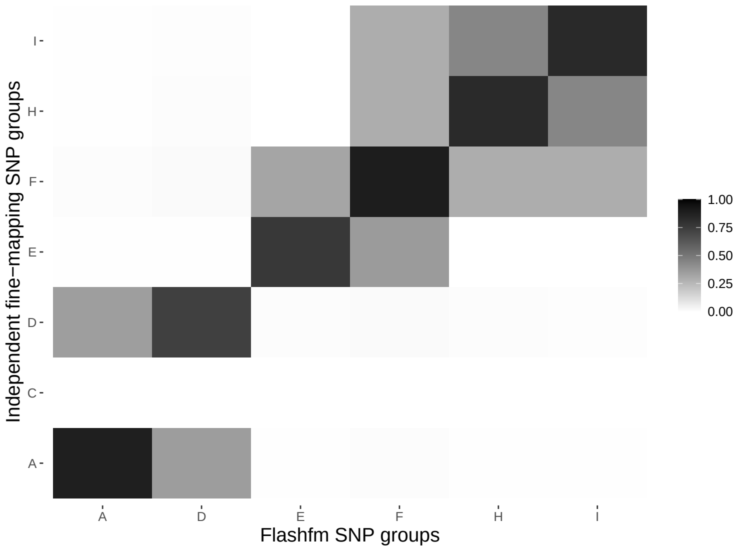

Supplementary Figure 65: Average LD between SNP groups constructed under independent fine-mapping (left) and Flashfm (right) for region 16:132064-198333.

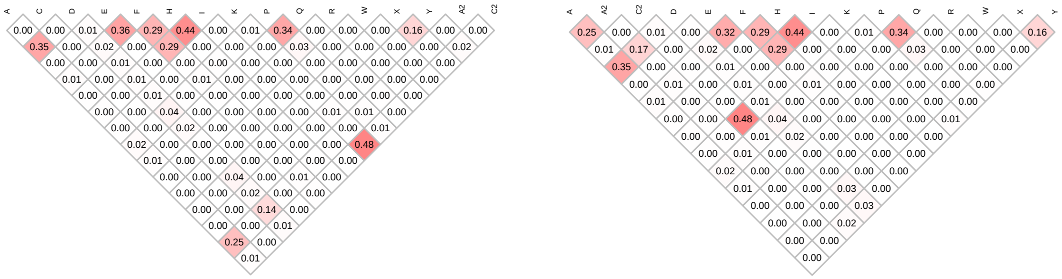

| | Stepwise | | Independent | | Flashfm | |
|---|---|---|---|---|---|---|
| Traits | SNP/Model | P-value | Model | PP | Model | PP |
| highdlipo | 16:57068393/D | 5.21357e-21 | A+D+E+P | 0.383 | A+D+E+P | 0.565 |
| | 16:57077683/P | 4.50615e-10 | D+E+I+P | 0.153 | A+D+G+P | 0.092 |
| | 16:57047299/A | 1.41616e-07 | D+E+P | 0.098 | A+E+J+P | 0.064 |
| | – | – | A+D+G+P | 0.063 | A+E+P | 0.060 |
| triglycerides | 16:57078233/M | 4.05655e-07 | A+M | 0.260 | A+M | 0.433 |
| | – | – | M | 0.142 | A+M+N | 0.137 |
| | – | – | A+M+N | 0.115 | A+M+P | 0.130 |
| | – | – | A+M+P | 0.089 | A+M+O | 0.054 |

Supplementary Table 73: Fine-mapping results for 16:57042326-57084037 by independent fine-mapping and Flashfm. Stepwise results are also given.

| | SNP Group | | | | | | |
|---|---|---|---|---|---|---|---|
| | A | D | E | G | I | J | M |
| Independent | 2 | 6 | 3 | 8 | 1 | 3 | 3 |
| | 16:57047299 | 16:57042326 | 16:57050200 | 16:57058057 | 16:57070005 | 16:57046931 | 16:57078228 |
| Flashfm | 2 | 8 | 3 | 8 | 1 | 3 | 3 |
| | 16:57047299 | 16:57042326 | 16:57050200 | 16:57058057 | 16:57070005 | 16:57046931 | 16:57078228 |
| | N | O | P | | | | |
| Independent | 2 | 1 | 2 | | | | |
| | 16:57052730 | 16:57051833 | 16:57077683 | | | | |
| Flashfm | 2 | 1 | 2 | | | | |
| | 16:57052730 | 16:57051833 | 16:57077683 | | | | |

Supplementary Table 74: SNP group sizes and representative variant for groups constructed by independent fine-mapping and by Flashfm for region 16:57042326-57084037.

Supplementary Figure 66: Average LD between SNP groups constructed under independent fine-mapping with those from flashfm for region 16:57042326-57084037.

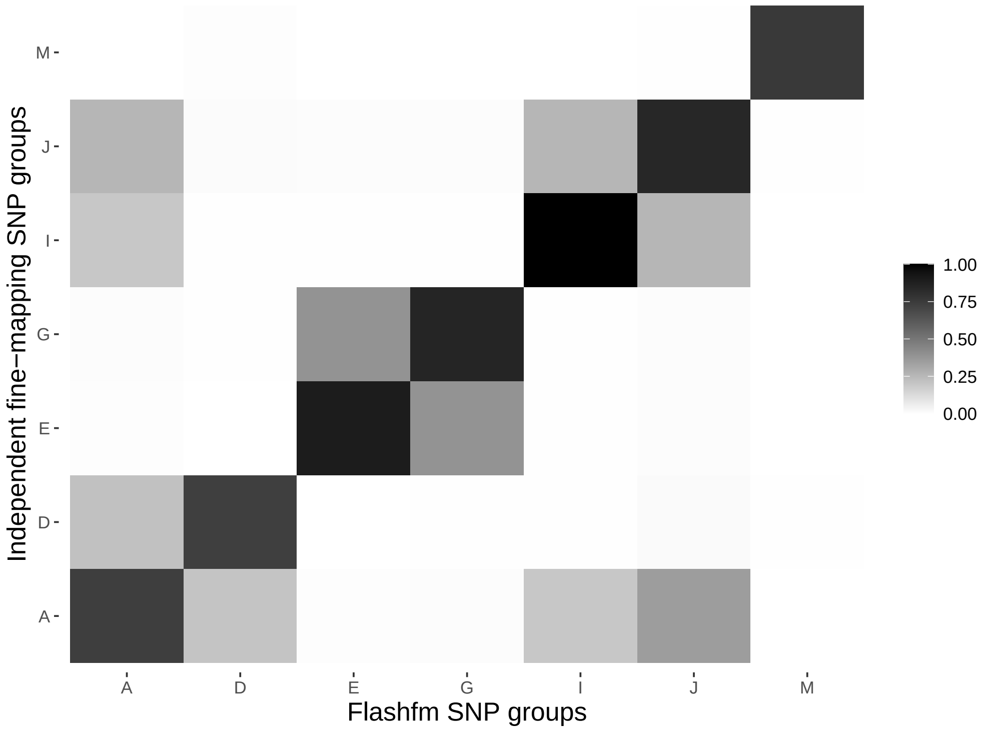

Supplementary Figure 67: Average LD between SNP groups constructed under independent fine-mapping (left) and Flashfm (right) for region 16:57042326-57084037.

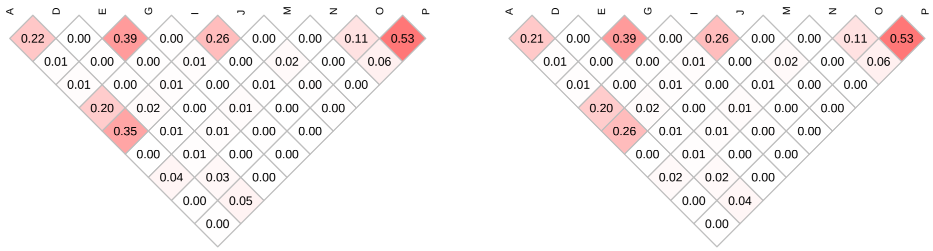

| | Stepwise | | Independent | | Flashfm | |
|---|---|---|---|---|---|---|
| Traits | SNP/Model | P-value | Model | PP | Model | PP |
| bilirubin | 16:321605/G | 9.08431e-12 | G | 0.972 | G | 0.972 |
| rbc | 16:311107/G | 4.2898e-10 | G+J | 0.988 | G+J | 0.988 |
| | 16:366048/J | 2.87208e-09 | G+J | 0.988 | G+J | 0.988 |
| mchc | 16:302161/G | 8.37996e-10 | G+J | 0.605 | G+J | 0.605 |
| | 16:366048/J | 1.85719e-07 | D+G+J | 0.27 | D+G+J | 0.27 |
| | – | – | G+H | 0.054 | G+H | 0.054 |
| hba1c2 | 16:321605/G | 1.16483e-11 | G | 0.745 | G | 0.745 |
| | – | – | C+G | 0.089 | C.1+G | 0.089 |
| mcv | 16:311107/G | 3.75442e-25 | C+G+J | 0.43 | C.2+G+J | 0.407 |
| | 16:366048/J | 2.57128e-21 | C+D+G+J | 0.33 | C.2+D+G+J | 0.314 |
| | 16:368259/C | 1.28459e-10 | C+G+J+K | 0.066 | C.2+G+J+K | 0.063 |
| mch | 16:311107/G | 1.12816e-27 | C+D+G+J+K | 0.486 | C.2+D+G+J+K | 0.482 |
| | 16:366048/J | 4.36966e-23 | A+D+G+J+K | 0.367 | A+D+G+J+K | 0.367 |
| | 16:368259/C | 2.4743e-10 | B+D+G+J+K | 0.088 | B+D+G+J+K | 0.088 |
| | 16:308732/D | 7.46373e-08 | – | – | – | – |

Supplementary Table 75: Fine-mapping results for 16:302161-440165 by independent fine-mapping and Flashfm. Stepwise results are also given.

| | SNP Group | | | | | | |
|---|---|---|---|---|---|---|---|
| | A | B | C | D | G | H | J |
| Independent | 1 | 1 | 5 | 3 | 5 | 1 | 6 |
| | 16:320750 | 16:324837 | 16:347850 | 16:314962 | 16:321605 | 16:320865 | 16:366048 |
| Flashfm | 1 | 1 | 0 | 3 | 5 | 1 | 6 |
| | 16:320750 | 16:324837 | – | 16:314962 | 16:321605 | 16:320865 | 16:366048 |
| | K | C.1 | C.2 | | | | |
| Independent | 2 | 0 | 0 | | | | |
| | 16:349810 | – | – | | | | |
| Flashfm | 2 | 3 | 2 | | | | |
| | 16:349810 | 16:347850 | 16:368259 | | | | |

Supplementary Table 76: SNP group sizes and representative variant for groups constructed by independent fine-mapping and by Flashfm for region 16:302161-440165.

Supplementary Figure 68: Average LD between SNP groups constructed under independent fine-mapping with those from flashfm for region 16:302161-440165.



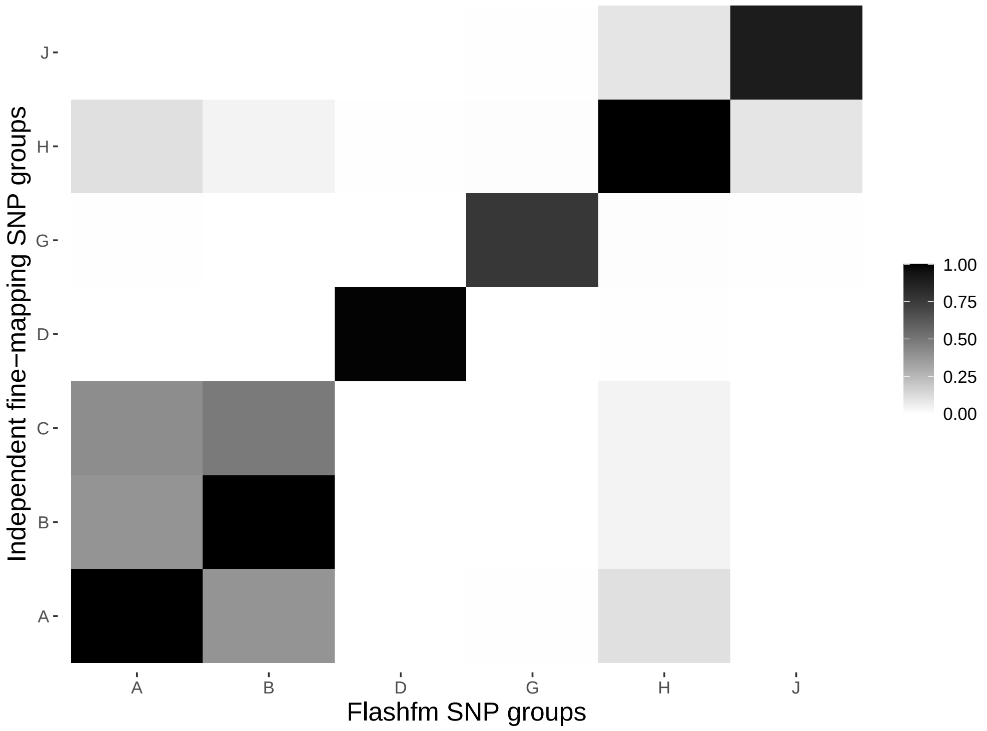

Supplementary Figure 69: Average LD between SNP groups constructed under independent fine-mapping (left) and Flashfm (right) for region 16:302161-440165.

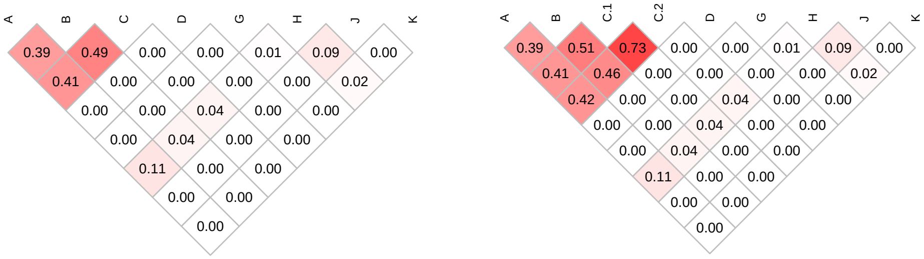

| | Stepwise | | Independent | | Flashfm | |
|---|---|---|---|---|---|---|
| Traits | SNP/Model | P-value | Model | PP | Model | PP |
| mch | 16:603562/V | 4.70522e-15 | K+M+V+W | 0.435 | K+M+V+W | 0.425 |
| | 16:600319/M | 7.56208e-10 | K+M+V | 0.108 | K+M+V | 0.2 |
| | 16:603871/K | 1.67585e-07 | L+M+V+W | 0.105 | L+M+V+W | 0.104 |
| mcv | 16:603562/V | 7.64402e-14 | K+M+V | 0.452 | K+M+V | 0.583 |
| | 16:600319/M | 7.442e-10 | K+M+V+W | 0.102 | K+M+V+W | 0.069 |
| | 16:603871/K | 5.48307e-07 | K+M+N+V | 0.050 | – | – |
| rbc | 16:603562/V | 1.55295e-07 | V | 0.259 | V | 0.517 |
| | – | – | C | 0.097 | E+V | 0.07 |
| | – | – | T | 0.072 | M+V | 0.056 |
| | – | – | E+V | 0.052 | – | – |
| bilirubin | 16:603562/V | 1.15022e-07 | V | 0.252 | V | 0.536 |
| | – | – | U | 0.166 | I+V | 0.071 |
| | – | – | C | 0.052 | L+V | 0.06 |
| | – | – | I+V | 0.051 | – | – |



Supplementary Table 77: Fine-mapping results for 16:589792-618334 by independent fine-mapping and Flashfm. Stepwise results are also given.

| | SNP Group | | | | | | |
|---|---|---|---|---|---|---|---|
| | C | E | I | K | L | M | N |
| Independent | 15 | 1 | 2 | 3 | 3 | 3 | 1 |
| | 16:618334 | 16:600974 | 16:604953 | 16:605794 | 16:601614 | 16:594263 | 16:615720 |
| Flashfm | 0 | 1 | 2 | 3 | 3 | 3 | 1 |
| | – | 16:600974 | 16:604953 | 16:605794 | 16:601614 | 16:594263 | 16:615720 |
| | T | U | V | W | | | |
| Independent | 1 | 1 | 1 | 1 | | | |
| | 16:594790 | 16:594329 | 16:603562 | 16:608254 | | | |
| Flashfm | 1 | 0 | 2 | 1 | | | |
| | 16:594790 | – | 16:603562 | 16:608254 | | | |

Supplementary Table 78: SNP group sizes and representative variant for groups constructed by independent fine-mapping and by Flashfm for region 16:589792-618334.

Supplementary Figure 70: Average LD between SNP groups constructed under independent fine-mapping with those from flashfm for region 16:589792-618334.

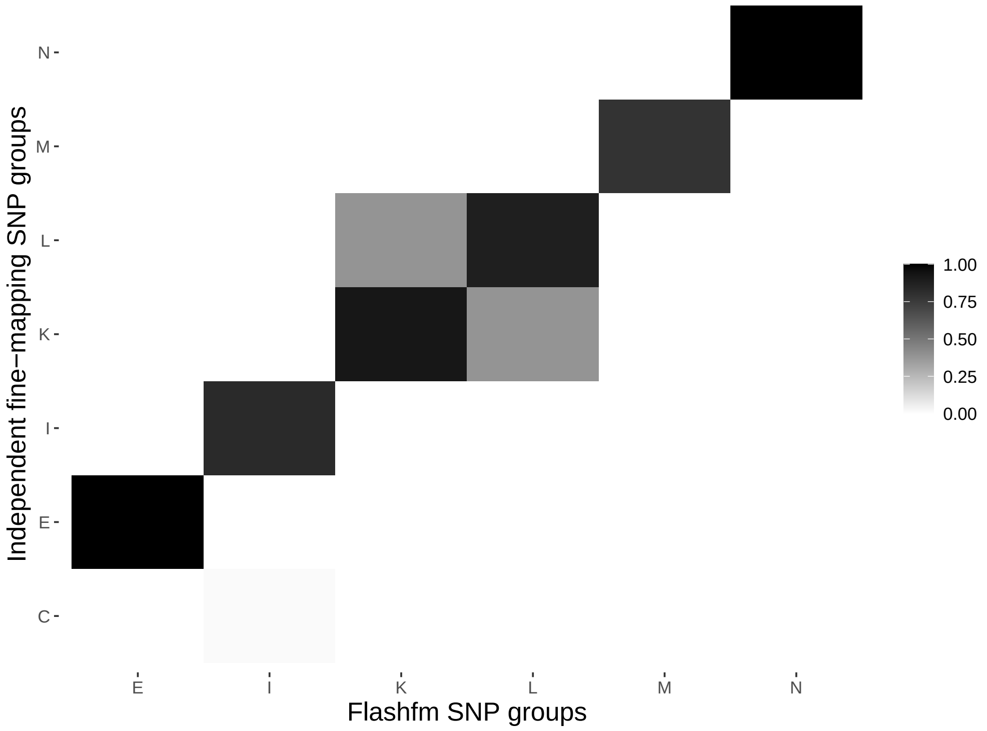

Supplementary Figure 71: Average LD between SNP groups constructed under independent fine-mapping (left) and Flashfm (right) for region 16:589792-618334.

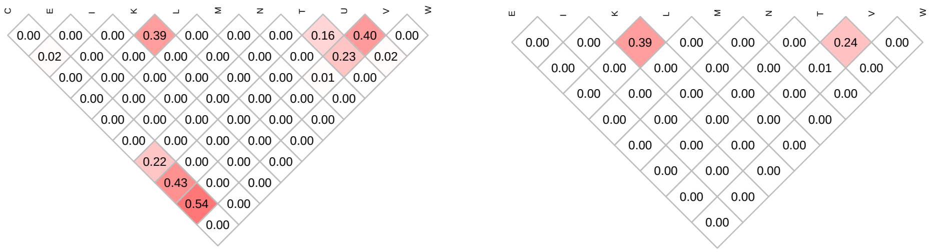

| | Stepwise | | Independent | | Flashfm | |
|---|---|---|---|---|---|---|
| Traits | SNP/Model | P-value | Model | PP | Model | PP |
| bilirubin | 16:603562/C | 1.14991e-07 | C | 0.313 | C | 0.705 |
| | – | – | A | 0.206 | – | – |
| | – | – | B | 0.064 | – | – |
| | – | – | Q | 0.052 | – | – |
| rbc | 16:603562/C | 1.54871e-07 | C | 0.215 | C | 0.553 |
| | – | – | C+I | 0.137 | C+I | 0.229 |
| | – | – | F | 0.072 | – | – |
| | – | – | B | 0.07 | – | – |
| | – | – | E | 0.059 | – | – |
| mcv | 16:603562/C | 7.55658e-14 | C+H+I | 0.619 | C+H+I | 0.688 |
| | 16:600319/I | 7.52621e-10 | C+H+I+L | 0.063 | – | – |
| | 16:603871/H | 5.51157e-07 | – | – | – | – |

Supplementary Table 79: Fine-mapping results for 16:594329-743845 by independent fine-mapping and Flashfm. Stepwise results are also given.

| | SNP Group | | | | | | |
|---|---|---|---|---|---|---|---|
| | A | B | C | E | F | H | I |
| Independent | 1 | 18 | 1 | 1 | 2 | 3 | 20 |
| | 16:594329 | 16:600756 | 16:603562 | 16:594790 | 16:621310 | 16:605794 | 16:600319 |
| Flashfm | 0 | 0 | 2 | 0 | 0 | 3 | 20 |
| | – | – | 16:603562 | – | – | 16:605794 | 16:600319 |
| | L | Q | | | | | |
| Independent | 3 | 3 | | | | | |
| | 16:681587 | 16:729825 | | | | | |
| Flashfm | 3 | 1 | | | | | |
| | 16:681587 | 16:730830 | | | | | |

Supplementary Table 80: SNP group sizes and representative variant for groups constructed by independent fine-mapping and by Flashfm for region 16:594329-743845.

Supplementary Figure 72: Average LD between SNP groups constructed under independent fine-mapping with those from flashfm for region 16:594329-743845.

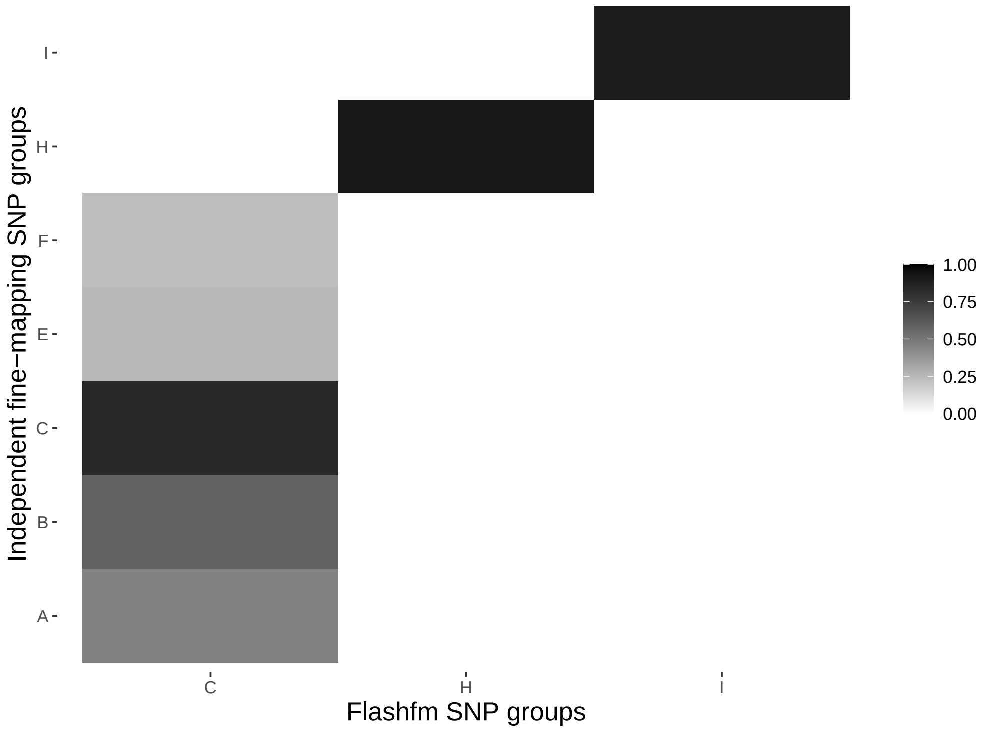

Supplementary Figure 73: Average LD between SNP groups constructed under independent fine-mapping (left) and Flashfm (right) for region 16:594329-743845.

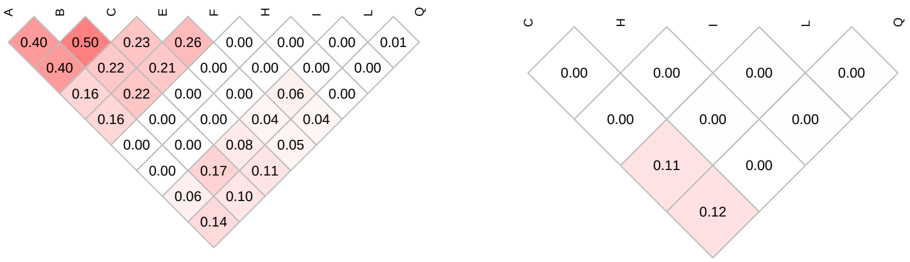

| | Stepwise | | Independent | | Flashfm | |
|---|---|---|---|---|---|---|
| Traits | SNP/Model | P-value | Model | PP | Model | PP |
| mcv | 16:482557/L | 3.25875e-11 | A+B+C+L | 0.423 | A+B+C+L | 0.457 |
| | 16:542642/C | 8.51277e-12 | A+C+L | 0.104 | A+C+L | 0.187 |
| | 16:527280/A | 1.63876e-11 | A+B+C+G+L | 0.095 | A+B+C+G+L | 0.066 |
| | – | – | A+B+C+F+L | 0.064 | – | – |
| mch | 16:482557/L | 3.63114e-11 | A+B+C+L | 0.165 | A+B+C+L | 0.267 |
| | 16:527280/A | 1.46122e-11 | A+B+C+D+L | 0.159 | A+B+C+D+L | 0.164 |
| | 16:542642/C | 1.06721e-10 | A+B+C+L+M | 0.140 | A+B+C+L+M | 0.145 |
| | 16:542041/B | 4.2608e-08 | A+B+C+D+L+M | 0.103 | A+B+C+D+L+M | 0.074 |
| | – | – | A+B+C+D+F+L | 0.055 | – | – |
| bilirubin | 16:482557/L | 1.18192e-07 | L | 0.450 | L | 0.632 |
| | – | – | E+L | 0.186 | E+L | 0.131 |

Supplementary Table 81: Fine-mapping results for 16:441156-557188 by independent fine-mapping and Flashfm. Stepwise results are also given.

| | SNP Group | | | | | | |
|---|---|---|---|---|---|---|---|
| | A | B | C | D | E | F | G |
| Independent | 14 | 1 | 2 | 2 | 6 | 1 | 2 |
| | 16:527280 | 16:542041 | 16:542642 | 16:532430 | 16:548573 | 16:542420 | 16:553919 |
| Flashfm | 14 | 1 | 2 | 2 | 6 | 1 | 2 |
| | 16:527280 | 16:542041 | 16:542642 | 16:532430 | 16:548573 | 16:542420 | 16:553919 |
| | L | M | | | | | |
| Independent | 4 | 1 | | | | | |
| | 16:445085 | 16:530988 | | | | | |
| Flashfm | 1 | 1 | | | | | |
| | 16:482557 | 16:530988 | | | | | |

Supplementary Table 82: SNP group sizes and representative variant for groups constructed by independent fine-mapping and by Flashfm for region 16:441156-557188.

Supplementary Figure 74: Average LD between SNP groups constructed under independent fine-mapping with those from flashfm for region 16:441156-557188.

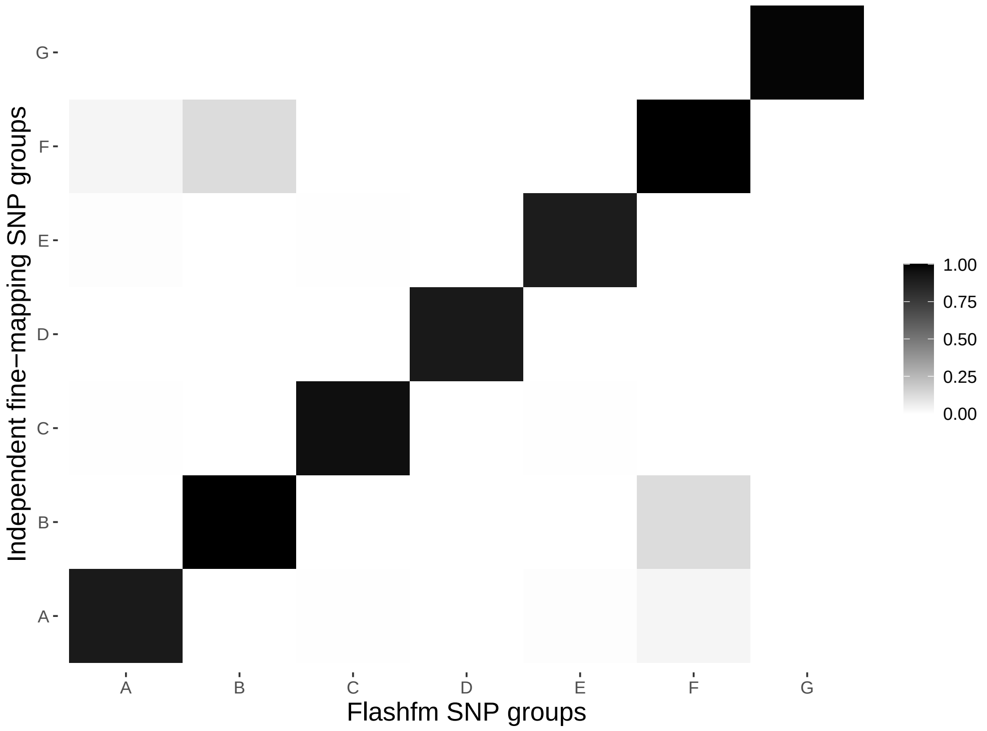

Supplementary Figure 75: Average LD between SNP groups constructed under independent fine-mapping (left) and Flashfm (right) for region 16:441156-557188.

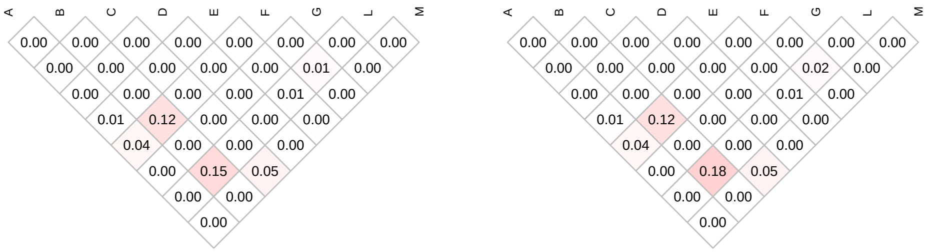

| | Stepwise | | Independent | | Flashfm | |
|---|---|---|---|---|---|---|
| Traits | SNP/Model | P-value | Model | PP | Model | PP |
| mch | 16:911357/D | 3.02107e-09 | D+F | 0.433 | D+F | 0.373 |
| | – | – | D | 0.208 | D | 0.36 |
| mcv | 16:911357/D | 8.6022e-08 | D | 0.323 | D | 0.491 |
| | – | – | D+F | 0.272 | D+F | 0.195 |
| | – | – | D+E | 0.064 | – | – |

Supplementary Table 83: Fine-mapping results for 16:898109-936815 by independent fine-mapping and Flashfm. Stepwise results are also given.

| | SNP Group | | |
|---|---|---|---|
| | D | E | F |
| Independent | 2 | 18 | 1 |
| | 16:911357 | 16:919395 | 16:902670 |
| Flashfm | 2 | 11 | 1 |
| | 16:911357 | 16:904358 | 16:902670 |

Supplementary Table 84: SNP group sizes and representative variant for groups constructed by independent fine-mapping and by Flashfm for region 16:898109-936815.

Supplementary Figure 76: Average LD between SNP groups constructed under independent fine-mapping with those from flashfm for region 16:898109-936815.

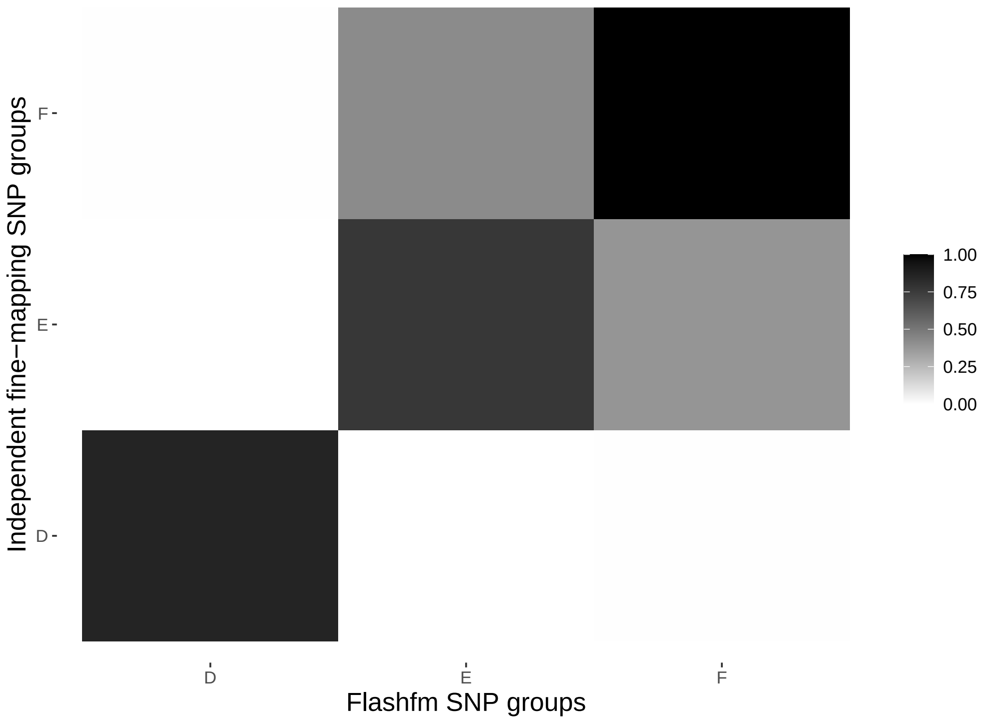

Supplementary Figure 77: Average LD between SNP groups constructed under independent fine-mapping (left) and Flashfm (right) for region 16:898109-936815.

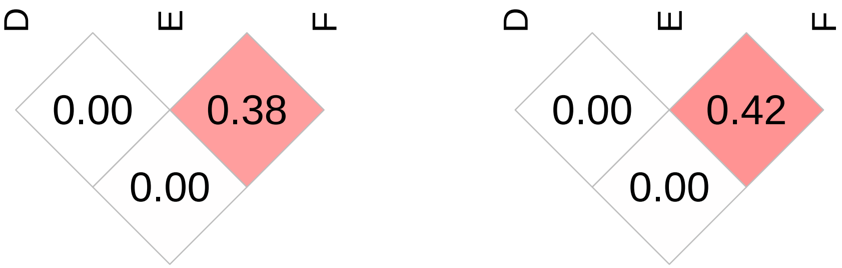

| | Stepwise | | Independent | | Flashfm | |
|---|---|---|---|---|---|---|
| Traits | SNP/Model | P-value | Model | PP | Model | PP |
| hba1c2 | 16:771234/J | 1.41831e-08 | J | 0.547 | J | 0.383 |
| | – | – | B+J | 0.057 | I+J | 0.275 |
| | – | – | – | – | I | 0.085 |
| mcv | 16:775654/L | 1.86525e-07 | A+I | 0.467 | A+I | 0.513 |
| | – | – | A+E+I | 0.082 | A+F+I | 0.051 |
| | – | – | A+F+I | 0.05 | – | – |
| bilirubin | 16:750762/K | 3.7458e-07 | K | 0.779 | K | 0.638 |
| | – | – | – | – | I | 0.092 |
| | – | – | – | – | I+K | 0.068 |

Supplementary Table 85: Fine-mapping results for 16:729957-794205 by independent fine-mapping and Flashfm. Stepwise results are also given.

| | SNP Group | | | | | | |
|---|---|---|---|---|---|---|---|
| | A | B | E | F | I | J | K |
| Independent | 1 | 10 | 14 | 1 | 4 | 3 | 22 |
| | 16:784765 | 16:767430 | 16:772848 | 16:776983 | 16:783004 | 16:766690 | 16:741581 |
| Flashfm | 1 | 13 | 10 | 1 | 4 | 4 | 22 |
| | 16:784765 | 16:766001 | 16:772848 | 16:776983 | 16:783004 | 16:771286 | 16:741581 |

Supplementary Table 86: SNP group sizes and representative variant for groups constructed by independent fine-mapping and by Flashfm for region 16:729957-794205.

Supplementary Figure 78: Average LD between SNP groups constructed under independent fine-mapping with those from flashfm for region 16:729957-794205.

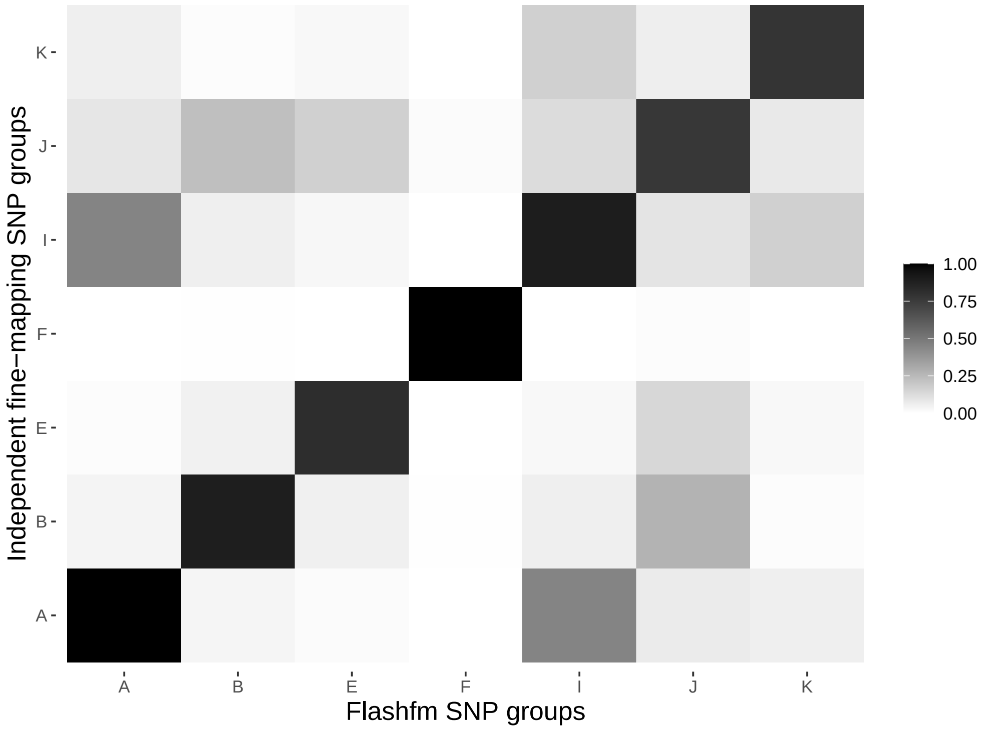

Supplementary Figure 79: Average LD between SNP groups constructed under independent fine-mapping (left) and Flashfm (right) for region 16:729957-794205.

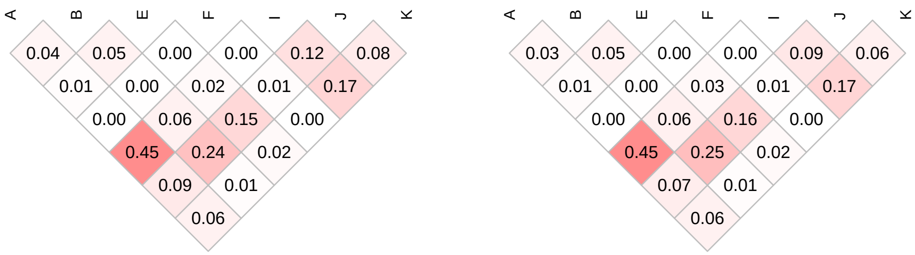

| | Stepwise | | Independent | | Flashfm | |
|---|---|---|---|---|---|---|
| Traits | SNP/Model | P-value | Model | PP | Model | PP |
| mcv | 16:881333/B | 1.10539e-07 | B+E | 0.343 | B | 0.312 |
| | – | – | B | 0.209 | B+E | 0.305 |
| | – | – | B+K | 0.063 | B+K | 0.056 |
| mch | 16:907039/D | 2.32685e-08 | D+J | 0.181 | D | 0.248 |
| | – | – | D | 0.142 | D+J | 0.153 |
| | – | – | B | 0.072 | B | 0.105 |
| | – | – | B+K | 0.068 | B+K | 0.057 |
| | – | – | B+E | 0.053 | – | – |

Supplementary Table 87: Fine-mapping results for 16:863336-907039 by independent fine-mapping and Flashfm. Stepwise results are also given.

| | SNP Group | | | | |
|---|---|---|---|---|---|
| | B | D | E | J | K |
| Independent | 4 | 1 | 2 | 1 | 1 |
| | 16:882657 | 16:907039 | 16:896321 | 16:902670 | 16:867961 |
| Flashfm | 4 | 1 | 2 | 1 | 1 |
| | 16:882657 | 16:907039 | 16:896321 | 16:902670 | 16:867961 |

Supplementary Table 88: SNP group sizes and representative variant for groups constructed by independent fine-mapping and by Flashfm for region 16:863336-907039.

Supplementary Figure 80: Average LD between SNP groups constructed under independent fine-mapping with those from flashfm for region 16:863336-907039.

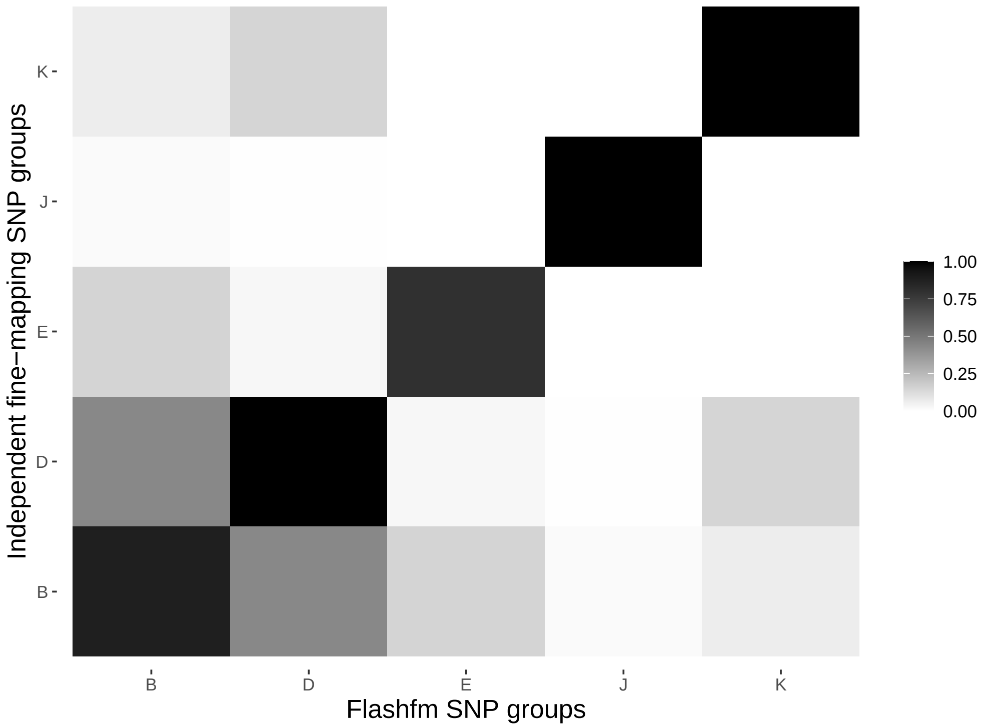

Supplementary Figure 81: Average LD between SNP groups constructed under independent fine-mapping (left) and Flashfm (right) for region 16:863336-907039.

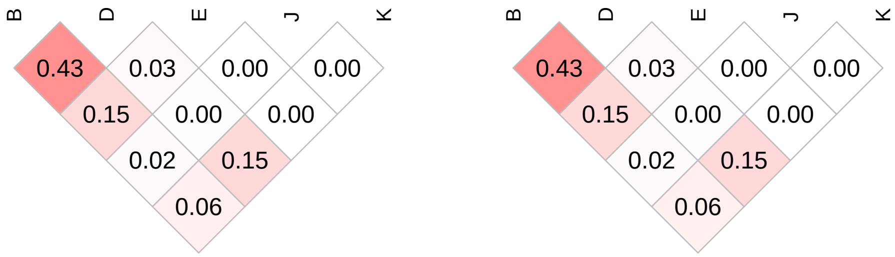

| | Stepwise | | Independent | | Flashfm | |
|---|---|---|---|---|---|---|
| Traits | SNP/Model | P-value | Model | PP | Model | PP |
| cholesterol | 17:75278052/G | 1.64919e-08 | G | 0.575 | G | 0.693 |
| lowdlipo | 17:75278824/K | 1.72825e-07 | K | 0.429 | K | 0.498 |
| | – | – | H | 0.116 | H | 0.135 |
| | – | – | G | 0.068 | G | 0.074 |

Supplementary Table 89: Fine-mapping results for 17:75251326-75306310 by independent fine-mapping and Flashfm. Stepwise results are also given.

| | SNP Group | | |
|---|---|---|---|
| | G | H | K |
| Independent | 3 | 3 | 2 |
| | 17:75286406 | 17:75255973 | 17:75278824 |
| Flashfm | 3 | 3 | 2 |
| | 17:75286406 | 17:75255973 | 17:75278824 |



Supplementary Table 90: SNP group sizes and representative variant for groups constructed by independent fine-mapping and by Flashfm for region 17:75251326-75306310.

Supplementary Figure 82: Average LD between SNP groups constructed under independent fine-mapping with those from flashfm for region 17:75251326-75306310.

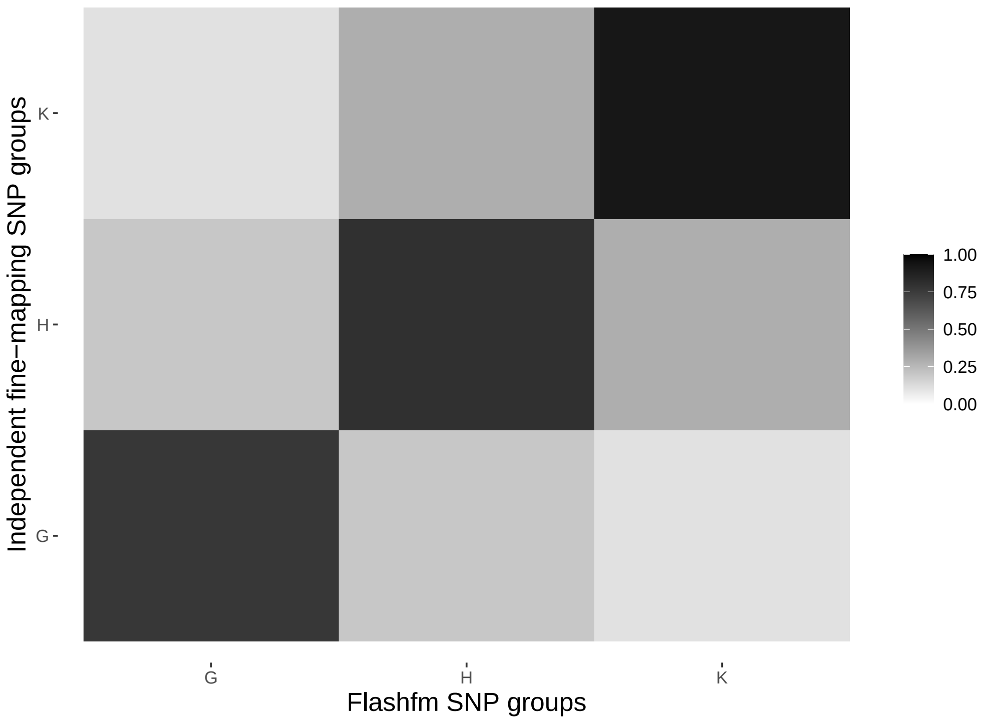

Supplementary Figure 83: Average LD between SNP groups constructed under independent fine-mapping (left) and Flashfm (right) for region 17:75251326-75306310.

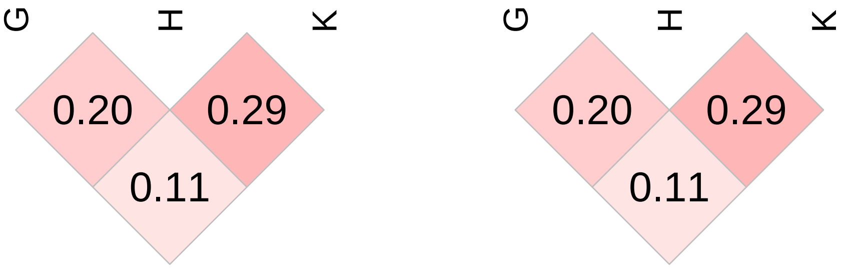

| | Stepwise | | Independent | | Flashfm | |
|---|---|---|---|---|---|---|
| Traits | SNP/Model | P-value | Model | PP | Model | PP |
| hip_circum | 18:8236712/A | 4.45867e-07 | A | 0.546 | A | 0.650 |
| | – | – | C | 0.101 | C | 0.062 |
| | – | – | A+E | 0.059 | A+E | 0.052 |
| waist_circum | 18:8236712/A | 4.74792e-07 | A | 0.690 | A | 0.773 |

Supplementary Table 91: Fine-mapping results for 18:8173997-8257404 by independent fine-mapping and Flashfm. Stepwise results are also given.

| | SNP Group | | |
|---|---|---|---|
| | A | C | E |
| Independent | 5 | 18 | 1 |
| | 18:8257404 | 18:8219357 | 18:8240811 |
| Flashfm | 5 | 11 | 1 |
| | 18:8257404 | 18:8219357 | 18:8240811 |

Supplementary Table 92: SNP group sizes and representative variant for groups constructed by independent fine-mapping and by Flashfm for region 18:8173997-8257404.

Supplementary Figure 84: Average LD between SNP groups constructed under independent fine-mapping with those from flashfm for region 18:8173997-8257404.

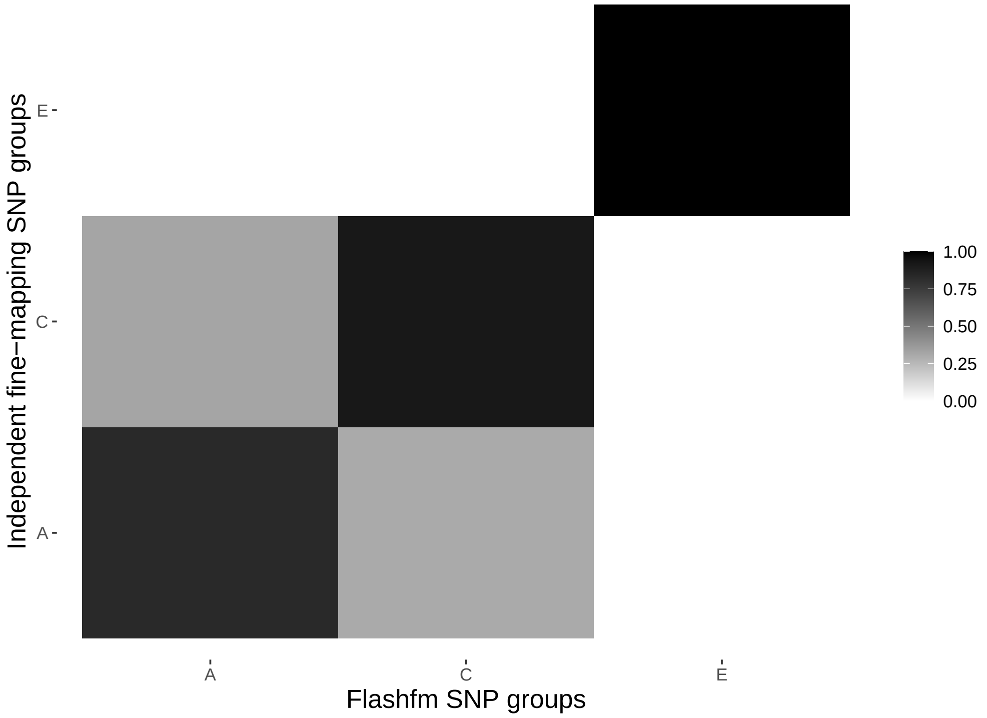

Supplementary Figure 85: Average LD between SNP groups constructed under independent fine-mapping (left) and Flashfm (right) for region 18:8173997-8257404.

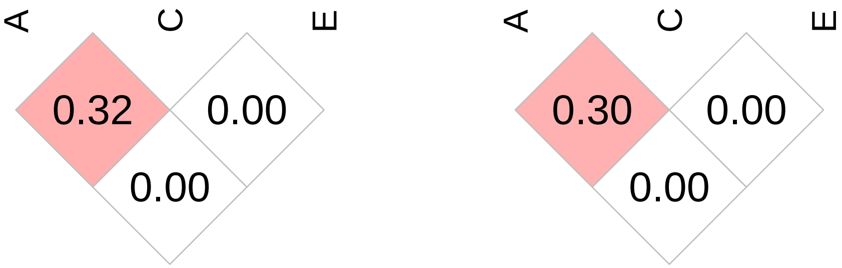

| | Stepwise | | Independent | | Flashfm | |
|---|---|---|---|---|---|---|
| Traits | SNP/Model | P-value | Model | PP | Model | PP |
| lowdlipo | 19:45437250/O | 7.08139e-19 | D+H+N+O | 0.095 | D+H+N+O | 0.126 |
| | 19:45513417/D | 1.82342e-12 | D+G+H+M+O | 0.093 | D+G+H+M+O | 0.079 |
| | – | – | D+H+M+O | 0.072 | D+H+M+O | 0.075 |
| | – | – | D+F+H+M+O | 0.056 | D+G+H+N+O | 0.052 |
| cholesterol | 19:45437250/O | 1.03633e-10 | D+O | 0.393 | D+O | 0.520 |
| | 19:45517497/D | 6.02608e-08 | C+D+O | 0.219 | C+D+O | 0.171 |
| | – | – | D+H+O | 0.153 | D+H+O | 0.119 |

Supplementary Table 93: Fine-mapping results for 19:45437250-45521976 by independent fine-mapping and Flashfm. Stepwise results are also given.

| | SNP Group | | | | | |
|---|---|---|---|---|---|---|
| | C | D | F | G | H | M |
| Independent | 4 | 3 | 1 | 18 | 17 | 1 |
| | 19:45451647 | 19:45513417 | 19:45519263 | 19:45518110 | 19:45480987 | 19:45439163 |
| Flashfm | 4 | 3 | 1 | 18 | 17 | 1 |
| | 19:45451647 | 19:45513417 | 19:45519263 | 19:45518110 | 19:45480987 | 19:45439163 |
| | N | O | | | | |
| Independent | 3 | 3 | | | | |
| | 19:45453353 | 19:45516768 | | | | |
| Flashfm | 3 | 3 | | | | |
| | 19:45453353 | 19:45516768 | | | | |

Supplementary Table 94: SNP group sizes and representative variant for groups constructed by independent fine-mapping and by Flashfm for region 19:45437250-45521976.

Supplementary Figure 86: Average LD between SNP groups constructed under independent fine-mapping with those from flashfm for region 19:45437250-45521976.

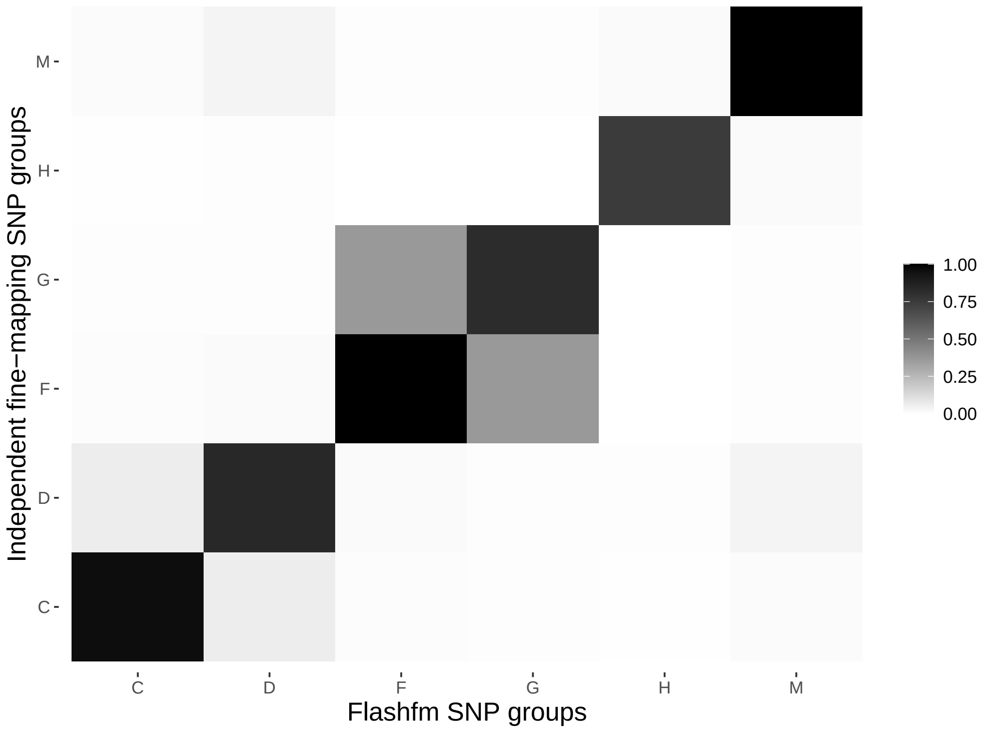

Supplementary Figure 87: Average LD between SNP groups constructed under independent fine-mapping (left) and Flashfm (right) for region 19:45437250-45521976.

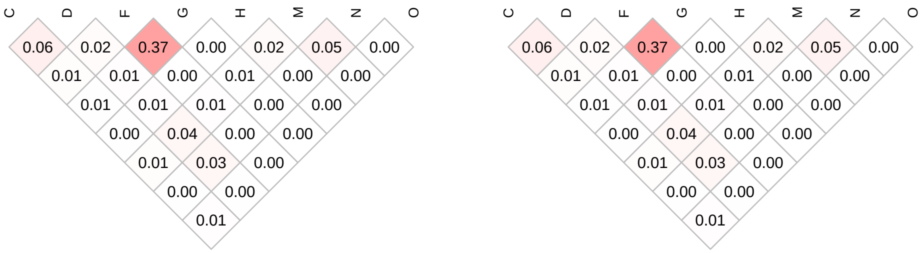

| | Stepwise | | Independent | | Flashfm | |
|---|---|---|---|---|---|---|
| Traits | SNP/Model | P-value | Model | PP | Model | PP |
| highdlipo | 19:21749298/C | 2.3559e-09 | C | 0.695 | C | 0.748 |
| eos_count | 19:21800425/A | 2.32322e-08 | A | 0.880 | A | 0.902 |

Supplementary Table 95: Fine-mapping results for 19:21431997-21822742 by independent fine-mapping and Flashfm. Stepwise results are also given.

| | SNP Group | |
|---|---|---|
| | A | C |
| Independent | 11 | 1 |
| | 19:21800425 | 19:21749298 |
| Flashfm | 11 | 1 |
| | 19:21800425 | 19:21749298 |

Supplementary Table 96: SNP group sizes and representative variant for groups constructed by independent fine-mapping and by Flashfm for region 19:21431997-21822742.

Supplementary Figure 88: Average LD between SNP groups constructed under independent fine-mapping with those from flashfm for region 19:21431997-21822742.

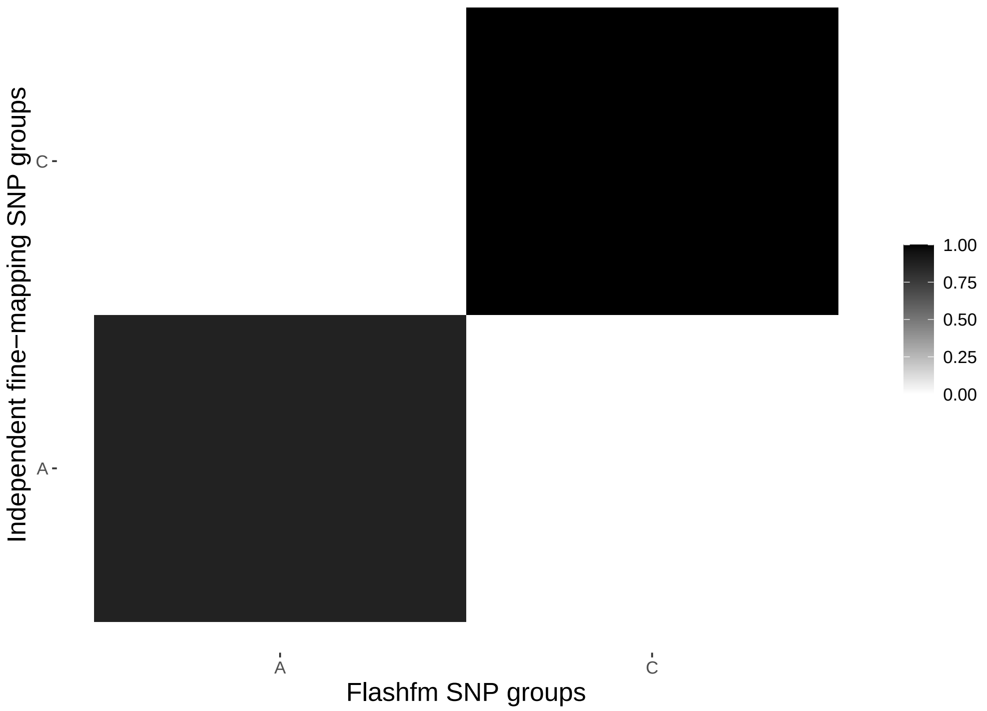

Supplementary Figure 89: Average LD between SNP groups constructed under independent fine-mapping (left) and Flashfm (right) for region 19:21431997-21822742.

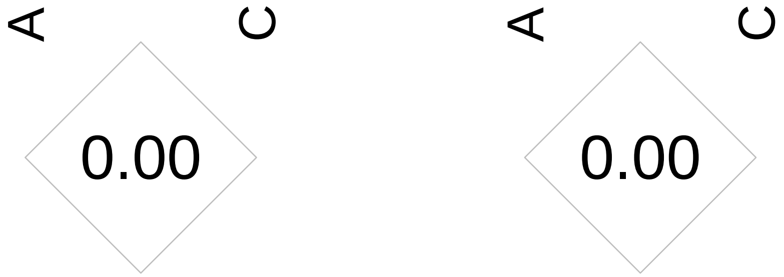

| | Stepwise | | Independent | | Flashfm | |
|---|---|---|---|---|---|---|
| Traits | SNP/Model | P-value | Model | PP | Model | PP |
| lowdlipo | 19:45412079/B | 3.42883e-81 | B+D2+E+L+V | 0.406 | B+D2+E+L+V | 0.391 |
| | 19:45406538/V | 3.04338e-23 | B+C2+D2+E+L+V | 0.279 | B+C2+D2+E+L+V | 0.338 |
| | 19:45410273/E | 7.36491e-16 | – | – | – | – |
| | 19:45411941/L | 6.51514e-13 | – | – | – | – |
| | 19:45427780/D2 | 3.62525e-07 | – | – | – | – |
| cholesterol | 19:45412079/B | 2.3615e-38 | B+E+L+V | 0.265 | B+E+L+V | 0.418 |
| | 19:45406538/V | 3.82465e-11 | B+D2+E+L+V | 0.256 | B+D2+E+L+V | 0.322 |
| | 19:45411941/L | 1.02278e-09 | B+E+L+N+V | 0.068 | B+E+L+N+V | 0.06 |
| | – | – | B+D2+E+L+N+V | 0.06 | – | – |
| triglycerides | 19:45422587/I | 8.78181e-15 | E+E2+I+Y | 0.424 | E+E2+I+X | 0.59 |
| | 19:45430280/X | 2.88626e-07 | E+E2+I+X | 0.214 | E+E2+I+Y | 0.174 |
| | – | – | E2+I+X | 0.067 | E+E2+I+W | 0.056 |
| highdlipo | 19:45413576/A | 1.73275e-09 | A | 0.382 | B | 0.414 |
| | – | – | B | 0.108 | B+I | 0.249 |
| | – | – | A+H | 0.052 | B+K | 0.097 |

Supplementary Table 97: Fine-mapping results for 19:45380937-45441453 by independent fine-mapping and Flashfm. Stepwise results are also given.

| | SNP Group | | | | | | |
|---|---|---|---|---|---|---|---|
| | A | B | C | D | E | H | I |
| Independent | 1 | 2 | 1 | 1 | 5 | 7 | 1 |
| | 19:45413576 | 19:45400747 | 19:45435661 | 19:45392254 | 19:45401587 | 19:45418961 | 19:45422587 |
| Flashfm | 1 | 1 | 0 | 0 | 5 | 0 | 1 |
| | 19:45413576 | 19:45412079 | – | – | 19:45401587 | – | 19:45422587 |
| | K | L | N | V | W | X | Y |
| Independent | 3 | 1 | 2 | 1 | 1 | 1 | 3 |
| | 19:45425171 | 19:45411941 | 19:45418086 | 19:45406538 | 19:45431658 | 19:45430280 | 19:45436216 |
| Flashfm | 4 | 1 | 2 | 1 | 1 | 1 | 3 |
| | 19:45429177 | 19:45411941 | 19:45418086 | 19:45406538 | 19:45431658 | 19:45430280 | 19:45436216 |
| | C2 | D2 | E2 | | | | |
| Independent | 1 | 2 | 4 | | | | |
| | 19:45439163 | 19:45427780 | 19:45422160 | | | | |
| Flashfm | 1 | 2 | 3 | | | | |
| | 19:45439163 | 19:45427780 | 19:45422160 | | | | |



Supplementary Table 98: SNP group sizes and representative variant for groups constructed by independent fine-mapping and by Flashfm for region 19:45380937-45441453.

Supplementary Figure 90: Average LD between SNP groups constructed under independent fine-mapping with those from flashfm for region 19:45380937-45441453.

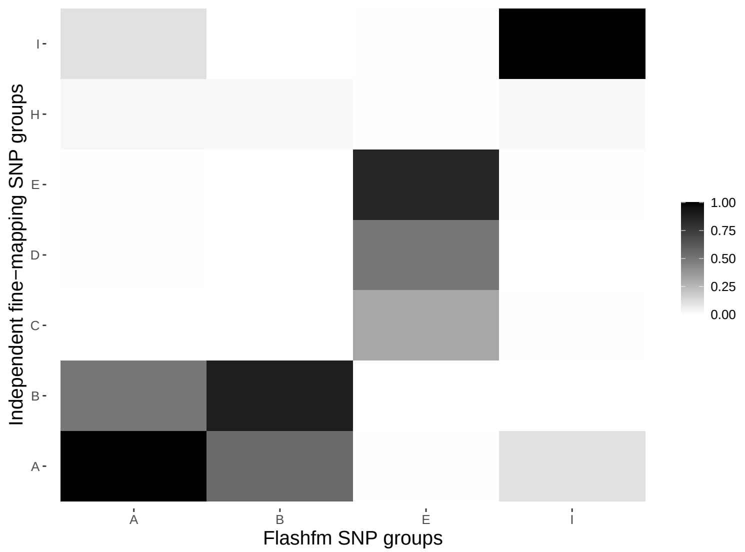

Supplementary Figure 91: Average LD between SNP groups constructed under independent fine-mapping (left) and Flashfm (right) for region 19:45380937-45441453.

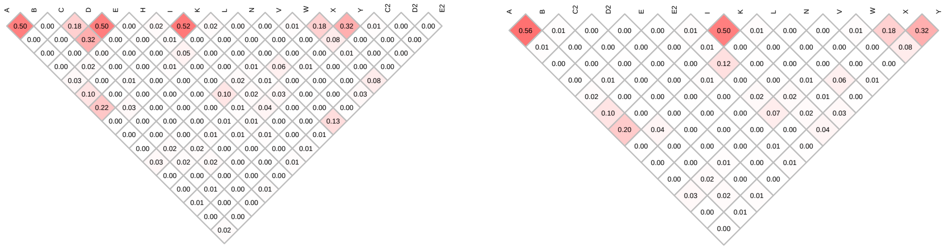

| | Stepwise | | Independent | | Flashfm | |
|---|---|---|---|---|---|---|
| Traits | SNP/Model | P-value | Model | PP | Model | PP |
| lowdlipo | 19:11197261/I | 5.54756e-17 | I+J | 0.136 | I+J | 0.175 |
| | – | – | F+I+J+M | 0.126 | F+I+J | 0.091 |
| | – | – | F+I+J | 0.093 | A.1+B+J | 0.08 |
| | – | – | A+B+J | 0.079 | F+I+J+M | 0.076 |
| | – | – | I+J+M | 0.068 | I+J+M | 0.064 |
| | – | – | G+I+J+M | 0.066 | A.1+J | 0.061 |
| | – | – | A+J+K | 0.058 | A.1+J+K | 0.058 |
| | – | – | – | – | G+I+J+M | 0.05 |
| cholesterol | 19:11197261/I | 2.16355e-11 | I+J | 0.481 | I+J | 0.518 |
| | – | – | A+J | 0.166 | A.1+J | 0.182 |
| | – | – | F+I+J | 0.061 | – | – |

Supplementary Table 99: Fine-mapping results for 19:11163968-11215226 by independent fine-mapping and Flashfm. Stepwise results are also given.

| | SNP Group | | | | | | |
|---|---|---|---|---|---|---|---|
| | A | B | F | G | I | J | K |
| Independent | 11 | 2 | 3 | 1 | 10 | 5 | 2 |
| | 19:11189764 | 19:11194245 | 19:11171793 | 19:11168261 | 19:11197750 | 19:11173045 | 19:11207457 |
| Flashfm | 0 | 2 | 3 | 1 | 17 | 5 | 2 |
| | – | 19:11194245 | 19:11171793 | 19:11168261 | 19:11189764 | 19:11173045 | 19:11207457 |
| | M | A.1 | | | | | |
| Independent | 6 | 0 | | | | | |
| | 19:11189047 | – | | | | | |
| Flashfm | 4 | 3 | | | | | |
| | 19:11189047 | 19:11196886 | | | | | |

Supplementary Table 100: SNP group sizes and representative variant for groups constructed by independent fine-mapping and by Flashfm for region 19:11163968-11215226.

Supplementary Figure 92: Average LD between SNP groups constructed under independent fine-mapping with those from flashfm for region 19:11163968-11215226.



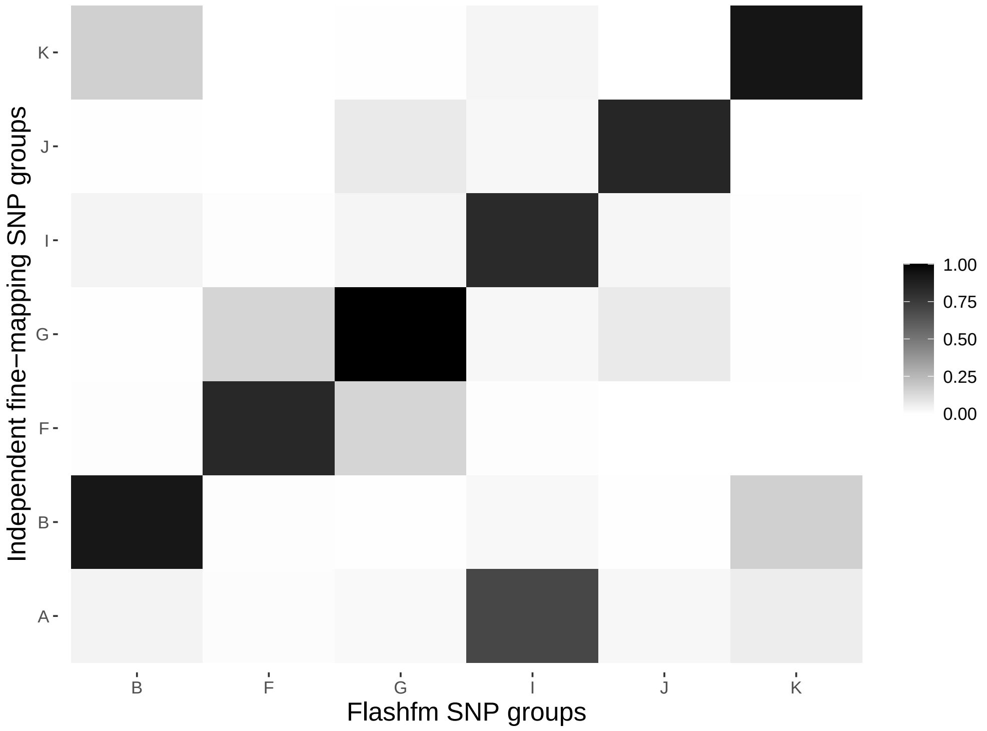

Supplementary Figure 93: Average LD between SNP groups constructed under independent fine-mapping (left) and Flashfm (right) for region 19:11163968-11215226.

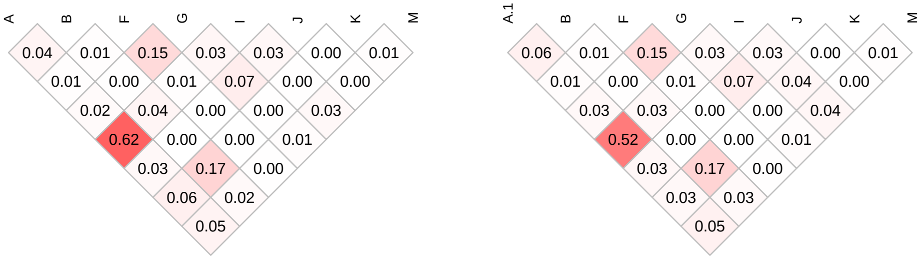

| | Stepwise | | Independent | | Flashfm | |
|---|---|---|---|---|---|---|
| Traits | SNP/Model | P-value | Model | PP | Model | PP |
| lowdlipo | 19:45397229/R | 4.41906e-42 | C+D+E+N+O+R | 0.183 | D+E+N+O+R | 0.195 |
| | 19:45396665/H | 4.78125e-10 | D+E+N+O+R | 0.146 | C+D+E+N+O+R | 0.160 |
| | 19:45390911/O | 7.74729e-07 | D+E+J+N+O+R | 0.110 | D+E+J+N+O+R | 0.113 |
| | – | – | C+D+E+I+N+O+R | 0.076 | C+D+E+I+N+O+R | 0.058 |
| cholesterol | 19:45397229/R | 1.17885e-18 | C+O+R | 0.210 | C+R | 0.235 |
| | – | – | C+R | 0.194 | C+O+R | 0.196 |
| | – | – | C+F+O+R | 0.079 | C+F+R | 0.074 |
| | – | – | C+F+R | 0.077 | C+F+O+R | 0.060 |

Supplementary Table 101: Fine-mapping results for 19:45359667-45397471 by independent fine-mapping and Flashfm. Stepwise results are also given.

| | SNP Group | | | | | | |
|---|---|---|---|---|---|---|---|
| | C | D | E | F | I | J | N |
| Independent | 7 | 2 | 1 | 2 | 1 | 6 | 1 |
| | 19:45374350 | 19:45388581 | 19:45396240 | 19:45375764 | 19:45381420 | 19:45377334 | 19:45392254 |
| Flashfm | 7 | 2 | 1 | 2 | 1 | 6 | 1 |
| | 19:45374350 | 19:45388581 | 19:45396240 | 19:45375764 | 19:45381420 | 19:45377334 | 19:45392254 |
| | O | R | | | | | |
| Independent | 3 | 1 | | | | | |
| | 19:45390911 | 19:45397229 | | | | | |
| Flashfm | 3 | 1 | | | | | |
| | 19:45390911 | 19:45397229 | | | | | |

Supplementary Table 102: SNP group sizes and representative variant for groups constructed by independent fine-mapping and by Flashfm for region 19:45359667-45397471.

Supplementary Figure 94: Average LD between SNP groups constructed under independent fine-mapping with those from flashfm for region 19:45359667-45397471.

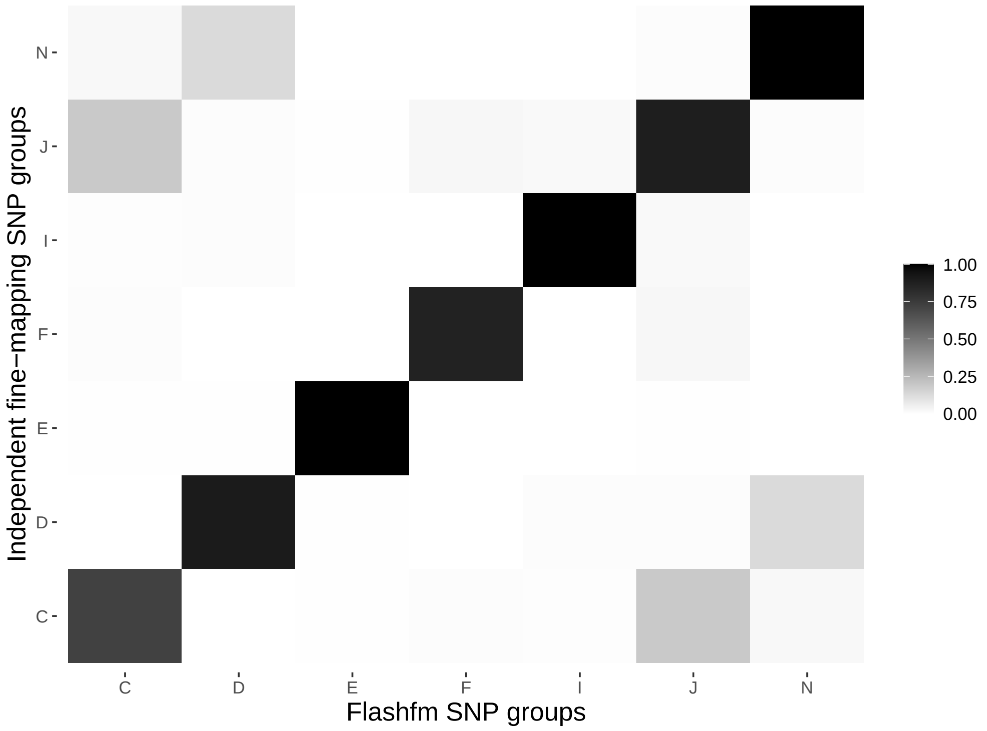

Supplementary Figure 95: Average LD between SNP groups constructed under independent fine-mapping (left) and Flashfm (right) for region 19:45359667-45397471.

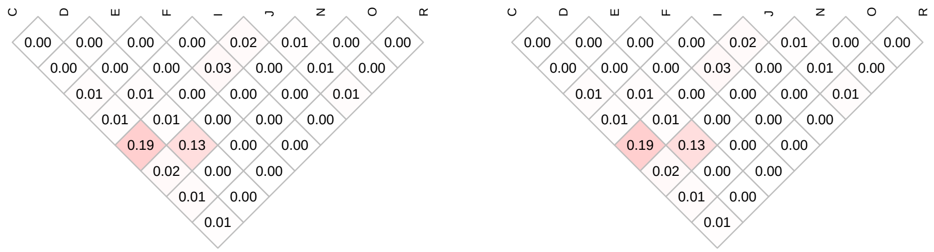

| | Stepwise | | Independent | | Flashfm | |
|---|---|---|---|---|---|---|
| Traits | SNP/Model | P-value | Model | PP | Model | PP |
| lowdlipo | 19:11197261/F | 5.54756e-17 | A+E+F | 0.347 | A+E+F | 0.338 |
| | – | – | A+F | 0.194 | E+F | 0.177 |
| | – | – | B+E+F | 0.133 | A+F | 0.17 |
| | – | – | E+F | 0.121 | B+E+F | 0.115 |
| | – | – | B+F | 0.053 | – | – |
| cholesterol | 19:11197261/F | 2.16355e-11 | E+F | 0.437 | E+F | 0.503 |
| | – | – | A+F | 0.150 | F | 0.16 |
| | – | – | A+E+F | 0.145 | A+F | 0.125 |
| | – | – | F | 0.132 | A+E+F | 0.1 |

Supplementary Table 103: Fine-mapping results for 19:10801367-11215226 by independent fine-mapping and Flashfm. Stepwise results are also given.

| | SNP Group | | | |
|---|---|---|---|---|
| | A | B | E | F |
| Independent | 24 | 10 | 13 | 13 |
| | 19:10837030 | 19:10833169 | 19:10801367 | 19:11196886 |
| Flashfm | 24 | 10 | 13 | 18 |
| | 19:10837030 | 19:10833169 | 19:10801367 | 19:11196886 |

Supplementary Table 104: SNP group sizes and representative variant for groups constructed by independent fine-mapping and by Flashfm for region 19:10801367-11215226.

Supplementary Figure 96: Average LD between SNP groups constructed under independent fine-mapping with those from flashfm for region 19:10801367-11215226.

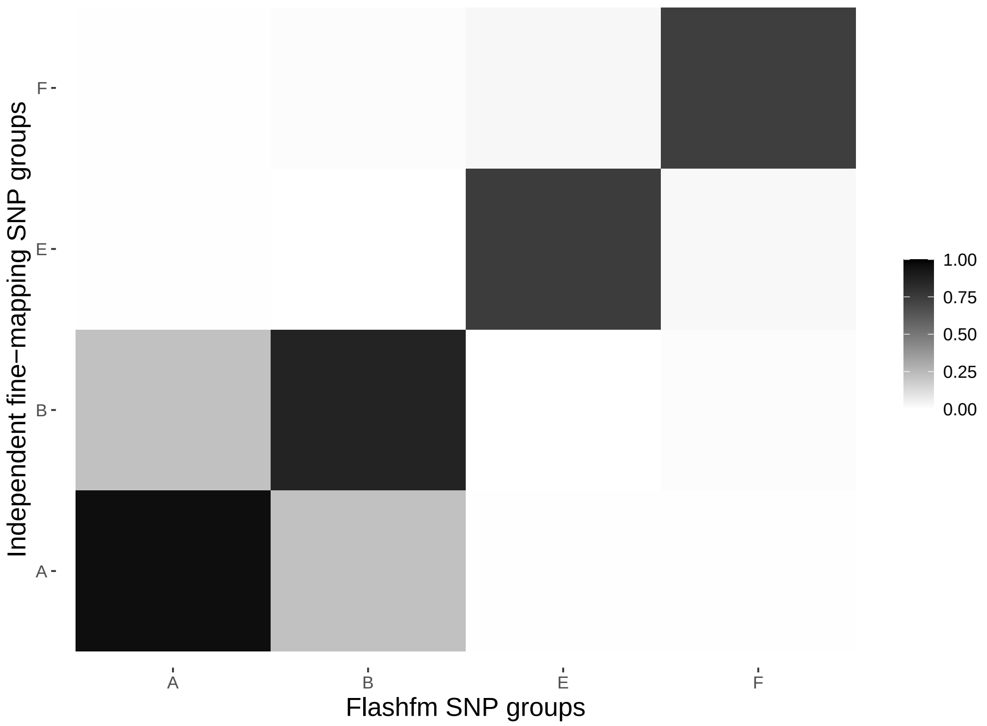

Supplementary Figure 97: Average LD between SNP groups constructed under independent fine-mapping (left) and Flashfm (right) for region 19:10801367-11215226.

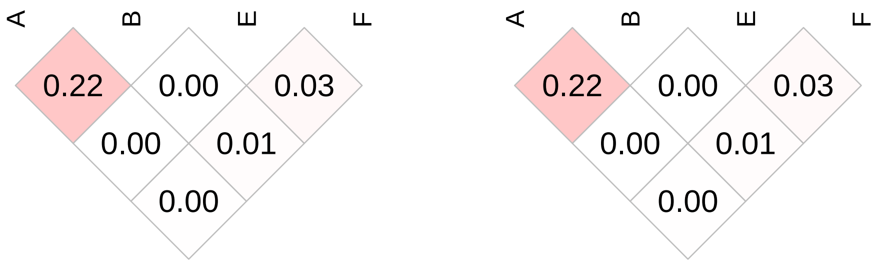

| | Stepwise | | Independent | | Flashfm | |
|---|---|---|---|---|---|---|
| Traits | SNP/Model | P-value | Model | PP | Model | PP |
| triglycerides | 19:11350488/T | 3.84117e-08 | E+G+H+I+O+P+V | 0.136 | E+G+H+I+O+P+V | 0.146 |
| | – | – | E+G+H+I+N+O+P+Y | 0.107 | E+G+H+I+O+P+Y | 0.109 |
| | – | – | E+G+H+I+O+P+Y | 0.102 | E+G+H+I+O+P | 0.096 |
| | – | – | E+G+H+I+O+P | 0.073 | E+G+H+I+N+O+P+Y | 0.091 |
| | – | – | E+G+H+I+N+O+P | 0.073 | E+G+H+I+N+O+P | 0.075 |
| | – | – | E+G+H+I+O+P+V+Y | 0.059 | A+E+G+H+I+O+P | 0.059 |
| | – | – | A+E+G+H+I+O+P | 0.056 | E+G+H+I+O+P+V+Y | 0.052 |
| | – | – | E+G+H+I+N+O+P+V | 0.051 | – | – |
| lowdlipo | 19:11368341/B | 1.31929e-07 | B | 0.538 | B+H | 0.69 |
| | – | – | – | – | B | 0.196 |
| cholesterol | 19:11370607/B | 3.14433e-07 | B | 0.53 | B+H | 0.699 |
| | – | – | B+M | 0.08 | H | 0.096 |
| | – | – | K | 0.07 | B | 0.051 |

Supplementary Table 105: Fine-mapping results for 19:11299189-11405641 by independent fine-mapping and Flashfm. Stepwise results are also given.

| | SNP Group | | | | | | |
|---|---|---|---|---|---|---|---|
| | A | B | E | G | H | I | K |
| Independent | 11 | 4 | 2 | 1 | 1 | 1 | 3 |
| | 19:11393565 | 19:11370607 | 19:11309871 | 19:11309642 | 19:11313801 | 19:11319978 | 19:11368451 |
| Flashfm | 11 | 4 | 2 | 1 | 1 | 1 | 2 |
| | 19:11393565 | 19:11370607 | 19:11309871 | 19:11309642 | 19:11313801 | 19:11319978 | 19:11366754 |
| | M | N | O | P | V | Y | |
| Independent | 11 | 14 | 1 | 1 | 4 | 3 | |
| | 19:11338881 | 19:11300862 | 19:11316315 | 19:11316547 | 19:11310834 | 19:11379717 | |
| Flashfm | 0 | 13 | 1 | 1 | 4 | 3 | |
| | – | 19:11300862 | 19:11316315 | 19:11316547 | 19:11310834 | 19:11379717 | |

Supplementary Table 106: SNP group sizes and representative variant for groups constructed by independent fine-mapping and by Flashfm for region 19:11299189-11405641.

Supplementary Figure 98: Average LD between SNP groups constructed under independent fine-mapping with those from flashfm for region 19:11299189-11405641.

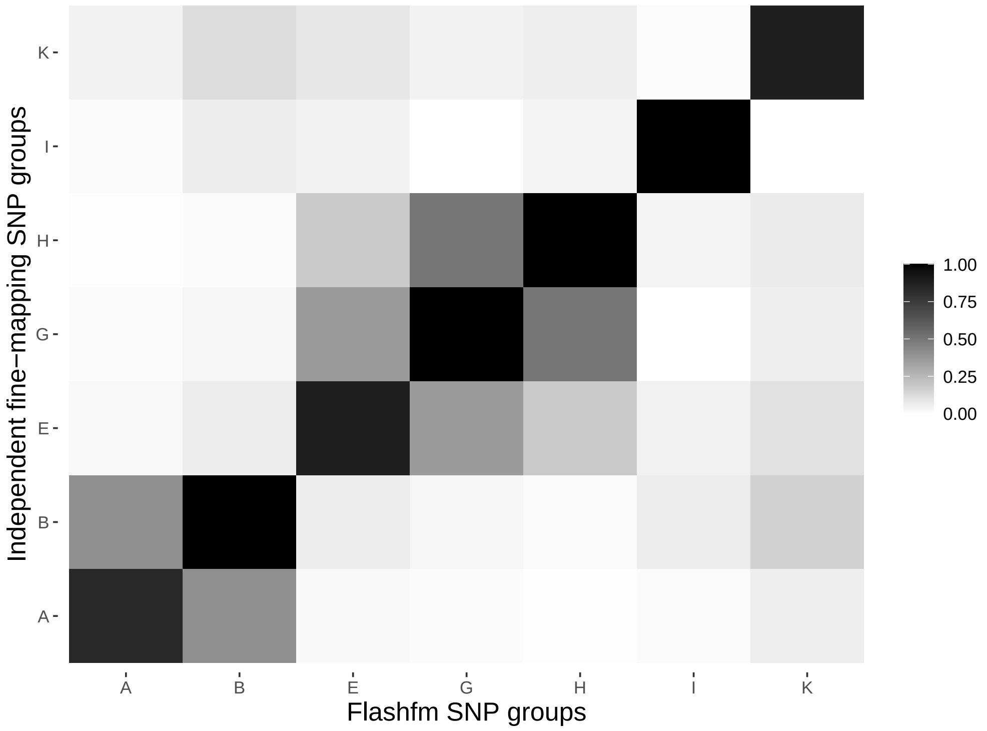

Supplementary Figure 99: Average LD between SNP groups constructed under independent fine-mapping (left) and Flashfm (right) for region 19:11299189-11405641.

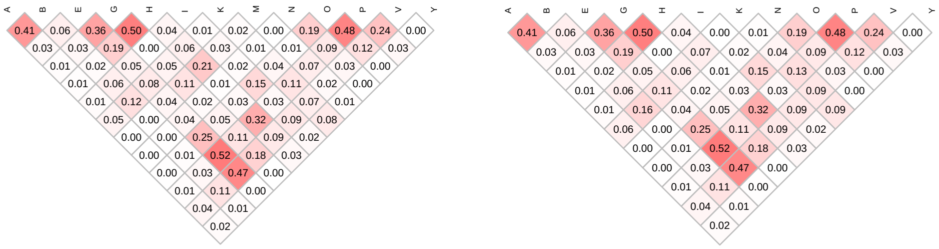

| | Stepwise | | Independent | | Flashfm | |
|---|---|---|---|---|---|---|
| Traits | SNP/Model | P-value | Model | PP | Model | PP |
| lowdlipo | 19:11281332/G | 5.65495e-08 | G | 0.570 | G | 0.679 |
| | – | – | C+G | 0.127 | C+G | 0.086 |
| cholesterol | 19:11281332/G | 2.11929e-07 | G | 0.735 | G | 0.811 |

Supplementary Table 107: Fine-mapping results for 19:11277607-11283222 by independent fine-mapping and Flashfm. Stepwise results are also given.

| | SNP Group | |
|---|---|---|
| | C | G |
| Independent | 2 | 1 |
| | 19:11279456 | 19:11281332 |
| Flashfm | 2 | 1 |
| | 19:11279456 | 19:11281332 |

Supplementary Table 108: SNP group sizes and representative variant for groups constructed by independent fine-mapping and by Flashfm for region 19:11277607-11283222.

Supplementary Figure 100: Average LD between SNP groups constructed under independent fine-mapping with those from flashfm for region 19:11277607-11283222.

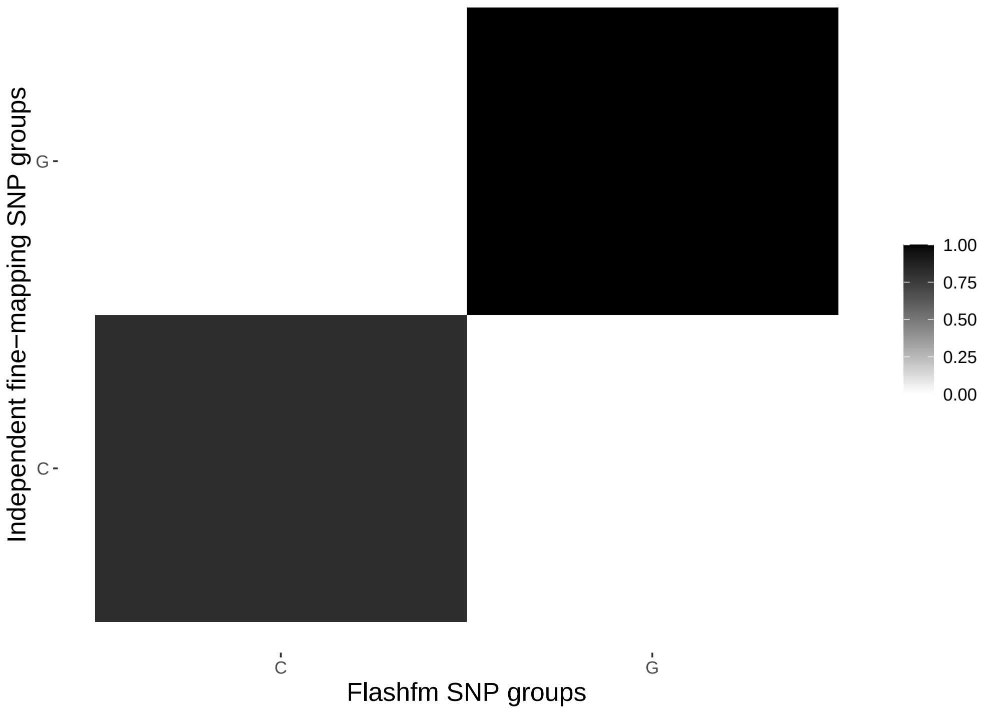

Supplementary Figure 101: Average LD between SNP groups constructed under independent fine-mapping (left) and Flashfm (right) for region 19:11277607-11283222.

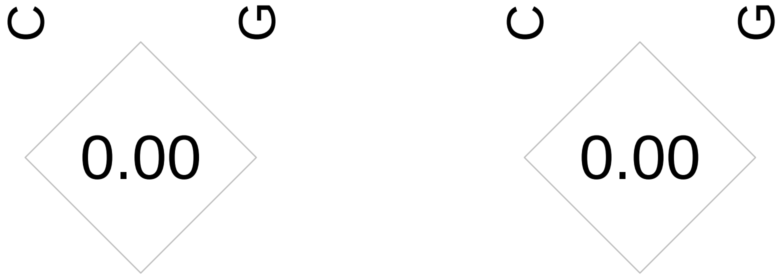

| | Stepwise | | Independent | | Flashfm | |
|---|---|---|---|---|---|---|
| Traits | SNP/Model | P-value | Model | PP | Model | PP |
| highdlipo | 20:22468476/H | 2.67091e-08 | H | 0.224 | F.1+H | 0.321 |
| | – | – | C+H | 0.21 | H | 0.204 |
| | – | – | F+H | 0.197 | C+H | 0.134 |
| | – | – | G | 0.057 | C+F.1+H | 0.062 |
| cholesterol | 20:22450576/F | 3.05885e-07 | F | 0.743 | F.1 | 0.782 |
| | – | – | – | – | F.1+H | 0.057 |

Supplementary Table 109: Fine-mapping results for 20:22424162-22486585 by independent fine-mapping and Flashfm. Stepwise results are also given.

| | SNP Group | | | | |
|---|---|---|---|---|---|
| | C | F | G | H | F.1 |
| Independent | 21 | 48 | 7 | 1 | 0 |
| | 20:22452490 | 20:22476028 | 20:22482193 | 20:22468476 | – |
| Flashfm | 21 | 0 | 7 | 1 | 18 |
| | 20:22452490 | – | 20:22482193 | 20:22468476 | 20:22450576 |

Supplementary Table 110: SNP group sizes and representative variant for groups constructed by independent fine-mapping and by Flashfm for region 20:22424162-22486585.

Supplementary Figure 102: Average LD between SNP groups constructed under independent fine-mapping with those from flashfm for region 20:22424162-22486585.

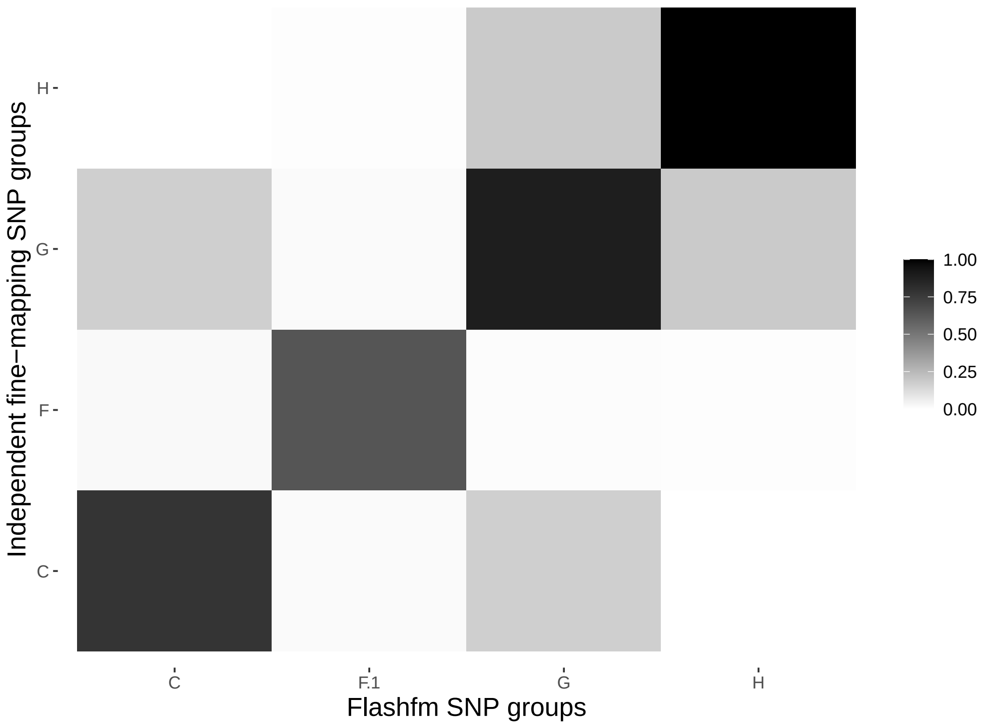

Supplementary Figure 103: Average LD between SNP groups constructed under independent fine-mapping (left) and Flashfm (right) for region 20:22424162-22486585.

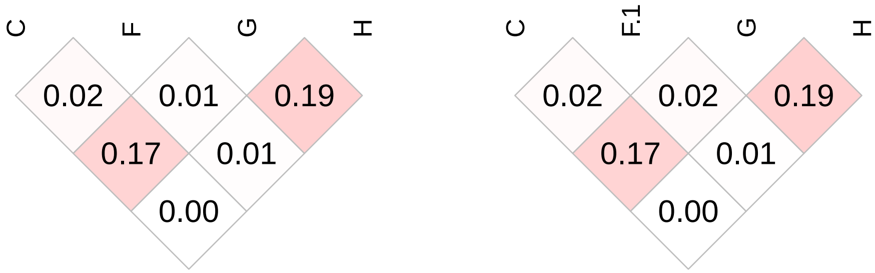

| | Stepwise | | Independent | | Flashfm | |
|---|---|---|---|---|---|---|
| Traits | SNP/Model | P-value | Model | PP | Model | PP |
| hgb | 22:34741507/B | 5.78131e-08 | B | 0.894 | B | 0.921 |
| rbc | 22:34724587/A | 7.44263e-08 | A | 0.336 | A | 0.747 |
| | – | – | B | 0.335 | – | – |
| | – | – | B+E | 0.064 | – | – |
| hct | 22:34741507/B | 1.1751e-07 | B | 0.883 | B | 0.816 |
| | – | – | – | – | A | 0.098 |

Supplementary Table 111: Fine-mapping results for 22:34644471-34814543 by independent fine-mapping and Flashfm. Stepwise results are also given.

| | SNP Group | | |
|---|---|---|---|
| | A | B | E |
| Independent | 3 | 47 | 4 |
| | 22:34706503 | 22:34792119 | 22:34672865 |
| Flashfm | 2 | 27 | 0 |
| | 22:34724587 | 22:34746385 | – |

Supplementary Table 112: SNP group sizes and representative variant for groups constructed by independent fine-mapping and by Flashfm for region 22:34644471-34814543.

Supplementary Figure 104: Average LD between SNP groups constructed under independent fine-mapping with those from flashfm for region 22:34644471-34814543.

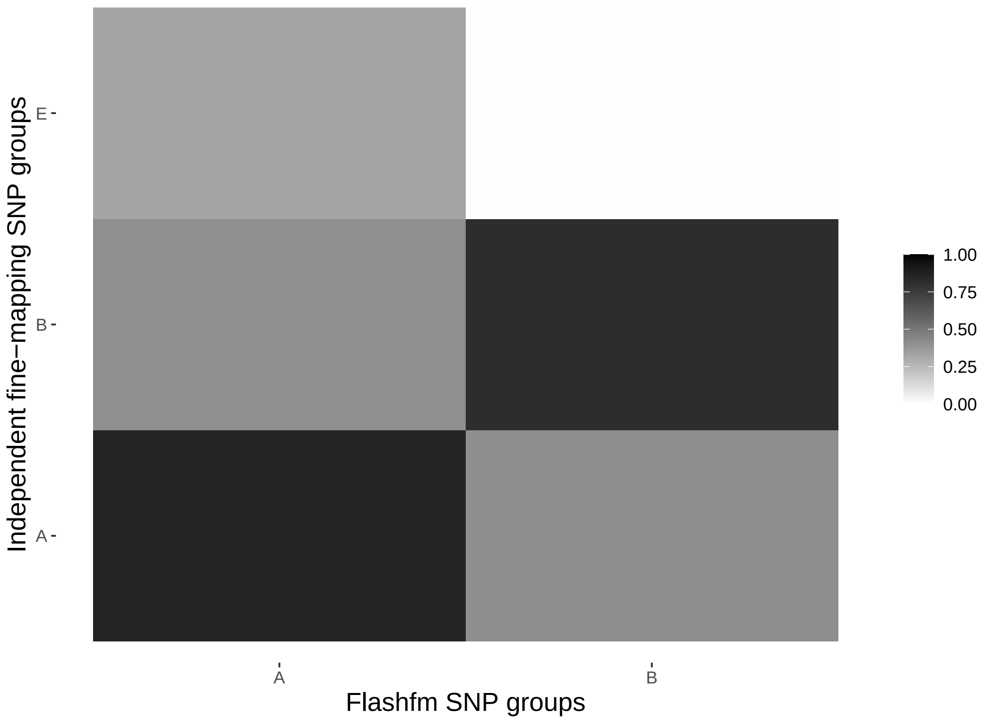

Supplementary Figure 105: Average LD between SNP groups constructed under independent fine-mapping (left) and Flashfm (right) for region 22:34644471-34814543.

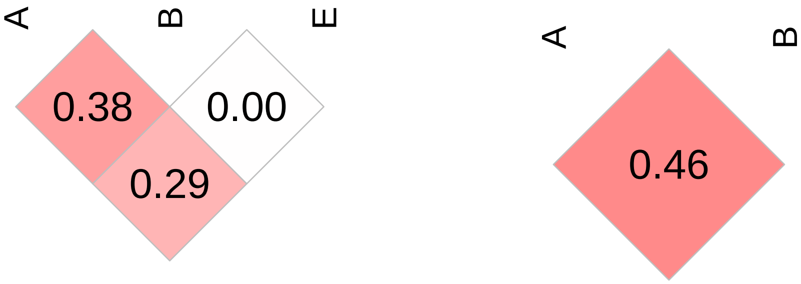
